# Engineering nanoparticle surface chemistry for antigen-presenting cell targeting improves specificity and safety of TLR3 agonist cancer immunotherapy

**DOI:** 10.64898/2026.06.23.733291

**Authors:** Victoria F. Gomerdinger, Cassandra Parada, Andrea Li, Aidan Kindopp, Justin A. Kaskow, Eva Cai, Julia B. Treese, Ivan S. Pires, Apoorv Shanker, Gil Covarrubias, Alexander D. Stoneman, Magalie Boucher, Paula T. Hammond

## Abstract

Innate immune agonists are promising therapeutic agents to induce immune responses against cancer. However, these agents have been limited by toxicity associated with systemic accumulation and activity in off-target cells. In this work, a targeted nanoparticle (NP) platform to encapsulate and protect the Toll-like receptor 3 (TLR3) agonist polyinosinic-polycytidylic acid (poly(I:C)) and promote its specific delivery to antigen presenting cells (APCs), macrophages and dendritic cells, for activation of this cell population was designed. To determine NP physiochemical properties that promote APC delivery, we developed a library of NP surface chemistries formed by electrostatic adsorption of polyanion coatings onto liposomes using layer-by-layer (LbL) assembly and screened the particles on APCs and off-target cells. Dextran sulfate was identified as a promising coating to enhance specific APC delivery. We applied these design parameters to develop a poly(I:C)-loaded NP for an APC-targeted immunotherapy. In a model of metastatic ovarian cancer, the LbL NP prolonged poly(I:C) retention in the peritoneal space—with 2-fold remaining 24-48hr after administration compared to free poly(I:C)—ultimately reducing systemic accumulation and associated toxicities. Compared to free drug, the NP reduced the increase in serum levels of TNFα, IL-6, and CXCL10 by 9-, 4-, and 31-fold respectively. NP-treated mice experienced lower weight loss and recovered more quickly at a higher poly(I:C) dose, indicating a widening of the therapeutic window. The NP formulation enhanced accumulation of poly(I:C) in the tumor 2-fold and activation of the target APC population compared to free drug, and ultimately slowed tumor growth and extended survival in combination with doxorubicin chemotherapy. Overall, this work demonstrates a modular NP delivery strategy to improve the delivery, safety, and therapeutic window of a TLR3 agonist.

## Introduction

High-grade serous ovarian cancer (HGSOC), also known as high grade serous carcinoma (HGSC) accounts for the majority of epithelial ovarian cancer diagnoses and ∼70% of gynecologic cancer deaths^1, 2^. It is an aggressive cancer that predominantly presents at stage III or IV^3, 4^ (∼80%), with widespread peritoneal and abdominal metastases^5^. Although standard treatment with cytoreductive surgery and platinum/taxane chemotherapy is initially effective^6^, over 80% of patients develop recurrent, platinum-resistant disease^7^. Immunotherapy to stimulate the patient’s immune system to attack cancer has shown great promise in generating robust, long-lived responses which have significantly improved outcomes for a multitude of cancers^8–10^. Unfortunately, HGSC has shown limited responsiveness to immune checkpoint inhibitors, underscoring the need for alternative immunotherapy strategies^11^.

Targeting innate immune cells in the tumor environment—particularly antigen-presenting cells (APCs) such as dendritic cells and macrophages—can enhance T cell priming^12^ and reprogram tumor-associated macrophages toward a pro-inflammatory, anti-tumor phenotype^13, 14^. Strategies to improve the activation and maturation of APCs include agonists of pattern-recognition receptors (PRRs) expressed by these cells^15–17^. While immune agonists are promising therapies, many experience dose-limiting toxicities due to PRR activation throughout the body which induces systemic cytokine release that leads to severe health effects^18–20^. Additionally, accumulation of agonists in non-immune cells such as fibroblasts^21, 22^, epithelial cells^23^, hepatocytes^24^, and endothelial cells^25–27^ elevate pro-inflammatory cytokine production in these cell populations, leading to additional adverse effects^28^. These immune-related adverse events have limited the use of these therapeutics. Various strategies to improve the stability, pharmacokinetics, and safety profile of immunomodulatory agents include micro– and nanocarriers^18, 29–32^, pro-drugs^33^, and local administration routes such as intratumoral and subcutaneous^32, 34, 35^ to retain the therapeutic at the tumor or administration site and to prevent burst systemic exposure. While these strategies are promising, there has been limited translation to immunologically cold, disperse and highly metastatic cancers such as HGSC.

As ovarian cancer is predominantly metastatically disseminated throughout the peritoneal cavity by the time of diagnosis^6^, intraperitoneal (IP) administration is a viable strategy to improve therapeutic accumulation in the tumor. IP administration can increase drug penetration due to higher local drug concentrations and slower drug clearance compared to intravenous administration^14, 36, 37^. However, intraperitoneally administered agents eventually enter systemic circulation^38^, so a delivery strategy that can promote prolonged retention in the peritoneal space and direct therapeutics to the cells of interest while mitigating accumulation in systemic circulation and off-target cell populations is desired to improve therapeutic efficacy and reduce the toxicity of immune agonists.

Layer-by-Layer (LbL) NPs, which are assembled by electrostatic adsorption of oppositely charged polymers onto a charged NP substrate, offer a strategy to develop targeted NPs to improve the safe delivery of potent therapeutics to APCs. The LbL NP system provides a modular platform to facilitate the encapsulation of a wide range of therapeutics—such as proteins^39^, small-molecules^40, 41^, and nucleic acids^41–43^. Additionally, the optimization of NP physiochemical properties enables the NPs to direct the targeting of the cargo to organs and cell populations as well as to modulate the pharmacokinetics via the choice of NP surface chemistry^43–45^ and core stiffness^46^. LbL NPs have also been previously demonstrated to reduce the toxicity of immunomodulatory agents^39^.

In this work, we aimed to develop an APC-directed nanoparticle (NP) formulation by optimizing the NP surface chemistry and core stiffness to facilitate selective and internal delivery to APCs, and to subsequently use these design parameters to demonstrate the targeted delivery of an immune agonist with improved safety. A range of strategies have been employed to target APCs, including the use of carbohydrates, antibodies, and other moieties to target receptors expressed on these cells^47–52^—such as C-type lectin receptors^53, 54^ and specific cell-surface antigens (e.g., CD11c, CD80/CD86, CD40)^54–56^. Leveraging the modular LbL system, we screened a library of anionic polypeptides and polysaccharides to determine whether surface chemistry composition could direct APC-selective association. We identified dextran sulfate (DXS) as a promising candidate to enhance accumulation in APCs while minimizing interactions with off-target cell types. Additionally, we demonstrated the *in vivo* targeting of DXS-coated NPs to tumor-associated APCs.

We then used these NP design parameters to develop polyinosinic-polycytidylic acid (poly(I:C))-loaded NPs to demonstrate safe, targeted delivery of an immune agonist. Poly(I:C), a double-stranded RNA analog and Toll-like receptor 3 (TLR3) and retinoic acid-inducible gene I (RIG-I) agonist^17, 57, 58^, was chosen as an example immune agonist as it promotes anti-tumor immunity^13, 17, 57^ but has been limited in the clinic by toxicity. Notably, TLR3 and RIG-I are expressed on dendritic cells (DCs) and macrophages, as well as non-immune cells^16, 59–62^, motivating a strategy to reduce off-target delivery. Various strategies have been employed to stabilize poly(I:C) from degradation^20^ and reduce systemic exposure, including complexation with polymers^31, 63–66^ and in lipid nanoparticles (LNPs)^67, 68^. However, many formulations are non-targeted and subsequently leak into systemic circulation, thereby often requiring local administration such as intratumoral^67, 68^ or subcutaneous^15, 31^ routes which are not as relevant for treating ovarian cancer.

Upon intraperitoneal administration, LbL encapsulation of poly(I:C) improved its pharmacokinetic profile and reduced systemic toxicity compared to free drug, as evidenced by lower levels of circulating inflammatory cytokines and reduced treatment-associated weight loss. The efficacy of the Poly(I:C) LbL NP in combination with the chemotherapeutic doxorubicin (Dox) was evaluated in a murine model of ovarian cancer. Dox was selected for combination therapy due to its clinical use as a second-line treatment for ovarian cancer and its role in immunogenic cell death. The additional source of adjuvanticity and generation of antigens in the tumor environment induced by Dox was hypothesized to synergize with the immunomodulation induced by poly(I:C)^69^. Compared to Dox alone, the addition of Poly(I:C) NP to Dox further reduced the growth of tumor and more substantially improved survival in the BPPNM metastatic model of ovarian cancer^70^, highlighting this targeted NP strategy as a viable option to improve the delivery of immune agonists for ovarian cancer.

## Results and Discussion

### Dextran sulfate and fucoidan surface chemistries selectively enhance NP association with APCs

To engineer a NP formulation that can promote more selective trafficking of NPs by APCs, the effects of NP surface chemistry on NP trafficking were evaluated using the modular LbL NP system (**Fig. 1a**). We first sought to screen a larger library of outer polymer chemistries to identify LbL surface chemistries that selectively enhance NP association with APCs. LbL NPs formulated with 10 different anionic polymers serving as the outermost NP layer were evaluated for their association with APCs, ovarian cancer cells, and other off-target cells (fibroblasts, epithelial cells, hepatocytes, and endothelial cells).

**Fig. 1.**
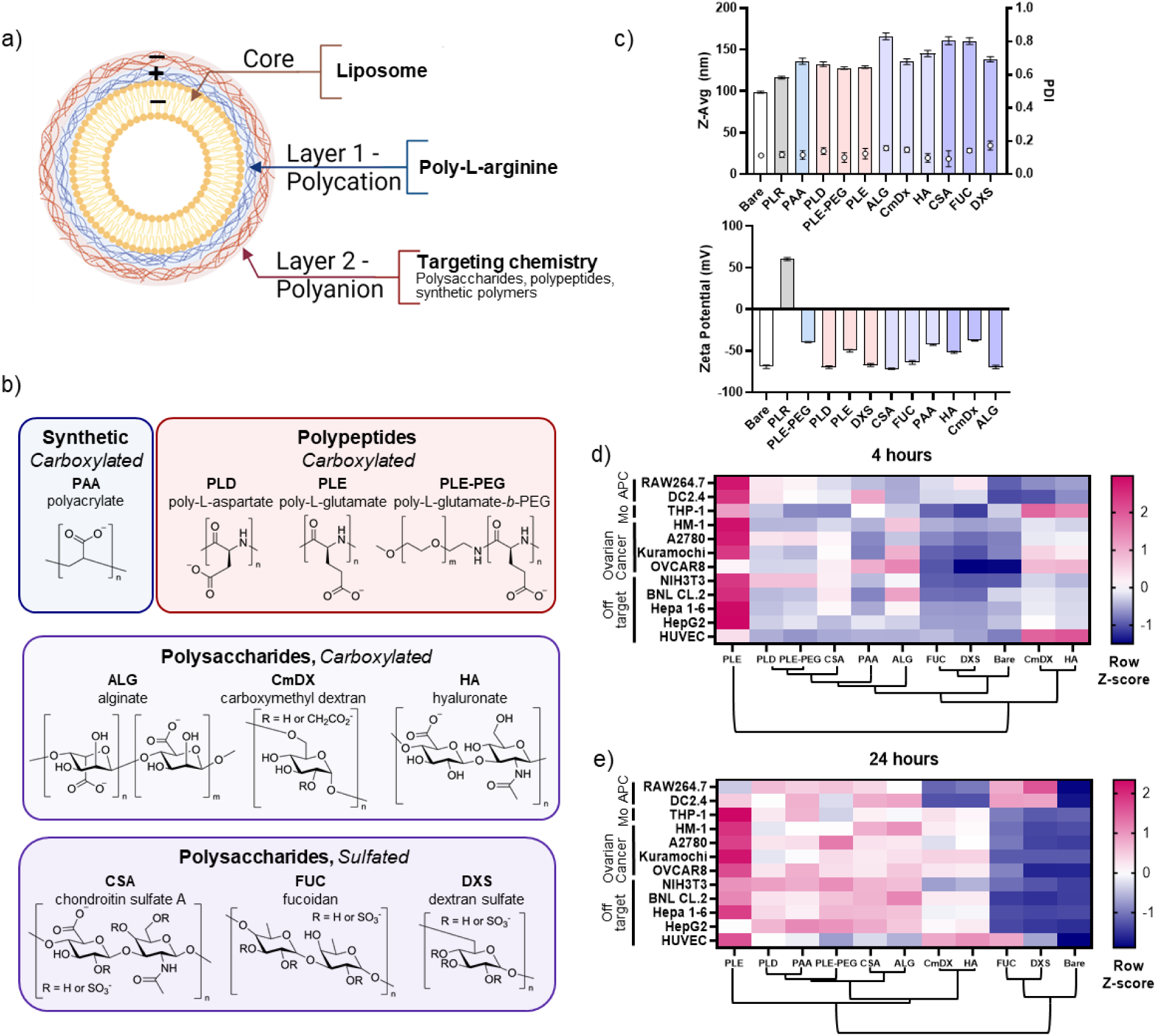
LbL NPs for intracellular and selective targeting of antigen presenting cells. a) Schematic of layer-by-layer NP, with anionic liposome core adsorbed with the polycation poly-L-arginine followed by a polyanion to serve as the targeting chemistry of the NP. b) NP outer layer library consisting of synthetic polymers, polypeptides, and polysaccharides with carboxylated and sulfated surface chemistries. c) Size, polydispersity index (PDI) and b) zeta potential (ZP) of liposomes unlayered (bare) or layered with poly-L-arginine followed by various polyanions as the terminal layer. Associated NP fluorescence for various cell lines after d) 4 hr and e) 24 hr incubation with 25 μg/mL fluorescently labeled NPs with varying surface chemistry. NP association was quantified using flow cytometry and normalized by assessing the fold change of median fluorescence intensity of each NP formulation over that of the bare liposome. Row Z-scores for each cell line were plotted in d-e, with hierarchical cluster analysis by surface chemistry displayed in the dendrograms below each heatmap. N=2-3 biological replicates per cell line, each with 3 technical replicates. APC: antigen presenting cell; Mo: monocyte.

For the outer layer NP library, we focused on negatively-charged polymers to serve as the LbL NP surface chemistry as anionic NPs demonstrate superior biocompatibility, reduced non-specific cellular uptake, and lower clearance rates *in vivo* compared to cationic NPs^71–73^. The library of polymers included sulfated and carboxylated chemistries and consisted of polysaccharides, homopolypeptides, a PEGylated polypeptide block copolymer, and a synthetic polymer (**Fig. 1b**). Polysaccharides were included as a dominant fraction of the library as APCs express receptors that bind carbohydrate moieties, facilitate internalization, and have been used for APC-targeting^74–77^. The homopolypeptides poly-L-aspartate (PLD) and poly-L-glutamate (PLE) have previously been demonstrated to enhance targeting with a number of cancer cells via association and clustering of specific overexpressed amino acid transporters^39, 43–45^. Poly(ethylene glycol)-b-poly-L-glutamate (PLE-PEG) and polyacrylic acid (PAA) were included as potential inert controls as PEGylation^14^ and PAA^78^ have been demonstrated to reduce NP interactions with myeloid cells. The molecular weight of all polymers was maintained in the 10-20 kDa range such that molecular weight was not a variable. Anionic liposomes with 34 mol% cholesterol were utilized as the NP core (referred to as ‘Compliant’ liposomes **Table S1**). Dye-labeled liposomes were layered with the polycation poly-L-arginine (PLR) followed by the various polyanions. The size and zeta potential of the library of LbL NPs exhibited similar sizes (100 to 150 nm), zeta potentials (–35 to –65 mV) and PDI (<0.2) (**Fig. 1c**).

Cells were treated with the fluorescently-labeled NPs at equivalent lipid concentrations, and at the noted timepoints, were prepared for flow cytometry to quantify the associated NP fluorescence. To facilitate comparison of the relative cellular association of the NP formulations, the associated NP fluorescence was normalized across the NP surface chemistries for each individual cell line using Z-scores, and the data was analyzed by hierarchical clustering to assess similarities in NP association across surface chemistries, with the dendrograms displayed below each heatmap (**Fig. 1d-e**, with raw data in **Fig. S1-2**). Poly-L-glutamate (PLE) had the highest association with APCs at 4hr, but also the highest association across many other cell types evaluated including fibroblasts (NIH3T3) and hepatocytes (BNL CL.2, Hepa 1-6, HepG2), making it a non-ideal surface chemistry candidate to target cargo with high specificity for APCs. Similarly, the sulfated polysaccharide chondroitin sulfate A (CSA), synthetic polymer polyacrylic acid (PAA), and polypeptide PLD also demonstrated improved association with APCs but additionally had high association with other cell types. Although PLE-PEG contains PEG, which is often used to minimize cellular interactions^14^, it generally had higher association across the cell types tested compared to the inert bare liposome core, possibly modulated by the PLE component of the block polymer or non-specific interactions of the PEG, but overall lower association compared to PLE. The polysaccharides hyaluronic acid (HA) and carboxymethyl dextran (CMDx) both exhibited poor association with APCs and high association with endothelial cells (HUVEC) and grouped similarly by hierarchical clustering analysis as shown in the dendrograms for both 4hr and 24hr. While HA has been widely used to target CD44 on macrophages, the demonstrated improvements in targeting are largely under inflammatory conditions^79–81^—which were not employed here—as high CD44 expression is exhibited on inflamed macrophages compared to anti-inflammatory/alternatively-activated (commonly referred to as M2) macrophages^82^.

Hierarchical cluster analysis revealed that the sulfated polysaccharides DXS and fucoidan (FUC) surface chemistries both similarly minimized NP association with ovarian cancer and off-target cells while maintaining high association with macrophages and dendritic cells at short (4hr) and long (24hr) timepoints. DXS conferred a slight advantage over FUC for lower association with ovarian and other off-target cells and had lower NP association with all non-APC cells except HUVECs compared to the inert bare liposome core at 4hr, and this trend largely held true at the 24hr timepoint. Both the DXS and FUC polymers are ligands of the scavenger receptor A (SR-A, CD204) which is expressed on APCs^76, 83^, and NP systems decorated with dextrans and derivatives have been used to target macrophages^84–87^. Pro-tumorigenic tumor-associated macrophages express CD204, with infiltration of CD204^+^ macrophages in tumors correlating with aggressiveness of a variety of cancers and poorer clinical outcomes^88–91^, highlighting it as a relevant target in the context of cancer.

DXS and CMDx serve as an interesting comparison for the influence of chemical functional groups as they are sulfated and carboxylated derivatives of dextran, respectively. LbL NPs with DXS and CMDx surface chemistries exhibit drastic differences in cell association, suggesting that sulfated surface chemistries may offer an advantage for enhancing NP association with APCs. DXS was selected as the APC-targeting formulation to move forward with as it had higher association with both macrophages and dendritic cells compared to FUC at both 4hr and 24hr. DXS was further validated to have low association with BPPNM ovarian cancer cells which were used for subsequent *in vivo* studies (**Fig. S3**).

Next, we characterized the *in vivo* pharmacokinetics, biodistribution, and cellular targeting of DXS LbL NPs in our ovarian cancer model of interest, BPPNM, upon intraperitoneal administration (**Fig. 2a**). BPPNM is a murine model of HGSC derived from fallopian epithelial cells, metastasizes to relevant organs, and is poorly responsive to immune checkpoint blockade. Due to its genotype (*p53^−/−R172H^Brca1^−/−^Pten^−/−^Nf1^−/−^Myc^OE^*), it is a model of homologous-repair deficient disease^70^. BPPNM-tumored mice were injected with equivalent doses (200 μg lipid) of unlayered (bare), PLD-coated—as a comparison a surface chemistry demonstrated to improve ovarian cancer cell targeting^40, 44^ (**Fig. S3**)—or DXS-coated liposomes. Liposomes were labeled with BDP-650 to enable visualization of the NPs using an In Vivo Imaging System (IVIS) and flow cytometry. PLR was used as the first polymer layer followed by PLD or DXS as the terminal layer. PLD and DXS LbL NPs exhibited slower clearance from the peritoneal space over 24hrs compared to bare liposomes (**Fig. 2b**, with raw data in **Fig. S4**). Twenty-four hours after NP injection, PLD and DXS LbL NPs had significantly higher signal in the omental tumor compared to bare liposomes, more substantially for the PLD LbL NPs, which is expected given the demonstrated ovarian cancer targeting of this surface chemistry^40, 44^, possibly due to expression of amino acid transporters which interact with PLD^92^ (**Fig. 2c-d**, with raw data in **Fig. S5**). PLD and DXS LbL NPs exhibited lower signal in the spleen, an off-target organ, compared to bare liposomes (**Fig. 2c**). To assess NP distribution throughout the mice with higher spatial resolution, cryo-fluorescence tomography (CFT) imaging of the whole mouse was performed. CFT showed that the DXS and PLD NPs were also able to target tumor metastases throughout the peritoneal cavity and had greater accumulation in the mediastinal lymph nodes which drain the peritoneal space compared to the bare liposomes (**Fig. S6**).

**Fig. 2.**
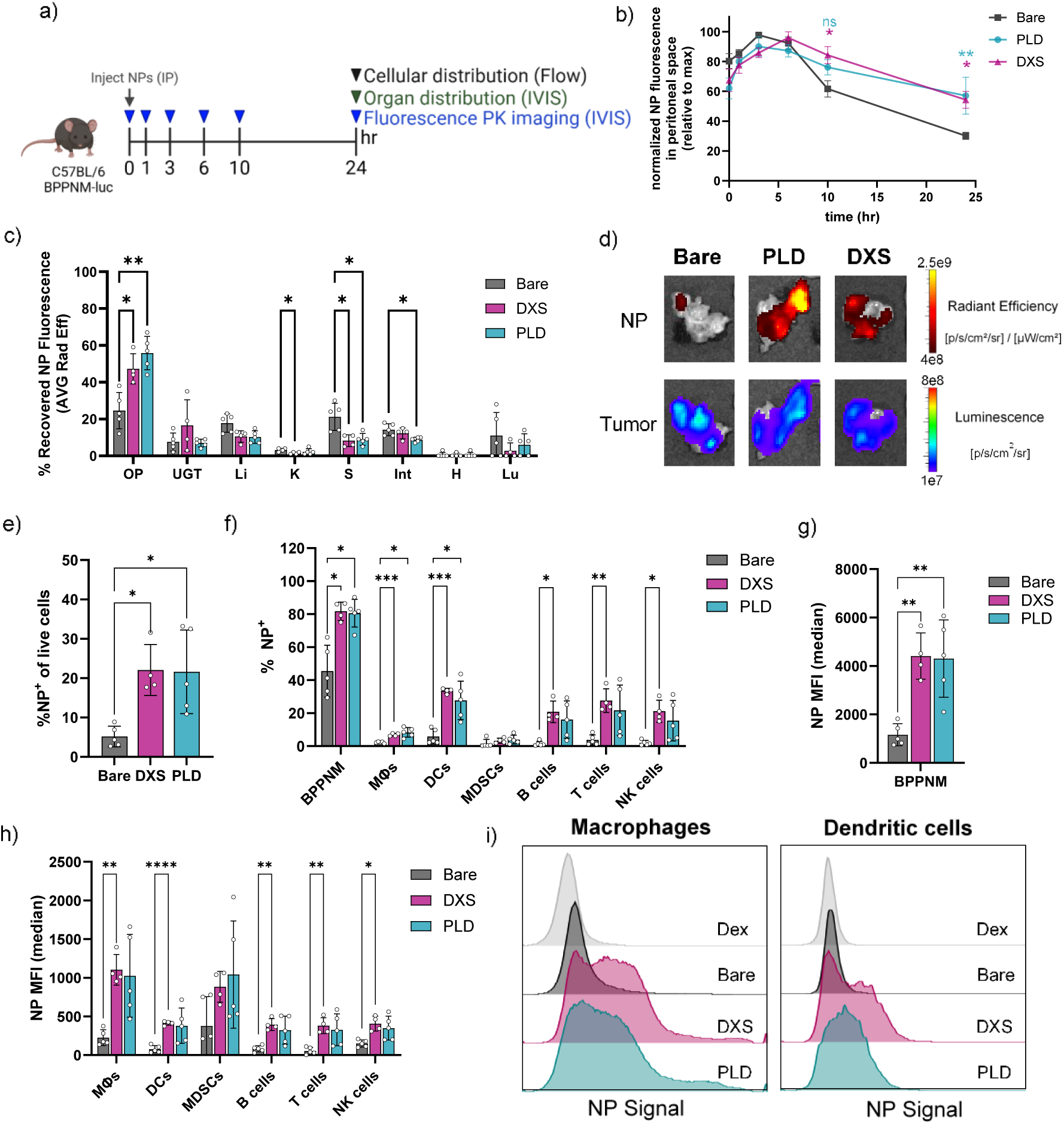
LbL NPs enhance tumor-tissue and tumor-associated immune cell accumulation. a) Female C57BL/6 mice with BPPNM-luc tumors (3 million, IP) were injected with 200 μg of BDP650-tagged bare liposomes, PLD LbL NPs, or DXS LbL NPs in 5% dextrose 10 days post tumor inoculation (N=4-5). Mice were imaged using an in vivo imaging system (IVIS) for NP fluorescence at various timepoints. b) NP radiant efficiency in the peritoneal space over time, normalized to the maximum NP signal for each mouse (mean±SEM). Organs were removed 24 hrs after NP injection and imaged for NP fluorescence and tumor bioluminescence *ex vivo* using IVIS. Mice injected with 5% dextrose (vehicle control) were used to subtract background organ signal. c) Percent of recovered NP fluorescence (average radiant efficiency, [p/s/cm²/sr]/ [µW/cm²]) in organs (mean±SD). d) Representative images of NP fluorescence and tumor bioluminescence in omental tumors. Ascites and tumors were collected 24 hrs after NP injection, digested, and subjected to flow cytometry analysis to assess cellular NP distribution. Gating strategy described in Fig. S8. e) Percentage of live cells in the tumor that are positive for NP signal. f) Percentage of each cell type in the tumor that are positive for NP signal. NP Median-Fluorescence Intensity (MFI) in g) BPPNM cells and h) tumor-associated immune cells. i) Representative histograms of NP signal in macrophages and dendritic cells. Statistically significant comparisons determined using 2-way ANOVA in c, f, h) and 1-way ANOVA in e, g) with Tukey multiple comparisons test. PLD – poly-L-aspartate, DXS – dextran sulfate, MΦs – macrophages, DCs – dendritic cells, MDSCs – myeloid-derived suppressor cells, NK – natural killer. (*p < 0.05, **p < 0.01, ***p < 0.001, ****p < 0.0001).

Twenty-four hours after NP injection, ascites and tumors were digested to isolate single cells and quantify cellular NP association using flow cytometry. Compared to bare liposomes, PLD and DXS LbL NPs had a significantly higher percentage of cells in the tumor that were NP^+^ (**Fig. 2e**). Macrophages were the most prevalent population in the NP^+^ cells in the tumor and ascites (**Fig. S7a-b, with gating strategy in Fig. S8**). PLD and DXS LbL NPs similarly resulted in a higher percentage of BPPNM cells positive for NPs and higher overall NP signal in this cell population in the tumor compared to bare liposomes (**Fig. 2f-g**). For the populations of tumor-associated immune cells profiled—macrophages, DCs, myeloid-derived suppressor cells (MDSCs), B cells, T cells, and natural killer (NK) cells—PLD and DXS LbL NPs resulted in similar percentages of cells that were NP^+^ as well as the average NP MFI; however, the DXS LbL NPs demonstrated lower variability in NP association compared to PLD, resulting in a significant improvement in net NP accumulation in macrophages, DCs, B cells, T cells, and NK cell compared to bare liposomes (**Fig. 2f, h, i**). Similar trends were observed in the ascites cells (**Fig. S7d, f**). This suggests that the DXS surface chemistry might be more consistent in improving NP delivery to these immune cell populations compared to PLD. High DXS NP association with T and B cells was unexpected as these cell populations do not express the scavenger receptor, SR-A^93^. To probe this, we assessed the *ex vivo* association of NPs with T and B cells isolated from the spleen and found that DXS NPs had lower association compared to PLD NPs in this setting (**Fig. S9**). The elevated *in vivo* NP association of DXS NPs with T and B cells could be due to more DXS NPs being bioavailable compared to PLD NPs as DXS NPs interact with fewer cell types based on earlier data, or possibly due to differences in protein coronas that may affect NP association^14, 94^. These results highlight a limitation of *in vitro* screening, so additional studies should be conducted to probe these differences.

### Poly(I:C) can be stably loaded into the LbL NP film for targeted delivery

We leveraged prior work on the incorporation of small nucleic acid cargos such as siRNA into LbL films adsorbed onto NPs^95–98^ to develop a method to load low-molecular weight poly(I:C) (200-1000 base pairs), a much larger nucleic acid. Moreover, unlike prior systems that electrostatically adsorbed poly(I:C) onto the surface of micro– and nano-particles as the terminal layer^99–102^, we engineered a tetralayer LbL NP with complexing polycations as first and third layers encapsulating layers, poly(I:C) embedded as the second layer, and DXS as the fourth layer to serve as the NP surface chemistry (**Fig. 3a**). This nanocarrier design aims to improve the stability of poly(I:C) in the film through the adsorption of subsequent layers as well as to improve the targeting of the therapeutic to APCs by using DXS as the surface chemistry.

**Fig. 3.**
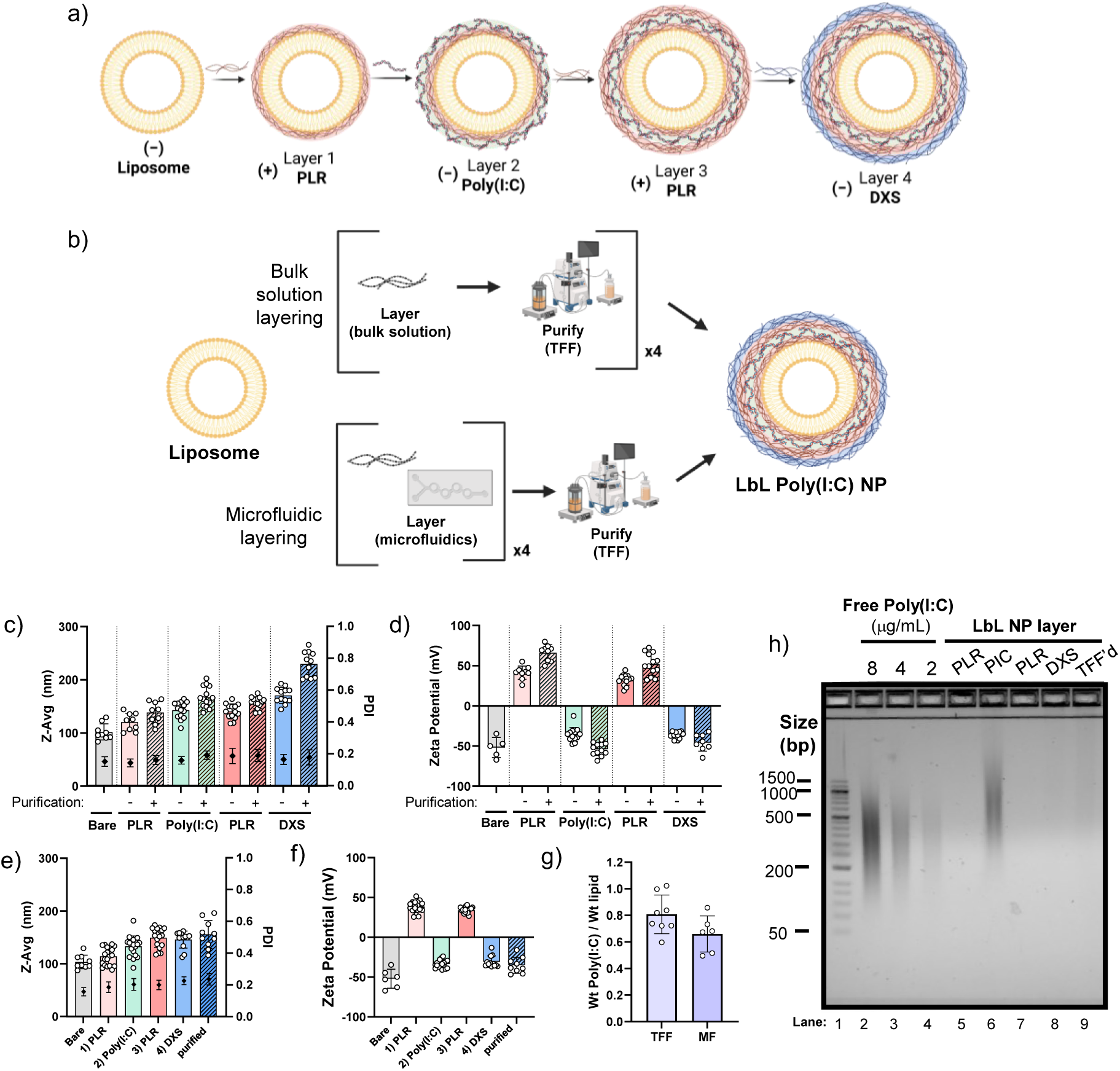
LbL NP assembly via two methods stably encapsulates poly(I:C) on NP. a) Schematic of tetralayer LbL NP system for targeted delivery of poly(I:C). Liposomes were layered with poly-L-arginine (PLR), then poly(I:C) low-molecular weight, PLR, and dextran sulfate (DXS). b) Schematic of LbL assembly using bulk layering with tangential flow filtration (TFF) purification in between each layer or microfluidic layering without purification between layering steps and final purification using TFF. c) Size, polydispersity index (PDI) (mean±SD) and d) zeta potential (ZP) of LbL Poly(I:C) NPs assembled using bulk layering+TFF purification. Data are shown as mean±standard deviation of 4-14 batches of NPs. e) Size, PDI and f) ZP of LbL Poly(I:C) NPs assembled using microfluidics. Data are shown as mean±standard deviation of 4-18 batches of NPs. g) Loading of poly(I:C) on NPs, represented as the weight ratio of poly(I:C) to lipid on the final NP construct. Data are shown as mean±standard deviation of 6-8 batches of NPs. h) Nucleic acid agarose gel of free poly(I:C) at varying concentrations and LbL-Poly(I:C) NP after assembly of each layer. NP samples were concentration matched to 4 μg/mL poly(I:C), or equivalent liposome concentration for the first PLR layer.

We demonstrate the synthesis of the LbL poly(I:C) NP using two methods: bulk layering coupled with tangential flow filtration (TFF) for purification after each layer as well as a microfluidic-based layering approach (**Fig. 3b**). TFF purification and microfluidics have been applied previously to simpler LbL NP systems^103,104^, but the incorporation of poly(I:C) required additional optimization due to its substantially larger molecular weight. For the bulk layering approach, the weight ratio of each polymer that enables sufficient charge conversion and stability was optimized, and NPs had to be maintained at a sufficiently low concentration to reduce NP bridging during the layering and TFF purification steps. (**Fig. S10; Table S6**). For microfluidic assembly, since there is no purification step between the addition of each layer, we aimed to layer near the charge conversion point which requires lower weight equivalents of polymer compared to the TFF method, consistent with previous observations with LbL microfluidic assembly (**Table S7**)^104^. After the final layer, these microfluidic-assembled NPs were subsequently purified to remove salt and concentrated using TFF. NPs formed using both methods of assembly exhibit stable size and charge conversion with the addition of each layer (**Fig. 3c-d**), while the size of the NPs is maintained smaller with microfluidic assembly (**Fig. 3e-f**). The microfluidic-assembled NPs had a marginally lower weight loading of poly(I:C) due to the lower weight equivalents of polymer adsorbed with this method (**Fig. 3g**).

To verify that poly(I:C) is stably encapsulated in the NP, gel electrophoresis was used to assess the migration of poly(I:C) from the NP. For the bilayer NP with poly(I:C) as the outermost layer, poly(I:C) freely migrated on the gel, indicating weak encapsulation. With the addition of the subsequent PLR and DXS layers to cap the NP, poly(I:C) migration of the gel was inhibited indicating stable encapsulation in the NP (**Fig. 3h, with uncropped gel in Fig. S11**).

### Poly(I:C) LbL NPs exhibit superior activation of macrophages and DCs *in vitro* compared to free drug

Poly(I:C) induces the maturation of APCs resulting in the upregulation of costimulatory and major histocompatibility complex (MHC) molecules—which are critical for T cell priming—and also promotes cross-presentation^17^, bridging the innate and adaptive immune systems to generate productive anti-tumor immune responses. In order to assess the delivery and functional activity of the LbL Poly(I:C) NPs, RAW264.7 macrophages and bone-marrow derived dendritic cells (BMDCs) were treated with free poly(I:C), poly(I:C) NPs, control NPs (PLR/DXS bilayer NPs without poly(I:C)), or bare liposomes for 24hr. Supernatant was assessed for inflammatory cytokines via ELISA, and poly(I:C) delivery and markers of cell activation (CD86, MHC-II) were assessed using flow cytometry. Of note, free soluble poly(I:C) can be internalized by cells via SR-A^105^, the same receptor that interacts with DXS.

LbL Poly(I:C) NPs improved poly(I:C) delivery to macrophages and DCs (**Fig. 4a, S12a**) as well as activation and maturation of these cells exemplified by increasing TNF-α cytokine secretion (**Fig. 4b**) and CD86 expression (**Fig. 4c, S12b, S12e**) compared to free poly(I:C) in a dose-dependent manner. Additionally, expression of MHC-II, a marker of the ability to present antigens and an additional sign of DC maturation, was upregulated in BMDCs (**Fig. 4d, Fig. S12f**). A positive correlation of poly(I:C) delivery and CD86 expression in BMDCs was observed, with more efficient activation achieved with the LbL NP compared to free poly(I:C) at the same 10 μg/mL dose, as exemplified in a representative flow plot of poly(I:C)-Cy5 and CD86 signal and corresponding histograms at this dose (**Fig. 4e**). These trends were reproducible across independent batches of poly(I:C) NPs made by three different researchers and biologically independent cultures of BMDCs (**Fig. 4f**).

**Fig. 4.**
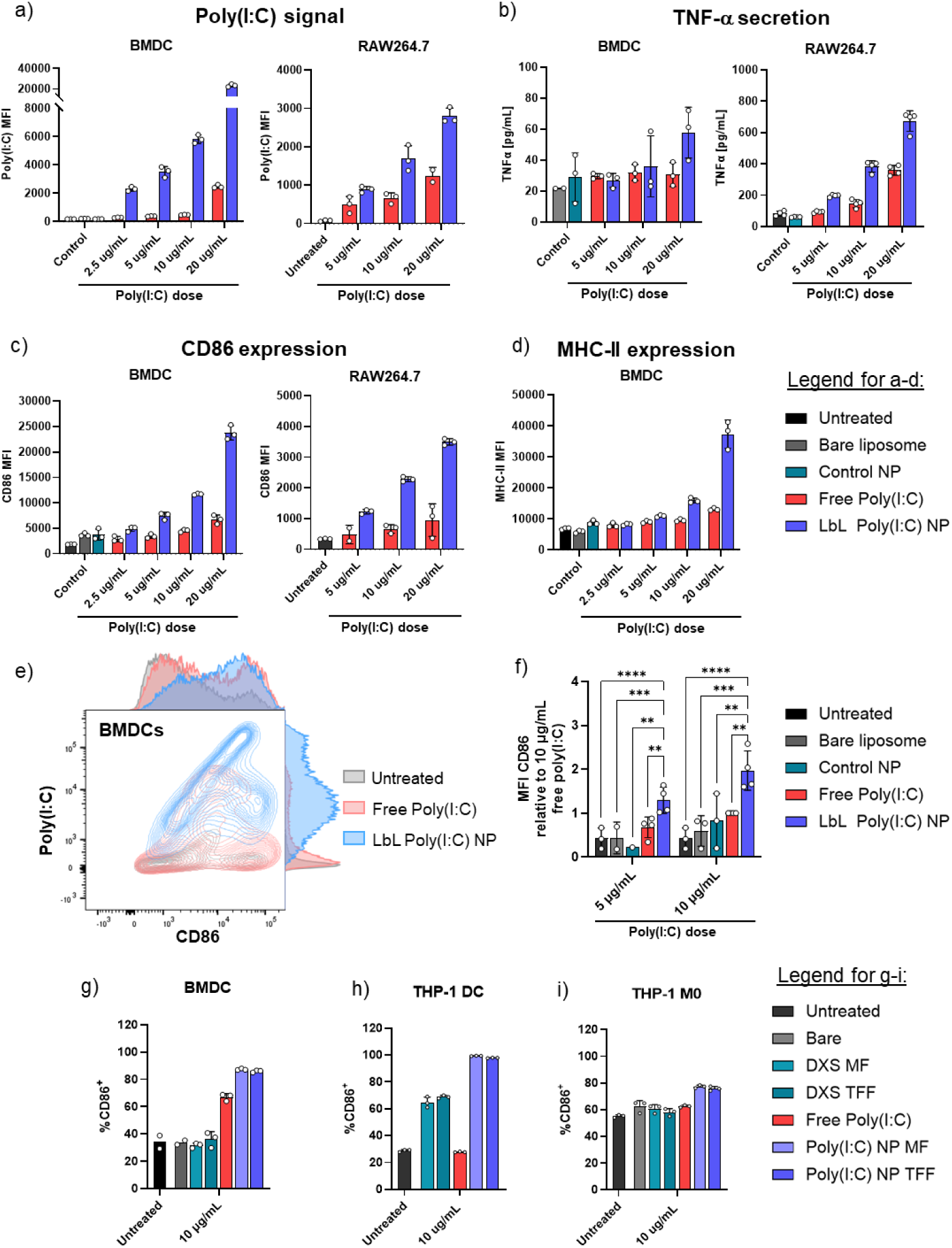
Targeted LbL NP enhances poly(I:C) delivery and APC activation. Bone marrow derived dendritic cells (BMDCs) and RAW264.7 macrophages were incubated with bare liposomes, empty control PLR/DXS LbL NPs (Control NP), free poly(I:C) or LbL Poly(I:C) NPs at varying poly(I:C) doses for 24 hrs. 0.5% Cy5-poly(I:C) was included in the formulation to enable visualization of the therapeutic. a) Poly(I:C) signal measured using flow cytometry for BMDCs and RAW264.7 cells. b) TNF-a secretion in cell supernatant of BMDCs and RAW264.7 cells quantified using ELISA. c) CD86 costimulatory molecule expression quantified using flow cytometry for BMDCs and RAW264.7 cells. d) MHC-II expression of BMDCs evaluated using flow cytometry. e) Representative flow plot of the median fluorescence intensity of poly(I:C)-Cy5 association (y-axis) and CD86 expression (x-axis) in BMDCs dosed at 10 μg/mL poly(I:C) as free poly(I:C) (red) or encapsulated in NP (blue) compared to untreated cells (gray). Histograms of poly(I:C) and CD86 signal are depicted on the outer vertical and horizontal edges, respectively, of the flow plot. f) Median fluorescence intensity (MFI) of CD86 expression across 3-4 independent cultures of BMDCs; data is normalized to the CD86 MFI for the 10 μg/mL free poly(I:C) dose for each replicate. BMDCs were gated as CD11c^+^MHC-II^+^ cells. Percentage of g) BMDCs, h) THP-1 derived DCs, and i) THP-1 derived macrophages that are CD86^+^ 24hr after treatment with bare liposomes, control NPs or Poly(I:C) NPs assembled using microfluidics (MF) or bulk layering+TFF (TFF). N = 3 technical replicates except where noted. Statistical significance determined using Two-way ANOVA with Tukey’s multiple comparisons test in i. (*p < 0.05, **p < 0.01, ***p < 0.001, ****p < 0.0001).

TFF– and MF-assembled LbL Poly(I:C) NPs were directly compared across multiple models of APCs to determine if cellular activation was comparable across the two methods of formulation. In murine BMDCs, human THP-1 polarized macrophages and polarized DCs, TFF and MF assembled LbL Poly(I:C) NPs resulted in similar levels of cellular activation measured by CD86 expression as well as poly(I:C) delivery (**Fig. 4g-i, S13-14**). Due to the similar bioactivity of the MF and TFF LbL Poly(I:C) NPs, but better scalability and size control of the MF NPs described above, the MF NPs were used for subsequent *in vivo* studies.

We briefly investigated the effect of using different complexing polycations (layers 1 and 3) in the LbL film for the Poly(I:C) NPs. For implementation as the complexing polycation, PLR, poly-L-lysine (PLK), and linear polyethyleneimine (PEI) were compared. PLR LbL NPs exhibited the most stable size while PLK and PEI LbL NPs were >250 nm and PDI >0.2 (**Fig. S15**). PLK LbL NPs overall performed similarly to PLR LbL NPs. PEI conferred an advantage to inducing CD86 expression in DCs and MHC-II expression in RAW264.7 macrophages as well as TNF-α and IFN-β secretion (**Fig. S16-17**). However, the LbL PEI NPs also exhibited decreases in cell viability at the 20 μg/mL dose in BMDCs unlike the PLR and PLK LbL NPs (**Fig. S14a**). PEI may serve as a stronger transfection agent as it improved IFN-β secretion, though at low levels, which is more strongly induced by cytosolic dsRNA sensors^58^, but has known toxicity issues^106^. Due to the substantially improved stability and lower cytotoxicity of PLR LbL NPs, changing the polycation was not pursued further, but it is an interesting formulation variable that could be tuned in future work.

To further understand the delivery mechanism of the NP system, we assessed the trafficking of poly(I:C) and the liposomal NP core in BMDCs. After a 4hr incubation with LbL Poly(I:C) NPs, poly(I:C) was internalized and co-localized with the NP core (**Fig. 5a**). To further probe the intracellular trafficking and localization, RAW264.7 cells were incubated with LbL Poly(I:C) NPs for 1hr and 4hr and subsequently stained for endosomes (EEA1) and lysosomes (LAMP-1). After 1hr, poly(I:C) can be found to co-localize with TLR3 expressed on the cell surface (**Fig. S18**). After 4hr, the same timepoint as the BMDCs, poly(I:C) is largely internalized within RAW264.7 cells and co-localizes with the NP core (**Figure S19**). After 4hr, the poly(I:C) can be found as punctate signal in vesicles, some of which are LAMP1^+^ lysosomes (**Fig. S19**) or EEA1^+^ endosomes (**Fig. S20a-b**) or more diffuse in the cytosol (**Fig. S19-20**). Importantly, the LbL NP delivered poly(I:C) was confirmed to co-localize with internal dsRNA sensors that should interact with poly(I:C)—endosomal TLR3 (**Fig. 5b, S20c**) and cytosolic RIG-I (**Fig. 5c**), with examples pointed out by yellow arrows. Of note, the poly(I:C) signal that co-localizes with RIG-I is lower, likely because the poly(I:C) has escaped into the cytosol and is therefore more diffuse.

**Fig. 5.**
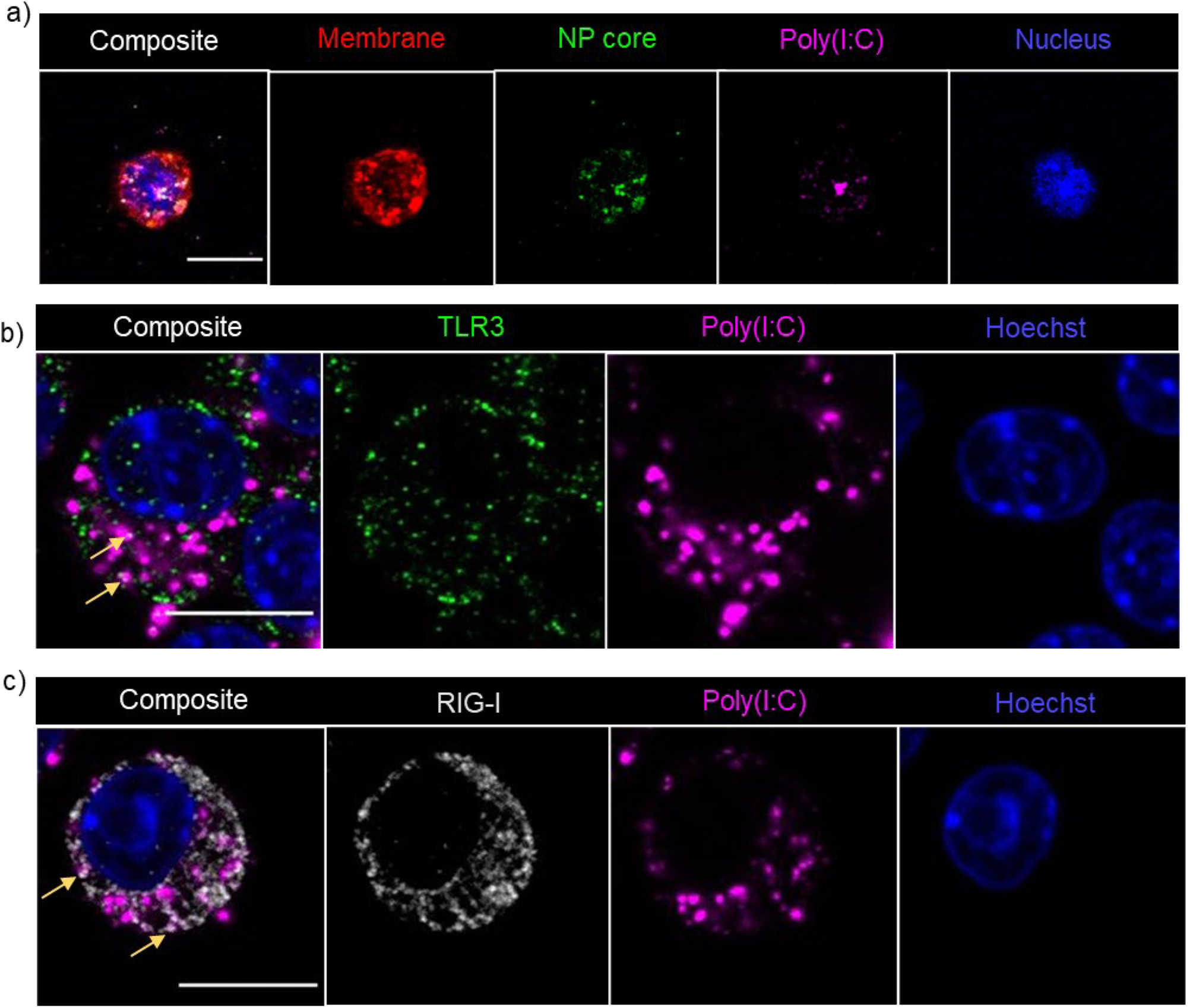
LbL poly(I:C) NP localizes intracellularly and interacts with dsRNA sensors. a) Confocal microscopy analysis (Z-projection) of BMDCs treated with LbL Poly(I:C) NPs at 10 μg/mL (15 μg/mL lipid dose) for 4 hrs. Cells were stained with wheat germ agglutinin-AF555 to visualize the membrane (red) and Hoechst to visualize nuclei (blue). NBD-labeled lipid was included in the liposome formulation to enable visualization of the liposomal core (green) and 2.5% Cy5-poly(I:C) was included to enable its visualization (magenta). BMDCs were enriched using negative selection magnetic-activated cell sorting prior to treating cells. Images were acquired on an Olympus FV1200 with 60X objective. b-c) Confocal microscopy (z-slice) of RAW264.7 cells treated with LbL Poly(I:C) NPs at 10 μg/mL for 4 hr. Cells were stained with Hoechst to visualize nuclei (blue) and 5% Cy5-poly(I:C) was included to enable its visualization (magenta). Poly(I:C) is internalized and can found punctate in vesicles and more diffuse through the cytosol. b) Instances of poly(I:C) co-localization with endosomal dsRNA sensor TLR3 (green) are observed (yellow arrows). c) Poly(I:C) co-localizes with RIG-I (gray), a cytosolic sensor for dsRNA (yellow arrows). The co-localized poly(I:C) signal is low, likely because it has escaped vesicular compartments and is more diffuse/less concentrated in the cytosol. Images were acquired on an Evident FV4000 with 100X objective. Scale bars = 10 μm.

### Poly(I:C) LbL NPs reduce toxicity of poly(I:C) therapeutic

Because ovarian cancer is a disease disseminated throughout the peritoneal space, this LbL Poly(I:C) NP therapy was administered intraperitoneally (IP). This route was chosen to maximize the potential for targeting APCs in the tumor microenvironment as IP administration has demonstrated superior NP accumulation within ovarian tumors compared to intravenous administration^44^. As intraperitoneally administered agents eventually enter into systemic circulation and immune agonists have demonstrated toxicity upon systemic accumulation, this NP design aims to minimize systemic exposure of the therapeutic and retain it in the peritoneal space and in order to mitigate toxicity.

Poly(I:C)-based therapeutics have demonstrated various toxicities upon systemic accumulation including transient increase in serum cytokine levels, decreased white blood cell counts (WBC), decline in platelet levels, fever, elevated liver enzymes, liver injury, and weight loss^19, 107, 108^. Accordingly, we evaluated the serum accumulation of poly(I:C) and systemic toxicity using several metrics including WBC counts, serum cytokines, liver enzymes, and organ histology to assess whether the LbL NP formulation of poly(I:C) can reduce the toxicity of the drug and improve the therapeutic window.

BPPNM tumor bearing mice were administered a single dose of soluble or NP encapsulated poly(I:C) intraperitoneally. The NP reduced the rapid clearance of poly(I:C) into the serum as exhibited by lower poly(I:C) signal in the serum at 1hr (**Fig. 6a**). Both free and NP-encapsulated poly(I:C) induced a transient decline in WBC 24hr post-injection compared to controls (**Fig. 6b**), but this largely recovered by 48hr (**Fig. 6c, Fig. S21a-d**). There were no significant differences in serum liver enzymes for both treatment groups at 24 and 48hr (**Fig. S21e-f**), correlating with an absence of treatment-related histopathologic findings in the liver (**Fig. S22**).

**Fig. 6.**
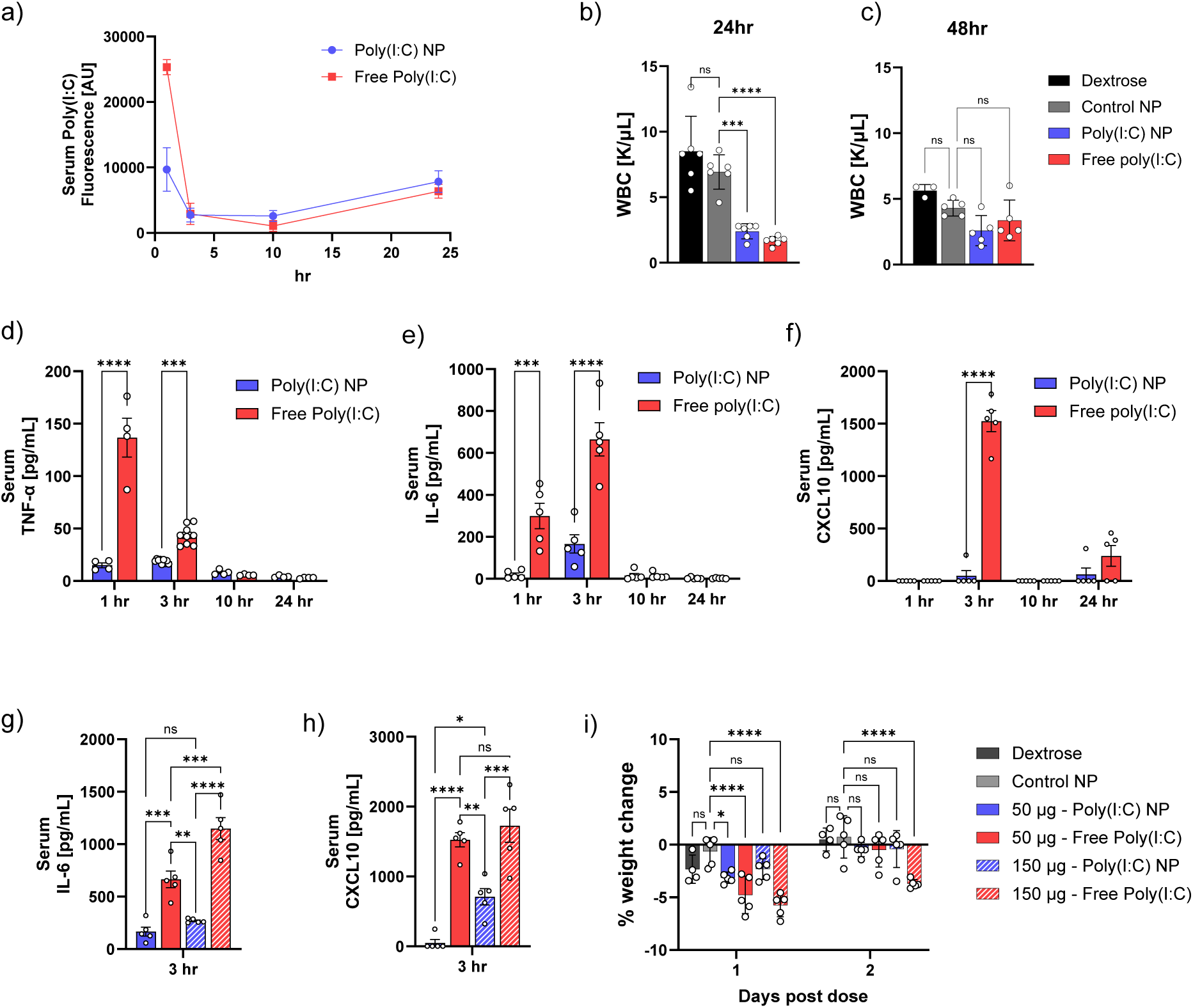
LbL NP improves toxicity profile of poly(I:C) therapeutic. BPPNM-tumor bearing mice were injected intraperitoneally with poly(I:C). a) Pharmacokinetic (PK) analysis of Cy5-poly(I:C) in the serum over time after 50 μg dose of poly(I:C). Serum from dextrose treated mice was used to subtract background fluorescence, mean±SD (N=4). White blood cell counts (WBC) b) 24hr (N=6) and c) 48hr (N=5 for all except Dextrose. N=3 for Dextrose group) after 50 μg dose of poly(I:C), mean±SD. Serum cytokine measurements at 1, 3, 10, and 24hr post-injection of 50 μg poly(I:C) of d) TNF-α (N=4-9 mice per timepoint), e) IL-6 (N=5), and f) CXCL10 (N=5), mean±SEM. g) IL-6 and h) CXCL10 serum cytokine measurements at 3 hr post-injection comparing 50 μg and 150 μg poly(I:C) dose—3 hr data for 50 μg doses reproduced in g-h from e-f to enable comparisons. i) Weight changes (n = 4-5) following IP administration of poly(I:C) as free drug or NP, control NP, or dextrose, mean±SD. Statistical significance determined using One-way ANOVA with Tukey’s multiple comparisons test in b-c, g-h); Mixed-effects model with Bonferroni’s multiple comparisons test in d); Two-way ANOVA with Tukey’s multiple comparisons test in e-f); and Two-way ANOVA with Bonferroni’s multiple comparisons test compared to Control NP in i). (*p < 0.05, **p < 0.01, ***p < 0.001, ****p < 0.0001).

Serum was collected and analyzed for pro-inflammatory cytokines that are secreted downstream of TLR3 and RIG-I activation. For the equivalent 50 μg dose of poly(I:C), serum levels of TNF-α were significantly lower for the Poly(I:C) NP compared to free drug 1hr and 3hr after dosing, peaking at 1hr at which the NP induced a 9-fold lower level of the cytokine compared to free drug (**Fig. 6d**). Kinetics were delayed for IL-6 (**Fig. 6e**) and CXCL10 (**Fig. 6f**), with both also showing higher serum levels for the free drug and peaking at 3hr. At 3hr, the free drug induced 4-fold and 31-fold higher levels of IL-6 and CXCL10, respectively, compared to the LbL Poly(I:C) NP (**Fig. 6e-f**). For baseline cytokine levels, concentrations for the dextrose vehicle control were typically below the limit of quantification (**Fig. S23**).

Comparing the 50 μg and 150 μg doses of poly(I:C), the serum levels of IL-6 (**Fig. 6g**) and CXCL10 (**Fig. 6h**) were ∼2.5X higher for the free drug at the 50 μg dose compared to the Poly(I:C) NP at 150 μg—a three-fold higher poly(I:C) dose—indicating an improvement in therapeutic window with NP encapsulation. Weight loss for the free drug group was also more significant at both doses, and animals were slower to recover to their baseline weight (**Fig. 6i**). Notably, the 150 μg free poly(I:C) group was the only group to not recover to baseline weights two days after dosing.

Given that toxicities can compound over multiple doses, changes in complete blood counts (CBCs) and weight were assessed in response to a daily dosing regimen. BPPNM-tumor bearing mice were given four daily doses of 50 μg free or NP-encapsulated poly(I:C). After 24hrs post final dose, the Poly(I:C) NP group maintained higher WBCs, though both the NP and free drug groups recovered by 48hr (**Fig. S24c-f**). Additionally, weight loss tended to be more dramatic and prolonged for the free poly(I:C) group compared to the NP (**Fig. S24k**). These results demonstrate that the NP encapsulation of poly(I:C) reduces toxicity, widens the therapeutic window, and makes repeat dosing safer compared to the free soluble drug.

### Poly(I:C) LbL NPs improve pharmacokinetics and APC activation in tumor environment

Given the improved safety profile and lower systemic accumulation, we evaluated whether the targeted NP design alters the pharmacokinetics of poly(I:C) in the peritoneal space and localizes the therapeutic to the tumor environment. BPPNM tumor-bearing mice were dosed intraperitoneally with Cy-5 labeled poly(I:C) as free soluble drug or encapsulated in LbL NPs (**Fig. 7a**). The LbL NP formulation increased the retention of poly(I:C) in the peritoneal space, with ∼2-fold higher signal remaining 1-3 days after dosing compared to free drug as measured by IVIS (**Fig. 7b**; raw data **Fig. S25-26**). This increase in poly(I:C) retention was confirmed using cryo-fluorescence tomography (CFT) 24hr after dosing (**Fig. 7c**). Interestingly, poly(I:C) accumulation appeared to be higher in the mediastinal lymph nodes, which drain the peritoneal space, for the Poly(I:C) NP compared to free drug. This could be due to the APC targeting of the formulation as lymph nodes are rich in these cell populations.

**Fig. 7.**
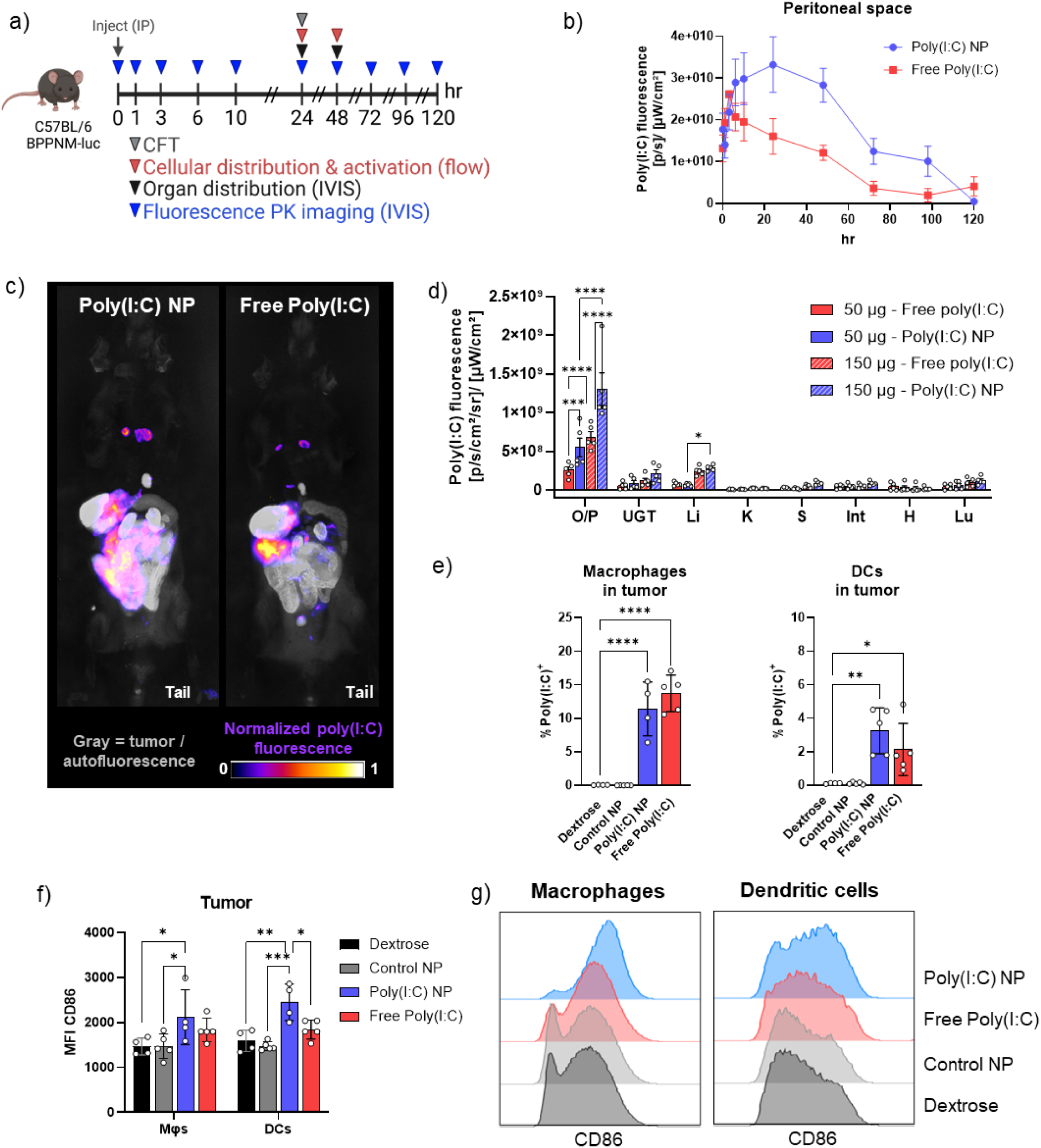
LbL NP encapsulation improves poly(I:C) pharmacokinetics, tumor accumulation, and APC activation. a) Schematic of experimental outline. BPPNM-tumor bearing C57BL/6 mice were dosed intraperitoneally with poly(I:C) as free drug or encapsulated in LbL NP to evaluate the pharmacokinetics, biodistribution, and activation of antigen presenting cells (APCs). Cy5-poly(I:C) was included in the formulations to enable visualization of poly(I:C) using in vivo imaging system (IVIS), cryo-fluorescence tomography (CFT), and flow cytometry. The activation of APCs in the tumor and ascites was additionally assessed using flow cytometry. b) Fluorescence of Cy5-poly(I:C) in the peritoneal space (total radiant efficiency) measured using IVIS after intraperitoneal administration of 50 μg poly(I:C) as free drug or encapsulated in LbL NP, mean±SEM (N=4). c) Distribution of poly(I:C) 24 hr after dosing assessed using CFT (purple-yellow-white). BPPNM tumors, which express GFP, were visualized in the GFP channel (gray) which also has some contribution from background autofluorescence. d) Poly(I:C) fluorescence (average radiant efficiency) in organs 48 hr after administration at 50 or 150 μg dose, mean±SEM (N=5). e) Percentage of macrophages (Mφs, CD45^+^F4/80^+^Gr-1^low^) and dendritic cells (DCs, CD45^+^F4/80^-^CD11c^+^MHC-II^+^) in the tumor that are positive for poly(I:C), mean±SD (N=4-5). f) Median fluorescence intensity (MFI) of CD86 expression on Mφs and DCs in the tumor, mean±SD (N=4-5). g) Representative histograms of CD86 expression on macrophages and dendritic cells. Gating strategy for cells in e-g depicted in Fig. S31. O/P – omentum/pancreas; UGT – upper genital tract; Li – liver; K – kidney; S – spleen; Int – intestines; H – heart; Lu – lungs. Statistical significance in e) determined using One-way ANOVA compared to dextrose control with Dunnett’s multiple comparison test, and in d, f) using Two-way ANOVA with Tukey’s multiple comparison test. (*p < 0.05, **p < 0.01, ***p < 0.001, ****p < 0.0001).

To evaluate the tumor and off-target accumulation of poly(I:C), poly(I:C) fluorescence was quantified using *ex vivo* IVIS imaging. At 24hr, the Poly(I:C) NP tended to have higher accumulation in the omental tumor compared to free poly(I:C) but was not statistically significant (**Fig. S27**). Given that the NP was found to prolong retention of poly(I:C) in the peritoneal space, we hypothesized that there may be greater differences in tumor accumulation between the two groups at later timepoints. At 48hr, the NP had significantly higher accumulation in the omental tumor for both the 50 μg and 150 μg dose of poly(I:C) compared to free drug (**Fig. 7d-e**; raw data in **Fig. S28**). A dose response was observed, with ∼3X higher poly(I:C) signal in the tumor for the 150 μg NP dose compared to 50 μg NP dose.

To explore whether APCs—the cell population of interest—are being targeted, tumors and ascites were dissociated and poly(I:C) fluorescence and CD86 expression were evaluated using flow cytometry. After 24hr, free poly(I:C) and Poly(I:C) NP comparably increased poly(I:C) delivery and functional activation (CD86^+^) of macrophages (CD45^+^F4/80+Gr-1^low^) and DCs (CD45^+^F4/80^-^CD11c^+^MHC-II^+^) in the tumor and ascites compared to dextrose and control NP (**Fig. S29,** gating strategy described in **S31**). While after 48hr, free poly(I:C) and Poly(I:C) NP comparably increased the percentage of macrophages in the tumor positive for poly(I:C), the NP had a more significant increase in the percentage of tumor-associated DCs positive for poly(I:C) (**Fig. 7f**). Additionally, after 48hr, the Poly(I:C) NP achieved significantly improved expression of CD86 on macrophages and DCs in the tumor over the controls and free poly(I:C) (**Fig. 7g**). This highlights that the prolonged retention of poly(I:C) in the peritoneal space by the NP can prolong activation of these tumor-associated APCs. In the ascites, after 48hr, the Poly(I:C) NP yielded a higher percentage of macrophages positive for poly(I:C) and CD86 expression at the 50 and 150 μg dose (**Fig. S30d, f**). There was also a trend of decreased levels of myeloid-derived suppressor cells (MDSCs, CD45^+^CD11b^+^Gr-1^hi^), which is a cell type that contributes to undesirable immunosuppression in the context of cancer^109^, in the tumor and ascites treated with the Poly(I:C) NP compared to free poly(I:C) though not statistically significant (**Fig. S30h-i**). These results indicate that the Poly(I:C) NP can achieve functional activation of APCs in the tumor environment and maintain this activation for longer, while mitigating systemic immune activation and toxicity.

### Poly(I:C) NP synergizes with chemotherapy to improve survival in metastatic ovarian cancer model

Combinations of immune agonists with chemotherapies have proven beneficial as a chemoimmunotherapy approach^110, 111^. Chemotherapy agents that induce immunogenic cell death (ICD) are particularly advantageous as they have immunomodulatory effects, including triggering the release of inflammatory molecules such as high-mobility group box 1 (HMGB1), the secretion of ATP which attracts DCs, and the increase in expression of calreticulin on the cell membrane which facilitates phagocytic uptake^69, 112^. We aimed to combine the Poly(I:C) LbL NP with the ICD-inducing chemotherapeutic doxorubicin (Dox) for a synergistic effect in treating ovarian cancer. Liposomal Dox is currently used as a second-line therapy for recurrent ovarian cancer^113, 114^ and has demonstrated improved safety profiles over free drug^115^. The BPPNM model of ovarian cancer was validated to be sensitive to Dox (**Fig. S32**). Additionally, the combination of poly(I:C) and Dox has shown positive effects in other models of cancer^67^.

Given that poly(I:C) can induce apoptosis and secretion of inflammatory cytokines from some cancer cells through interactions with endosomal TLR3 and/or cytosolic RIG-I and MDA5 receptors^116, 117^, we tested the therapeutic effects of LbL Poly(I:C) NPs on BPPNM ovarian cells, the cells used for the *in vivo* model in subsequent analyses of the therapeutic efficacy of LbL Poly(I:C) NPs. At a dose of 1 or 10 µg/mL DXS and PLD coated LbL Poly(I:C) NPs did not elicit changes in cell viability nor substantial IFN-β or CXCL10 production (**Fig. S33**). This could be due to this poly(I:C) being a lower molecular weight than other studies, as MDA5 activation is dependent on dsRNA length^58^; alternatively, BPPNM cells may rely more strongly on signaling from cytosolic receptors rather than endosomal TLR3, as these NPs have demonstrated stronger retention in endosomes—for APCs—compared to cytosolic delivery (**Fig. 5, S19-20**). Therefore, any observed therapeutic benefits of the LbL Poly(I:C) NPs will be due to immune cell activation and not direct killing or immunomodulation of cancer cells.

We developed a LbL liposomal NP encapsulating Dox for tumor delivery of the therapeutic (Dox NP). Dox was incorporated into the liposome via active loading with a pH gradient and subsequently layered with PLR followed by PLD for tumor-targeting capabilities (**Table S8**). The NPs had stable size and charge and achieved high weight loading of Dox (**Fig. S34**). The bare and LbL Dox-loaded liposomes maintained bioactivity of the drug in vitro, although the IC_50_ was slightly lower compared to free drug (**Fig. S35, Table S9**).

To assess the efficacy of adding poly(I:C) to chemotherapy, the BPPNM model was established and mice were treated with Dox NP (2 mg/kg) with or without the addition of Poly(I:C) NP (50 μg) or 5% dextrose vehicle control (Untreated) twice per week for two weeks starting 10 days after tumor inoculation (**Fig. 8a**). By day 10, the model had already developed significant metastatic disease. Two independent biological replicate studies were performed in this model and the data was combined. As the tumor growth kinetics differed between the two cohorts of mice, survival data was normalized relative to the median survival of the Untreated group for each cohort. Data for each replicate study is in **Fig. S36-37** and **Table S10**. Interestingly, the Poly(I:C) NP alone performed similarly to Untreated, likely because Poly(I:C) alone does not induce cancer cell death as shown previously (**Fig. S36**). Dox NP alone slowed tumor growth, and the addition of Poly(I:C) NP further reduced tumor growth (**Fig. 8b-c**). This led to a significant extension in survival, demonstrating the potential for this combination therapy (**Fig. 8d-e**). These results were consistent across both replicate experiments (**Fig. S36**). While the combination of Dox NP + Free Poly(I:C) performed similarly to Dox NP + Poly(I:C) NP (**Fig. 8f**), the Poly(I:C) NP is advantageous due to the demonstrated superior toxicity profile.

**Fig. 8.**
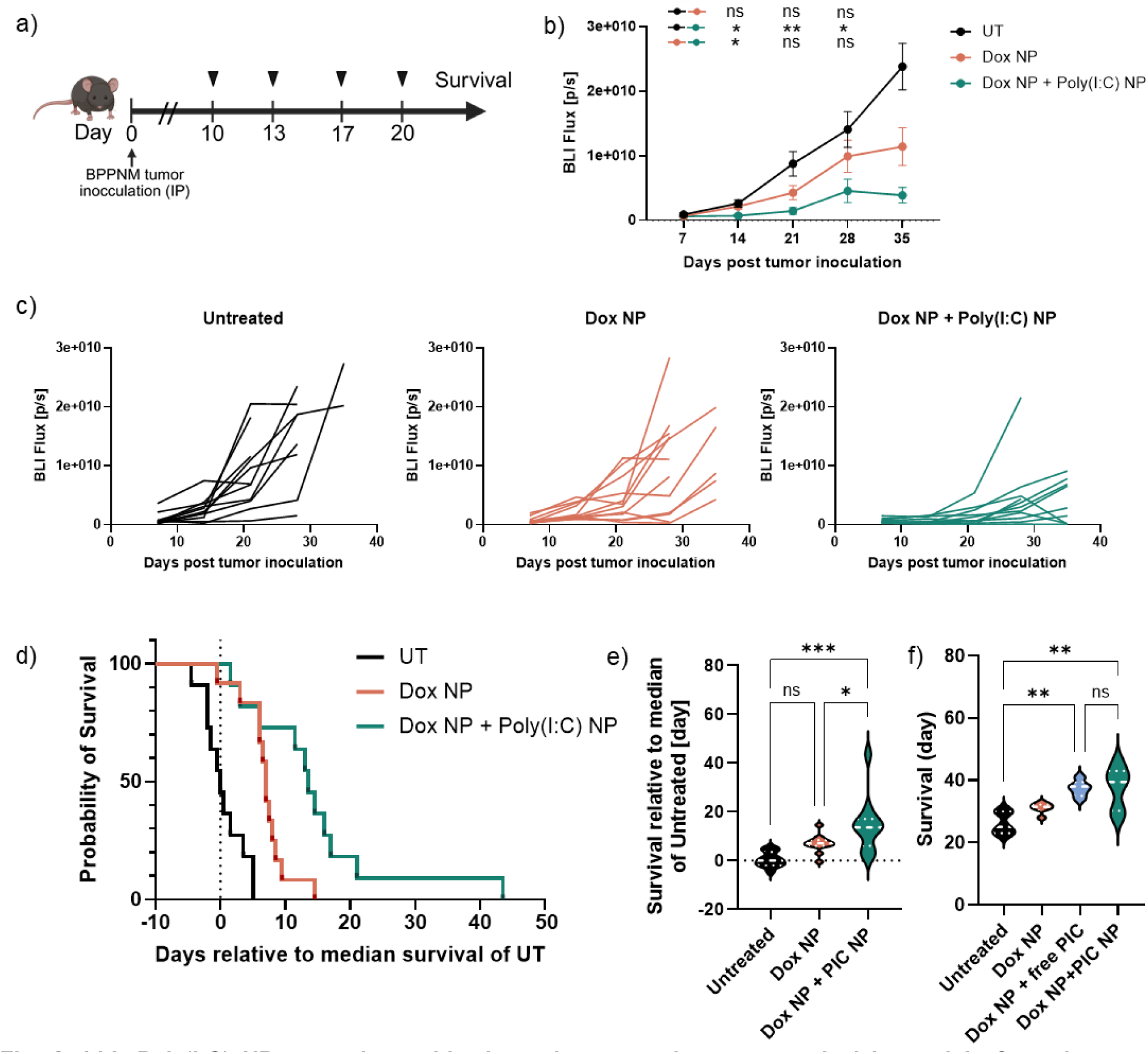
LbL Poly(I:C) NP synergizes with chemotherapy to improve survival in model of ovarian cancer. Schematic of experiment. C57BL/6 mice were injected intraperitoneally with 3×10^6^ BPPNM-luc cells. Ten days post tumor inoculation, mice were dosed intraperitoneally twice weekly with the Dox NP (2 mg/kg) with or without addition of Poly(I:C) NP (50 μg), or 5% dextrose vehicle control (Untreated, UT) on Days 10, 13, 17, and 20. Data in b-e) is combined data from two independent studies (N=11-12 per group in total). b) Tumor burden was measured by bioluminescent IVIS imaging, mean ± SD. Statistical differences between groups plotted above. c) Tumor bioluminescence over time for each subject. d) Survival analysis of mice relative to the median survival day of the respective Untreated control for each cohort of mice for both experiments combined, to normalize for differences in tumor growth between the studies. e) Day of survival relative to median of Untreated control for both experiments combined. Poly(I:C) denoted as PIC. f) Day of survival for same cohort of mice. In e-f) Dashed lines denote median and dotted lines denote quartiles. Statistical significance in b) determined using Mixed-effects analysis with Tukey’s multiple comparisons test and in e-f) using One-way ANOVA with Tukey’s multiple comparisons test. (*p < 0.05, **p < 0.01, ***p < 0.001, ****p < 0.0001).

## Conclusions

In this work, we demonstrate a NP design strategy to improve the retention, tissue distribution, cellular targeting, APC activation, and safety of the immune agonist poly(I:C) via the encapsulation of the therapeutic in a rationally designed tetralayer LbL NP system. We optimized the selection of NP surface chemistry to promote intracellular delivery and targeting of the APC cell populations and found dextran sulfate to be a promising surface chemistry candidate to promote specificity for APCs. The LbL encapsulation of poly(I:C) significantly improved the retention of the therapeutic in the peritoneal space, with about 2-fold higher poly(I:C) remaining in the peritoneal space 1-2 days after administration for the NP compared to free poly(I:C). By retaining the drug within the peritoneal space where the cancer is located and reducing rapid clearance into systemic circulation, this markedly improved the safety and therapeutic window. Compared to free drug, the NP reduced the peak in the serum levels of the inflammatory cytokines TNFα, IL-6, and CXCL10 by 9-, 4-, and 31-fold for the 50 μg dose. Even at a 3-times higher poly(I:C) dose of 150 μg, the NP maintained lower serum cytokine levels compared to 50 μg of free poly(I:C), demonstrating that the NP widens the therapeutic window. As high serum levels of these cytokines induce toxicity, we accordingly observed lower weight loss in NP treated mice compared to free drug, including upon repeat daily dosing, indicating that this NP strategy improves the safety and ability for repeat dosing of this therapeutic. The addition of the Poly(I:C) NP immunotherapy to the ICD-inducing chemotherapeutic doxorubicin further slowed tumor growth and more than doubled survival in an aggressive metastatic model of ovarian cancer. By enhancing the safety and expanding the applicability of this potent TLR3 agonist to a non-localized administration route, future work may expand on this targeted LbL NP strategy for the delivery of other immunomodulatory agents that are currently limited by toxicity concerns.

## Methods

### Materials and Reagents

Dextran sulfate (DXS, 15 kDa, Cat#51227), fucoidan (#F8190), polyacrylic acid (PAA, 15 kDa, #416037), poly(diallyldimethylammonium chloride) solution (PDADMAC, 100kD, 20 wt% in water; #409014), 3 M sodium acetate (#S7899), ammonium sulfate (#A4418) doxorubicin hydrochloride (#PHR1789), Eppendorf DNA LoBind tubes, Tween 20 (#P9416), Collagenase from Clostridium histolyticum Type V (#C9263), DNase I (Roche, #11284932001), Hyaluronidase Type I-S (#H3506), bovine serum albumin (BSA) (#A2934), methanol (#34860), heat-inactivated fetal bovine serum (#F4135) were purchased from Millipore-Sigma. Poly-L-arginine (PLR) hydrochloride (PLR200, 38.5 kDa), poly-L-lysine (PLK) hydrochloride (PLK250, 41 kDa), poly-L-aspartic acid (PLD) sodium salt (PLD100, 14 kDa), and poly-L-glutamic acid (PLE) sodium salt (PLE100, 15 kDa), methoxy-poly(ethylene glycol)-block-poly(L-glutamic acid sodium salt) (PLE-PEG) (5 kDa PEG, 15 kDa PLE) were purchased from Alamanda Polymers. Linear polyethylenimine MW∼25kD (#23966) was purchased form Polysciences Inc. Chondroitin sulfate A (CSA, 10-30 kDa, #YC15288) and carboxymethyl dextran (CMDx, 10-20kDa, # YC58709) were purchased from Carbosynth. Hyaluronic acid (HA, 20 kDa, #HA20K) was purchased from Lifecore Biomedical. Very low-viscosity sodium alginate, with a MW < 75 kDa, was purchased from NovaMatrix.

Roswell Park Memorial Institute (RPMI) 1640 with glutamine and HEPES (#10-041), and Dulbecco′s Modified Eagle′s Medium (DMEM) with high glucose, glutamine, sodium pyruvate (#10-013), and 1x Phosphate Buffered Saline (PBS) without calcium and magnesium (#21-040-CV) was purchased from Corning. ATCC modification RPMI 1640 with 2 mM L-glutamine, 10 mM HEPES, 1 mM sodium pyruvate, 4500 mg/L glucose, and 1500 mg/L sodium bicarbonate (#30-2001) was purchased from ATCC. Sodium chloride (#BP358-1), 5 M molecular biology grade NaCl solution (Invitrogen, #AM9760G), 1 M bioreagent-grade HEPES (N-2-hydroxyethylpiperazine-N-2-ethane sulfonic acid) (Gibco, # 15630080), ACK Lysis Buffer (Gibco, # A1049201), 0.5 M EDTA, pH 8.0 (Invitrogen, #AM9260G), Fetal bovine serum (FBS) (Gibco, #A52567-01), 10,000 U/mL penicillin and 10,000 µg/mL streptomycin solution (Gibco, #15140122), 0.25% Trypsin-EDTA (Gibco, # 25200056), Insulin-Transferrin-Selenium (ITS –G) (100X) (Gibco, #41-400-045), DMEM with high glucose, HEPES (Gibco #12430062), DMEM:F12 1:1 with glutamine and HEPES (Cytiva, # SH3002301), 16% formaldehyde (#28908), UltraComp eBeads™ Plus Compensation Beads (#01-3333-42), D-Luciferin Sodium Salt (GoldBio, #LUCNA), Pierce High Capacity Endotoxin Removal Spin Columns (#88276), Pierce Endotoxin-Free Water (#A43883), dimethyl sulfoxide (DMSO) (#BP231100) were purchased from ThermoFisher and Fisher Scientific. EGM®-2 Bulletkit was purchased from Lonza (#CC-3162).

Tangential flow filtration (TFF) membranes D02-E100-05-N, C02-E100-05-N, D02-E750-05-N, C02-E750-05-N, reservoirs (15 mL, 50 mL, 250 mL) and Float-A-Lyzer G2, 8 – 10 kD (#G235031) were purchased from Repligen. Teflon-coated tubing, sizes 14 and 16, used for filtration, was purchased from Saint-Gobain (Waltham, MA). UV-transparent low-volume cuvettes (67.758) for the Malvern Zetasizer were purchased from Sarstedt and DTS1070 folded capillary cells from Malvern. Cholesterol, 1,2-distearoyl-sn-glycero-3-phosphocholine (DSPC), 1,2-distearoyl-sn-glycero-3-phospho-(1′-rac-glycerol) (DPSG), 1,2-distearoyl-sn-glycero-3-phosphocholine (N-azidoethyl) (18:0 azidoethyl PC), 1,2-distearoyl-sn-glycero-3-phosphoethanolamine-N-(7-nitro-2-1,3-benzoxadiazol-4-yl) (ammonium salt) (NBD-DSPE), 1-palmitoyl-2-oleoyl-glycero-3-phosphocholine (POPC), and 1-palmitoyl-2-oleoyl-sn-glycero-3-phospho-(1’-rac-glycerol) (sodium salt) (POPG) were purchased from Avanti Polar Lipids, Inc.

Membranes used for liposome extrusion were purchased from GE Healthcare. Low molecular weight Poly(I:C) (#tlrl-picw) was purchased from Invivogen. Indocyanine green dibenzoazacyclooctyne (ICG DBCO, #RL-2870) was purchased from Iris Biotech. BDP 650/665 (#AE4F0) and BDP TMR (#124F0) with DBCO handles were purchased from Lumiprobe. ICG-DBCO (#RL-2870.0005) was purchased from Iris Biotech. Type I Ultrapure water generated with a Milli-Q IQ 7000 Ultrapure Lab Water System equipped with a Biopak polisher (Millipore) was used. Vented tissue culture treated T25, T75, and T175 flasks, tissue culture treated 96-well plates, were purchased from VWR. Black 96-well plates were purchased from Thermo and Grenier. Ethanol (#76270-156), chloroform ethanol stabilized (#TCC1111), serum Gel collection tubes (Sarstedt, # 41.1500.005), EDTA K3E collection tubes (Sarstedt, #20.1278.100), 20 mL scintillation vials were purchased from VWR. 0.22 μm PES syringe filters were purchased from VWR and Celltreat.

### Nanoparticle preparation and characterization

#### Preparation of empty liposomes

DSPC and cholesterol were dissolved at 25 mg/mL in chloroform and DSPG was dissolved at 25 mg/mL in a 1:1 chloroform/methanol (v/v) mixture. Dye-labeled lipids were dissolved at 0.5-5 mg/mL in chloroform. Liposomes were prepared using the film hydration method. Lipids and cholesterol were mixed in a round-bottom flask (ChemGlass). For both dye-labeled liposome formulations, the DSPC fractions were adjusted based on the amount of dye-lipid incorporated, so the lipid mol ratios used are 40:34:25:1 for NBD labeled and 40.8:34:25.5:0.2 for BDP650, BDP-TMR, or ICG labeled (DSPC:cholesterol:DSPG:dye-lipid) formulations (Table S1). A thin lipid film was formed by evaporating the solvent using a Buchi RotoVap system at 25°C and 50 mbar for at least 30 min. The round-bottom flask was partially submerged in a Branson sonicator bath heated to 60-65°C, and the lipid film was hydrated by adding water, 20 mM HEPES 20 mM sodium chloride, or 20 mM sodium acetate solution to a lipid concentration of 1 mg/mL under sonication. The liposomes were sonicated in the bath for 2 min and subsequently extruded through an Avestin LiposoFast LF-50 extruder complete with a heated jacket connected to a Cole-Parmer Polystat Heated Recirculator Bath to maintain the temperature at 65 °C (above the liposome phase transition temperature). The liposomes were passed through stacked 400 and 200 nm filters once, then through a 100 nm filter twice, and optionally through a 50nm to attain a Z-average diameter of 100-120 nm. The liposomes were then allowed to slowly cool to room temperature for at least 30 minutes.

### Preparation of doxorubicin-loaded liposomes

Doxorubicin (Dox) was actively loaded into liposomes using a pH gradient. In a 20 mL glass scintillation vial, 60 mg POPC, 28 mg cholesterol, and 7.5 mg POPG in chloroform (molar ratio 49:45:6 POPC:cholesterol:POPG) were combined and chloroform removed using a RotoVap. The molar ratio of lipid to cholesterol is similar to that of the non-PEGylated liposomal Dox formulation Myocet (55:45 eggPC:cholesterol)^115, 118^, with POPG included in the formulation to facilitate an anionic surface of the liposomes for subsequent LbL assembly. Subsequently, 800 μL ethanol was added and solution stirred in a 65°C sand bath until fully dissolved, typically 5-10 minutes. To the vial, 7.2 mL of 300 mM ammonium sulfate (pH 5) was added and stirred for 1hr at 65°C then sonicated at 65°C for 20 minutes in a bath sonicator. The liposomes were then extruded (see above) by passing through stacked 400 and 200nm filters once, a 100 nm filter twice, and a 50nm twice. The liposomes were dialyzed in 3.5 kDa dialysis membrane against 300 mM NaCl for 2 hours to remove unencapsulated ammonium sulfate buffer. The liposomes were heated to 65°C under stirring in a sand bath and 10 mg/mL doxorubicin hydrochloride (12 mg, in water) was quickly added at 1:5 Dox:POPC weight ratio and the solution stirred for 1hr at 65°C. The liposomes were cooled to room temperature in a water bath then dialyzed against water for 18hr (3.5 kD MWCO) to remove unencapsulated Dox.

### Layer-by-Layer assembly of nanoparticles

#### Bulk solution layering coupled with TFF

In this method, polymers are electrostatically adsorbed onto nanoparticles by rapidly mixing bulk solutions of the nanoparticles with solutions of polymer of the opposite charge at weight ratios of NP core:polymer that impart sufficient charge conversion, followed by purification of non-adsorbed polymer using TFF before the addition of the subsequent layer. In brief, liposomes were layered by rapidly pipetting in equal volumes of NP solution to polyelectrolyte solution under sonication followed by vigorous vortexing for 10 s. Polyelectrolyte layering solutions were prepared in water, HEPES / sodium chloride or sodium acetate buffers depending on the particular polyelectrolyte. The weight equivalents of each polyelectrolyte to liposome core was optimized to achieve a zeta potential greater than 30 mV or less than –30 mV—as this is a typical zeta potential threshold to achieve colloidal stability^119^—and an acceptable size. Detailed assembly conditions for bilayer empty liposomes (Table S4), tetralayer poly(I:C) NPs (Table S6), and bilayer Dox liposomes (Table S8) can be found in Supplementary information. Layered particles were incubated for at least 30 min on ice before being purified using TFF, as described previously^120^. Polymers were tested for endotoxin and endotoxin removed using Pierce™ High-Capacity Endotoxin Removal Spin Columns when applicable.

TFF was used to purify the NPs after deposition of each polyelectrolyte layer, whereby the solution was connected to a KrosFlo II or KR2i (Repligen) system using Teflon-coated tubing with 100 kDa hollow-fiber filter units or 750 kDa hollow-fiber filter units for poly(I:C) containing formulations due to the large molecular weight of poly(I:C). Depending on the NP volume, NP solutions were passed through either the C02-E100-05-N 100 kDa at 13 mL/min for non-poly(I:C) formulations or C02-E750-05-N 750 kD membrane at 7 mL/min for poly(I:C) formulations—as the lower flow rate imparts less shear on the NPs—with size 14 tubing (<15 mL NPs), or D02-E100-05-N 100 kD at 80 mL/min for non-poly(I:C) formulations or D02-E750-05-N 750 kD membrane at 60 mL/min for poly(I:C) formulations with size 16 tubing (>15 mL NPs). The NPs were purified with 3-5 diafiltration volumes of Milli-Q water, and then concentrated and recovered by running 1-3 mL water through the tubing to improve yield of the particles. The NPs were kept on ice throughout the layering and TFF processes. Following purification, NPs were characterized for size and zeta potential (see below) and stored at 4°C until later use.

#### Microfluidic LbL Assembly

Microfluidic mixing was performed on NanoAssblr Ignite NxGen^TM^ cartridges (Precision Nanosystems). Equivalent volumes of liposomes or polyelectrolyte solution were filled in separate 1, 3, 5, or 10 mL luer-lock syringes, depending on batch size, at the appropriate wt:wt ratio and attached to each port of the microfluidics chip. The liposome and polyelectrolyte solution were simultaneously infused into the chip at a flow rate of 5 mL/min (total flow rate 10 mL/min) using a syringe pump (Pump 11 Elite, Harvard Apparatus). Detailed assembly conditions for tetralayer poly(I:C) NPs (Table S7) can be found in Supplementary information. After the last layer, particles were incubated for at least 30 min on ice before being purified and concentrated using TFF.

### NP Characterization

Size, polydispersity index (PDI), and zeta potential of the nanoparticles were measured using dynamic light scattering (DLS) and electrophoretic light scattering (ELS) respectively using a Malvern Zetasizer Pro Red (λ = 633 nm, 173° detection angle, optical fluorescence filter) or Malvern Zetasizer Nano ZS90s (λ = 633 nm, 90° scattering detector angle). For the liposomes, a refractive index of 1.45 and absorption coefficients of 0.001 was used; and for the dispersant refractive index and viscosity values for water were used (1.330, 0.8872 cP at 25°C). Samples were measured in 50 uL Sarstedt 67.758 ultraviolet cuvettes (for size measurement) or DTS1070 Malvern-folded capillary tubes (for zeta potential measurement).

### Nucleic acid gel to assess poly(I:C) complexation on NP

E-Gel EX 2% Agarose gel (Thermo, #G401002) was loaded with 20 μL sample per well and ran on E-Gel iBase electrophoresis system according to the manufacturer’s instructions and imaged on a ChemiDoc imaging system (BioRad) using SYBR gold setting. The samples included free poly(I:C) at varying concentrations and LbL-Poly(I:C) NPs after assembly of each layer which were concentration matched to 4 μg/mL poly(I:C), or equivalent liposome concentration for the first PLR layer as the NP as this stage does not contain poly(I:C). For the ladder, E-Gel™ Sizing DNA Ladder (Thermo, #10488100) was used undiluted at 20 μL per well.

### In vitro assays

#### Microscopy of BMDCs to assess NP localization

*Preparation of chamber slides:* To enable adherence of BMDCs, 8-chamber slides (Lab-Tek, Cat #155409) were coated with 0.01% poly-L-lysine (Sigma, Cat #RNBK2055) at room-temperature for 15 min. Poly-L-lysine solution was removed and chamber slides were ready to be seeded with BMDCs.

*Preparation of cells:* For fixed cell imaging of NP localization in BMDCs, frozen BMDCs were thawed, subjected to MACS to enrich for a purer DC population, and then resuspended in full BMDC media. BMDCs were plated at 50,000 cells in 90 μL full BMDC media and treated with 10 μL NPs or poly(I:C) in a 96 well non-TC treated V-bottom plate. After 4hr, DCs were centrifuged and rinsed with PBS and then two replicates per condition were combined into one chamber slide well to achieve 100k cells/well. Cells were incubated for 15 min at RT to allow cells to adhere and fixed in 300 μL 4% formaldehyde (Thermo, Cat# 28906) in PBS at RT for 15 min, protected from light. Cells were subsequently stained per below.

*Cell staining without antibody staining*: Fixed cells were washed twice with PBS then stained with 10 µg/mL wheat germ agglutinin-AF488 (Invitrogen, Cat# W11261) or wheat germ agglutinin-AF555 (#W32464) and 1 ug/mL Hoechst (Invitrogen, #H3570) in HBSS (Gibco, Cat# 14175-095) in 300 μL for 10 min at RT. Cells were gently washed twice with PBS then fixed in 4% formaldehyde for 2 min at RT. Cells were stored in PBS at 4°C, covered from light, until ready to image.

#### Microscopy of RAW264.7 to assess NP localization and trafficking

*Preparation of chamber slides:* To improve adherence of cells, 8-chamber slides were coated with rat tail collagen (Sigma #08-115), 300 μL of 50 μg/mL in 0.02 N acetic acid per well at room-temperature for 1 hr. Collagen solution was removed and chamber slides were rinsed with PBS, and were then ready to be seeded with cells.

*Preparation of cells:* RAW264.7 cells were seeded at 25,000 cells/well in 270 μL media and allowed to adhere overnight. Cells were treated with 30 μL of treatment. At the end of the incubation period, cells were rinsed with PBS and fixed in 300 μL 4% formaldehyde in PBS at RT for 15 min, protected from light. Cells were subsequently stained per below.

*Cell staining with intracellular antibodies*: Fixed cells were washed twice with PBS then stained with 5 µg/mL wheat germ agglutinin-AF488 or wheat germ agglutinin-AF555 in 300 μL for 10 min at RT. Cells were gently washed twice with PBS then fixed in 4% formaldehyde for 2 min at RT. Cells were treated with blocked with 0.025% saponin and 5% goat serum in PBS for 1 hr at RT. Blocking buffer was removed and primary antibodies diluted into 300 μL antibody diluent buffer (0.025% saponin and 1% BSA in PBS) were added. Cells were incubated in the primary antibody overnight at 4 °C, then washed three times with PBS. Fluorophore conjugated secondary antibodies or DyLight 554 phalloidin (1:400 dilution) in 0.025% saponin and 1% BSA in PBS were added to each well and incubated for 1 hr at RT. Cells were washed three times with PBS, fixed again in 1% formaldehyde for 5 min at RT, and then washed with PBS. Cells were stained with 1 ug/mL Hoechst in 300 μL for 5 min at RT, washed three times in PBS and stored in PBS at 4°C, covered from light, until ready to image. Antibodies and dilutions can be found in SI.

#### Microscopy Imaging and data processing

Before imaging BMDCs or RAW264.7 cells, PBS was removed and Vectashield Antifade Mounting Medium (#H-1000) added to cover each well. Cells were imaged with a confocal laser-scanning microscope—Olympus FV1200 equipped with 405, 473, 559, and 635 nm lasers or Evident FV4000 equipped with 405, 488, 561, 640 nm, and 730 lasers. Images were acquired with 60x or 100x objectives and processed using ImageJ software.

### Cellular association of nanoparticles

To evaluate the cellular association of different nanoparticle formulations, cells were seeded in tissue culture-treated 96-well plates at the optimal density to achieve cell confluency (see SI for conditions for each cell line). For adherent cell lines, cells were allowed to adhere overnight prior to treatment with nanoparticles. Fluorescently labeled nanoparticles were diluted 10X into the well (10 μL NPs into 90 μL cell media) and incubated with the cells at a final concentration of 15-25 μg/mL of lipid per well in triplicate. At the respective timepoints, cells were washed 1X with PBS and for adherent cell lines, cell were then removed using 0.25% trypsin-EDTA followed by quenching with media. Cells were stained with 1:600 dilution of Zombie Violet or Zombie NIR viability stain (Biolegend #423113, 423105) in 100 μl of PBS for 10 min at room temperature, subsequently fixed in 2% formaldehyde for 20 min at 4°C, and resuspended in flow cytometry buffer (1X PBS, 1% BSA, 2 mM EDTA). Data was collected using BD Symphony A3 or Fortessa cytometers equipped with high-throughput sampler and analyzed with FlowJo version 10.

For comparison of the relative uptake of NP formulations, the NP-associated fluorescence (median fluorescence intensity, MFI) was first normalized to the MFI of bare liposomes (MFI_NP / MFI_bare) for each run as this enabled consistent comparison of the data acquired on different flow cytometers. Then the normalized MFI for each formulation was Z-scored across each cell line and timepoint to better enable comparisons across cell lines as fold changes in NP signal differed substantially across cell lines. Data analysis was performed on the mean normalized NP association for each outer layer and cell line, for each of the 4– and 24-hour datasets. Principal component analysis was used as a dimensionality reduction method, with mean cell line association as features. After standard scaling with scikit-learn, the first two principal components explained over 80% of variance for both 4– and 24-hour data (see Supplemental). Ward clustering was used on the standard-scaled data to obtain the hierarchal clusters presented in Figures 1d and 1e, Ward clustering on the PCA-transformed data yielded similar clusters (see Supplemental).

### Poly(I:C) NP in vitro bioactivity

Treatments were added to a 96-well plate in 10 µL then 50,000 live RAW264.7 cells or BMDCs were added in 90 µL of media and allowed to incubate for 24 h. Flat-bottom tissue-culture treated plates and DMEM supplemented with 10% FBS and 1% Pen/Strep were used for RAW264.7 cells. V-bottom non-tissue culture treated plates and complete cytokine-containing BMDC media (described above) were used for BMDCs. BMDCs were used from fresh or cryopreserved for which frozen vials of cells were thawed in a 37°C bead bath, added to excess media to dilute out DMSO, centrifuged at 500xg for 3 minutes, and resuspended in fresh, warm BMDC media. Live cells were counted, diluted to the desired concentration, and immediately seeded into the plate. After the 24hr incubation, the supernatant was sampled and stored at −20°C until downstream cytokine analysis and cells were processed for flow cytometry analysis.

For flow cytometry processing, RAW264.7 cells were rinsed with PBS and lifted using 0.25% trypsin-EDTA for 5-7 minutes at 37°C followed by quenching with media and transferred to a V-bottom 96-well plate for flow staining. As BMDCs as do not adhere to non-TC treated plates, trypsin was not needed. Cells were rinsed with PBS, stained with Zombie live/dead dye for 10-15 min at room temperature, Fc receptors blocked with anti-CD16/32 for 15 min on ice, stained with antibodies for 30 min on ice, and then fixed in 2% formaldehyde. Antibodies and dilutions can be found in Supplementary information and Table S3.

### Enzyme Linked Immunosorbent Assay (ELISA) Assays

Mouse TNF-α, IFN-β, IL-6, and CXCL10 were quantified using DuoSet ELISA kits (R&D Systems DY410, DY8234, DY406, DY466). Half-area high binding polystyrene 96-well plates (Corning #3690) were coated with capture antibody in PBS overnight at room temperature, washed three times with PBST (PBS + 0.05% Tween 20). Plates were then blocked with 1% BSA in PBS for at least 1hr at room temperature and washed three times with PBST. For serum samples, 1% BSA in PBS was utilized as the sample and standard diluent, while cell media was used as the diluent for cell supernatant samples. Samples were added at 25 μL/well and incubated on the plates for 2 hours at room temperature and then washed three times with PBST. Detection antibody (in 1% BSA in PBS, 25 μL/well) was added to the wells and incubated for 2 hours, washed three times with PBST, followed by incubation with 25 μL/well streptavidin-HRP for 20 min covered from light. Plates were washed four times with PBST and then developed using 1-Step TMB ELISA Substrate Solution (Thermo Scientific #34028) for 20 min covered from light. The solution was stopped using 2 N sulfuric acid and absorbance measured immediately on a plate reader at 450 nm with 540 nm reference wavelength. Capture antibody, detection antibody, streptavidin-HRP, and standards were prepared and used per the manufacturer’s instructions for each lot. Cytokine concentrations were calculated by interpolation using a four-parameter logistic curve-fit of a standard curve (GraphPad PRISM version 10.2.3). The limit of quantification (LOQ) was calculated by adding ten standard deviations to the mean absorbance of the blank and converting this to a concentration using the 4PL or cubic standard curve. For samples less than the LOQ, the concentration was set to zero for plotting and statistics.

### In vivo experiments

#### Animal Care and Use

All animal experiments were approved by the MIT Committee on Animal Care (CAC, protocol number 2404000660) and were conducted under the oversight of the Division of Comparative Medicine (DCM). Female C57BL/6 mice were purchased from Jackson Laboratory and housed at Koch Institute for Integrative Cancer Research at MIT animal facility in cages of no more than five animals with controlled temperature (25°C), 12 h light–dark cycles and free access to food and water.

### BPPNM-luc tumor model establishment

Three million luciferase-expressing BPPNM cells were suspended in 200 µL PBS and the injected intraperitoneally in 7-9 week-old female C57BL/6 mice. To assess tumor burden, mice were intraperitoneally injected with 200 µL of 15 mg/kg D-luciferin, sodium salt (GoldBio, LUCNA), and luminescence measured after 15 minutes using an In Vivo Imaging System (IVIS) Spectrum (Perkin Elmer). Images were analyzed with Living Image Software. Luminescence is reported as total flux in photons per second [p/s].

### Pharmacokinetics, biodistribution, and cellular activation state analysis

At least 7 days prior to NP treatment, BPPNM tumor-bearing mice (described above) were placed onto AIN-93M Maintenance Purified Diet (TestDiet 1810541) to reduce fluorescence contribution from the chow. Prior to NP treatment, tumor bioluminescence signal was measured using IVIS and mice were subsequently divided into groups with comparable distributions of tumor burden. Hair on the abdomen region was removed using Nair as it attenuates fluorescence signal. Ten days after tumor cell inoculation, mice were injected intraperitoneally with various NP and poly(I:C) formulations to assess the biodistribution. For the biodistribution of bare liposomes and bilayer liposomes, mice were dosed with 200 μg BDP-650 labeled NPs in 200 μL. For the biodistribution of poly(I:C), mice were dosed with 50 or 150 μg poly(I:C) as free drug or LbL Poly(I:C) NP in 100-200 μL. All NPs and free poly(I:C) were dosed in sterile-filtered isotonic 5% dextrose. Mice treated with 5% dextrose were included as vehicle controls.

*Pharmacokinetics:* Mice were longitudinally imaged for fluorescence of BDP650-labeled NPs or Cy5-labeled poly(I:C) at multiple timepoints after dosing using an IVIS Spectrum (640 nm excitation filter, 680 nm emission filter). For the terminal timepoint, mice were injected with 200 μL of 15 mg/mL D-luciferin and imaged for NP or poly(I:C) fluorescence and then tumor bioluminescence. In Living Image software, ROIs were drawn around the peritoneal space to quantify fluorescence (total radiant efficiency in [p/s] / [µW/cm²]) in this space. To correct for background fluorescence, the average peritoneal fluorescence of the 5% dextrose vehicle treated mice was subtracted from each measurement.

*Organ-level biodistribution:* Organs of interest were removed and placed in cold RPMI-1640 media and kept on ice. Organs were imaged for BDP650-NP or poly(I:C)-Cy5 fluorescence then tumor bioluminescence using IVIS. Living Image Software was used to measure the bioluminescent total flux in [p/s], fluorescent total radiant efficiency in [p/s] / [µW/cm²], fluorescence average radiant efficiency in [p/s/cm²/sr] / [µW/cm²], and area in [cm²] of each organ. To correct for background fluorescence, the average fluorescence of the organ in the vehicle 5% dextrose treated mice was subtracted from each measurement. The fraction of recovered fluorescence was computed as the total radiant efficiency of the organ divided by sum of the total radiant efficiency for all organs recovered from an individual mouse.

*Cellular-level biodistribution and cell activation state:* At the designated timepoint, tumors and ascites were removed and processed into single cell suspensions using enzymatic digestion and manual dissociation (see details below). Single cells were then stained for various markers and cell populations and BDP-650 NP or Cy5-Poly(I:C) fluorescence assessed using flow cytometry (see details below). BDP-650 NPs and Cy5-Poly(I:C) were visualized in the APC channel.

### Whole-body cryo-fluorescence tomography

Mice were placed onto AIN-93M Maintenance Purified Diet 7 days prior to treatment and divided into groups with comparable distributions of tumor burden. For poly(I:C) distribution studies, BPPNM-luc tumor-bearing mice were dosed with 50 μg of poly(I:C)—as a free drug or LbL NP—of which 5% was Cy5 labeled. For comparison of PLD-layered, DXS-layered, and bare liposomes, mice with BPPNM-SIY-mCherry-luc tumors were dosed with 100 μg lipid of each formulation. Mice were euthanized by CO2 asphyxiation 24hr after dosing and immediately frozen by immersion in bath of hexanes and dry ice for 5 minutes. Frozen mice were stored at –20°C for at least 24 h to allow hexanes to evaporate and then embedded in optimum cutting temperature (OCT) compound. Mice were sectioned and imaged using an Xerra Cryofluorecence Tomography (CFT) imaging system (EMIT Imaging). The depth between sections was 55 μM. For each section, white light and fluorescence images were captured. Fluorescence images were captured with 470/511 nm excitation/emission to measure GFP expressed by BPPNM; 640/680 nm excitation/emission to measure BDP650 (liposomes) or Cy5 (poly(I:C)). Images were processed with ImageJ.

### Poly(I:C) serum pharmacokinetics

BPPNM-tumor bearing mice were dosed intraperitoneally with 50 μg free or NP-encapsulated poly(I:C). Cy5-labeled poly(I:C) was included at 5 wt/wt% of total poly(I:C) to enable quantification using fluorescence, and both the free and NP-encapsulated poly(I:C) were prepared from the same poly(I:C) stock solutions. At each timepoint after poly(I:C) dosing, blood samples were drawn via submandibular bleed and collected in serum collection tubes (Sarstedt #41.1500.005), centrifuged at 10,000xG for 5 minutes, and serum isolated and stored at –20°C until further use. Serum was thawed to room temperature, mixed well, then 20 μL undiluted serum added to well of 384-well black plate (Corning #3573) and read on plate reader for Cy5-poly(I:C) signal (645/675nm). Serum from dextrose-treated mice was used to subtract background fluorescence signal.

### Assessment of Acute Toxicity

*Single dose:* BPPNM-tumor bearing mice were intraperitoneally administered a single dose of poly(I:C) as free drug or encapsulated in LbL NP or lipid dose matched PLR/DXS bilayer Control NP in 5% dextrose 10 days after tumor inoculation. 24hr after treatment, blood was collected via submandibular bleeding into K3E EDTA Microtubes for complete blood counts (CBC) or Serum Gel CAT Microtubes for serum chemistry, and livers were fixed in 10% buffered formalin for downstream histological analysis. Blood analysis was performed by the Division of Comparative Medicine Comparative Pathology Laboratory at MIT, with CBCs analyzed using a HemaVet 950FS (Drew Scientific) and serum chemistry analyzed using a custom IDEXX panel. A separate cohort of mice was similarly treated, and their weights were monitored daily starting on the day of treatment to assess weight loss. 48hr after treatment, blood was collected for CBC and serum chemistry analysis.

*Daily dose:* BPPNM-tumor bearing mice were intraperitoneally administered 50 μg poly(I:C) as free drug or encapsulated in LbL NP in 5% dextrose daily for four days. Weights were monitored daily starting on the day of treatment to assess weight loss. 24hr and 48hr after the last treatment, blood was collected via submandibular bleeding into EDTA Microtubes for complete blood counts (CBC) per above.

### *In vivo* serum cytokine quantification

BPPNM-tumor bearing mice were dosed intraperitoneally with free or NP-encapsulated poly(I:C). At each timepoint after poly(I:C) dosing, blood samples were drawn via submandibular bleed and collected in Serum Gel CAT microtubes, centrifuged at 10,000xG for 5 minutes, and serum isolated and stored at –20°C until further use. Serum was thawed to room temperature, mixed well, then 20 μL undiluted serum added to well of 384-well black plate (Corning #3573) and read on plate reader for Cy5-poly(I:C) signal (645/675nm). Mouse TNF-α, CXCL10, and IL-6 levels were measured by ELISA, as described above using 1% BSA in PBS as the reagent diluent.

### Flow staining of tumor and ascites samples

Based on an average cell count, 10^6^ cells of each tumor sample and all of the ascites cells (typically 5×10^5^-10^6^ cells) were transferred to a 96-well V bottom plate for staining. Plate was centrifuged (500xg for 3 min at 4°C for this and subsequent steps), dumped, and cells resuspended in 50 μL Zombie Live/Dead stain (1:100) diluted in PBS and incubated for 10 min at room temperature. Cells were washed with 150 μL FACS buffer then Fc receptors were blocked with anti-CD16/32 (Biolegend #101339) at 1 μg per well in 50 μL for 15 min at 4°C. Cells were washed with 150 μL FACS buffer then stained with antibodies in 50 μL, diluted in FACS buffer and Brilliant Stain Buffer (BD #BDB563794) at specified dilutions listed in Supplementary information and Table S3, and incubated for 30 min at 4°C Cells were washed with 150 μL FACS buffer then fixed in 200 μL of 2% formaldehyde in PBS for 20 min at 4°C. Cells were centrifuged and resuspended in FACS buffer and stored at 4°C until analyzed. Compensation was calculated with UltraComp eBeads™ Plus Compensation Beads and single stained cells where appropriate. Flow cytometry was performed using a BD Symphony A3 equipped with HTS (BD Biosciences, New Jersey), and data analyzed in FlowJo V10.

### Efficacy

BPPNM-luc tumor-bearing mice were injected intraperitoneally with 2 mg/kg Dox encapsulated in LbL NP and/or 50 µg of poly(I:C) as free drug or encapsulated in NP in 5% dextrose control twice weekly for 2 weeks starting 10 days after tumor inoculation (doses on day 10, 13, 17, 20). 5% dextrose was used as the vehicle control. Mice were monitored and humanely euthanized when body condition score dropped below 2, weight loss exceeded 20%, or poor responsiveness was observed.

### Data Analysis and Statistics

All statistical tests were performed using GraphPad PRISM 10. Statistical significance was computed by ordinary one-way or two-way one-way analysis of variance (ANOVA) with Tukey or Bonferroni correction for multiple comparisons, as specified in figure captions. Results were considered significant at: **** = P < 0.001, *** = P < 0.001, ** = P < 0.01, * = P < 0.05, and ns = not significant. Data is presented as mean ± standard deviation (SD) or mean ± standard error of the mean (SEM), as specified in the figure caption.

## ACKNOWLEDGEMENT

This work was supported in part by the Koch Institute for Integrative Cancer Research Support (core) Grant 5P30-CA014051 from the National Cancer Institute. We thank the Koch Institute’s Robert A. Swanson (1969) Biotechnology Center for technical support, specifically the Flow Cytometry Core, Microscopy Core, The Peterson (1957) Nanotechnology Materials Core Facility, Preclinical Modeling, Imaging & Testing (PMIT) Core, and The Hope Babette Tang (1983) Histology Facility. We especially would like to thank DongSoo Yun for assistance with CryoTEM; Howard Mak and Anderson Scott for processing of Cryo-fluorescence tomography samples and data; Aurora Burds Connor, and Noranne Enzer for help with cell lines; and Kathy Cormier for histology preparation advice. We also would like to thank Grace Wolczanski and Stefani Spranger for providing luciferized BPPNM cells and the Department of Comparative Medicine at the Massachusetts Institute of Technology.

V.F.G. acknowledges support from the NSF Graduate Research Fellowship Program under Grant No. 2141064, MIT UCEM scholarship, and graduate fellowship from the Ludwig Center at MIT’s Koch Institute. Any opinions, findings, and conclusions or recommendations expressed in this material are those of the authors and do not necessarily reflect the views of the NSF. Fig. 1 was partially created using Biorender and ChemDraw. Fig. 2, 3, 7, and 8 were partially created using Biorender. Prism software was used for statistical analysis and data visualization. Finally, we are grateful to the entire Hammond lab for their advice and support, as well as Mickael Dang for review of this manuscript.

