## Supplementary information for "Engineering nanoparticle surface chemistry for antigen-presenting cell targeting improves specificity and safety of TLR3 agonist cancer immunotherapy"

### Additional methods

#### Preparation of BDP650-PC lipid

BDP650/665-DBCO, BDP TMR-DBCO, or ICG-DBCO dye (5mg/mL in chloroform) was combined with 1.2 equivalents of 18:0 azidoethyl PC (10 mg/mL and chloroform) in a glass vial and incubated overnight covered from light to allow for the conjugation of the dye to the lipid head group via strain promoted click chemistry[116]. The dye-lipid conjugates were assayed by thin-layer chromatography (TLC) on a silica plate with 65:25:4 (v/v/v) chloroform : methanol : water solvent to ensure absence of free dye. A stream of nitrogen was used to remove the chloroform from the reaction mixture. The dye-labeled lipids were then suspended in fresh chloroform at 0.5 mg/mL and stored at -20°C for subsequent use in liposome formulations.

#### Poly(I:C) reconstitution and dye-labeling

Low molecular weight Poly(I:C) (Invivogen #tlrl-picw) was reconstituted in the supplied endotoxin-free physiological water at 5 mg/mL, pipetted to homogenize, and subsequently annealed at 55°C for 15 minutes and allowed to cool to room temperature slowly. Poly(I:C) was aliquoted in DNA loBind tubes and stored at 20°C for long-term storage or at 4°C for short-term use. Prior to layering onto NPs, poly(I:C) stock solution was diluted to final concentration of 20 mM sodium acetate and approximately 10 mM sodium chloride which was remaining from the physiological water used to make the concentrated poly(I:C) stock solution.

For generation of Cy5 labeled poly(I:C), the Mirus Label IT® Nucleic Acid Labeling Kit, Cy5 (MIR 3700) was utilized at a labeling ratio of 50 μg poly(I:C) to 15 μL labeling reagent. In brief, poly(I:C), labeling buffer, and labeling reagent were combined in DNA loBind tubes and incubated at 37°C for 3 hr with occasional mixing. Labeled poly(I:C) was purified via ethanol precipitation. To the reaction mixture, 2.5X volumes of ice-cold ethanol and 0.1X volume of 3 M sodium acetate were added and subsequently incubated at -80°C for 1 hour to precipitate the poly(I:C). The solution was centrifuged at 30,000 rcf for 10 min at 4°C to pellet poly(I:C). Poly(I:C) was then washed 3X with 70% ethanol, dried to remove ethanol, and resuspended in milliQ water and stored at -20°C.

#### Quantification of poly(I:C) concentration

Cy5-labeled poly(I:C) was doped in at a wt/wt% of 0.25-5% relative to unlabeled poly(I:C), depending on the downstream application of the NPs, which enabled quantification of poly(I:C) concentration by measuring the fluorescence of the Cy5 dye (645/675nm). Samples and standards were diluted to 0.08 mg/mL 1000 kD PAA in water and fluorescence read in a Tecan microplate reader. The poly(I:C) concentration was calculated based on a calibration curve of the initial poly(I:C) solution of known concentration.

#### Quantification of doxorubicin loading

Encapsulated Dox concentration was quantified using fluorescence (470/560nm) based on a calibration curve of doxorubicin hydrochloride. Samples were diluted with 5 mg/mL poly(diallyldimethylammonium chloride) solution (PDADMAC) in water and DMSO at a ratio of 10:10:80 sample:PDADMAC:DMSO in black 96 well plates and read on a Tecan microplate reader, up to 12.5 μg/mL Dox to be within the linear fluorescence range. The polycation PDADMAC was included to displace Dox as the polyanions in the LbL NP formulations sequester and quench Dox which is positively charged. DMSO was also investigated as the dissociation agent but the Dox fluorescence standard curve had a broader linear range in methanol compared to DMSO. This method was validated against HPLC quantification and was preferred as it reduces sample preparation and quantification time and necessary volumes of organic solutions.

**Cell culture**

RAW264.7 cells were purchased from ATCC. HUVEC cells were purchased from Lonza. A2780 cells were a gift from the laboratory of Stephen Howell (UC San Diego). Kuramochi-mCherry-luc cells were a gift from the laboratory of Ronny Drapkin (University of Pennsylvania). OVCAR8 cells were a gift from the laboratory of Sangeeta Bhatia (MIT). DC.4 cells were a gift from the laboratory of Kenneth Rock (University of Massachusetts Medical School). NIH3T3, HepG2, BNL CL.2, Hepa 1-6, and THP-1 cells were obtained from the Koch Institute cell repository. HEK-Blue mTLR4 (hkb-mtlr4) reporter cells were purchased from InvivoGen. Luciferase-expressing BPPNM and KPCA.C cells and BPPNM cells expressing SIY-mCherry-luciferase were a gift from the laboratory of Stefani Spranger (MIT). Parental BPPNM and KPCA.C cells were a gift from the laboratory of Robert Weinberg (MIT)[64] and were transduced with a plasmid containing firefly luciferase and a blasticidin resistance gene using lentivirus, then selected for using blasticidin (15 µg/mL). Media conditions for each cell line are listed in SI Table S4. Cell lines were authenticated using STR profiling (ATCC) and periodically tested for mycoplasma with a MycoAlert kit (Lonza). Cells were cultured at 37°C in humidified 5% $CO_2$. Live cells were counted using Trypan Blue stain with a Cellometer Auto 1000 (Nexcelom).

**Bone marrow derived dendritic cell (BMDC) differentiation**

BMDCs were generated as follows, adapted from Zagorulya et al[124]. Bone marrow was extracted from the hind limbs of 7-10 week old C57BL/6 mice via centrifugation through a 70 μm strainer (Pluriselect, Cat# 431007060). After ACK buffer lysis, cells were cultured in RPMI-1640 media (Gibco) supplemented with 10% FBS, 1% Pen/Strep, 55μM β-mercaptoethanol (Gibco, Cat# 21985023), 25 mM HEPES (Gibco, Cat# 15630080), 1X non-essential amino acids (Gibco Cat# 11140050), 1 mM Sodium pyruvate (Gibco Cat# 11360070), 5 ng/ml murine GM-CSF (BioLegend, Cat# 576304) and 100 ng/mL Flt-3L-Ig (BioXCell, Cat# BE0098) in 10 cm non-tissue culture treated plates (VWR, #10861-594) at $2 \times 10^6$ cells/ml in 10 mL at 37°C and 5% CO2. On day 4, 4 mL of BMDC media was added to the cells. After 6 days of culture, non-adherent and semi-adherent cells were collected as the BMDC fraction and frozen in 10% DMSO (Fisher, Cat# BP231-100) / 90% FBS and stored in liquid nitrogen for later use. For assays using BMDCs, cells were seeded in non-tissue culture treated, sterile V-bottom plates (Thermo, #249662).

**Magnetic Cell Separation (MACS) of BMDCs**
*Composition of MACS buffer:* MACS buffer was composed of sterile 5% FBS in PBS (w/o $Ca^{2+}$ or $Mg^{2+}$), 2 mM EDTA. The solution was filtered through a 0.2 μm syringe filter and degassed before use.

*Separation of BMDCs:* An enriched population of BMDCs was obtained using the PanDC Isolation kit (Miltenyi, Cat# 130-100-875) according to the manufacturer's protocol, which depletes non-target (non-DC) cells. Throughout the process, cells are kept cool and steps done as quickly as possible to improve the yield of viable cells. In brief, frozen BMDCs were thawed, centrifuged, and resuspended in 350 μL MACS buffer. To the cells, 50 μL of FcR Blocking Reagent and 100 μL of Pan Dendritic Cell Biotin-Antibody Cocktail was added and mixed well. Cells were incubated for 10 min at 4°C. To wash, 5 mL of MACS buffer was added and cells were centrifuged at 500 x g for 3 min to pellet cells. Supernatant was removed completely and cells were resuspended in 800 μL MACS buffer followed by the addition of 200 μL of anti-biotin microbeads, mixed well, and incubated for 10 min at 4°C. BMDCs were then enriched via magnetic separation: cells were added to an LS column (Cat# 130-042-401) in a QuadroMACS™ Separator (Cat# 130-090-976) attached to a MACS magnetic stand (# 130-042-303) and the column washed twice with 3 mL of MACS buffer. All eluents were collected, combined, and centrifuged to obtain enriched BMDCs. Cells were resuspended in full BMDC media and counted.

**Microscopy of BMDCs to assess NP localization**
*Preparation of chamber slides:* To enable adherence of BMDCs, 8-chamber slides (Lab-Tek, Cat #155409) were coated with 0.01% poly-L-lysine (Sigma, Cat #RNBK2055) at room-temperature for 15 min. Poly-L-lysine solution was removed and chamber slides were ready to be seeded with BMDCs.

*Preparation of cells:* For fixed cell imaging of NP localization in BMDCs, frozen BMDCs were thawed, subjected to MACS to enrich for a purer DC population, and then resuspended in full BMDC media. BMDCs were plated at 50,000 cells in 90 mL full BMDC media and treated with 10 μL NPs or poly(I:C) in a 96 well non-TC treated V-bottom plate. After 4hr, DCs were centrifuged and rinsed with PBS and then two replicates per condition were combined into one chamber slide well to achieve 100k cells/well. Cells were incubated for 15 min at RT to allow cells to adhere and fixed in 300 mL 4% formaldehyde (Thermo, Cat# 28906) in PBS at RT for 15 min, protected from light. Cells were subsequently stained per below.

*Cell staining without antibody staining*: Fixed cells were washed twice with PBS then stained with 10 μg/mL wheat germ agglutinin-AF488 (Invitrogen, Cat# W11261) or wheat germ agglutinin-AF555 (#W32464) and 1 ug/mL Hoechst (Invitrogen, #H3570) in HBSS (Gibco, Cat# 14175-095) in 300 μL for 10 min at RT. Cells were gently washed twice with PBS then fixed in 4% formaldehyde for 2 min at RT. Cells were stored in PBS at 4°C, covered from light, until ready to image.

**Microscopy of RAW264.7 to assess NP localization and trafficking**

*Preparation of chamber slides:* To improve adherence of cells, 8-chamber slides were coated with rat tail collagen (Sigma #08-115), 300 µL of 50 µg/mL in 0.02 N acetic acid per well at room-temperature for 1 hr. Collagen solution was removed and chamber slides were rinsed with PBS, and were then ready to be seeded with cells.

*Preparation of cells:* RAW264.7 cells were seeded at 25,000 cells/well in 270 µL media and allowed to adhere overnight. Cells were treated with 30 µL of treatment. At the end of the incubation period, cells were rinsed with PBS and fixed in 300 µL 4% formaldehyde in PBS at RT for 15 min, protected from light. Cells were subsequently stained per below.

*Cell staining with intracellular antibodies*: Fixed cells were washed twice with PBS then stained with 5 µg/mL wheat germ agglutinin-AF488 or wheat germ agglutinin-AF555 in 300 µL for 10 min at RT. Cells were gently washed twice with PBS then fixed in 4% formaldehyde for 2 min at RT. Cells were treated with blocked with 0.025% saponin and 5% goat serum in PBS for 1 hr at RT. Blocking buffer was removed and primary antibodies diluted into 300 µL antibody diluent buffer (0.025% saponin and 1% BSA in PBS) were added. Cells were incubated in the primary antibody overnight at 4 °C, then washed three times with PBS. Fluorophore conjugated secondary antibodies or DyLight 554 phalloidin (1:400 dilution) in 0.025% saponin and 1% BSA in PBS were added to each well and incubated for 1 hr at RT. Cells were washed three times with PBS, fixed again in 1% formaldehyde for 5 min at RT, and then washed with PBS. Cells were stained with 1 ug/mL Hoechst in 300 µL for 5 min at RT, washed three times in PBS and stored in PBS at 4°C, covered from light, until ready to image. Antibodies and dilutions can be found in SI.

**Microscopy Imaging and data processing**

Before imaging BMDCs or RAW264.7 cells, PBS was removed and Vectashield Antifade Mounting Medium (#H-1000) added to cover each well. Cells were imaged with a confocal laser-scanning microscope—Olympus FV1200 equipped with 405, 473, 559, and 635 nm lasers or Evident FV4000 equipped with 405, 488, 561, 640 nm, and 730 lasers. Images were acquired with 60x or 100x objectives and processed using ImageJ software.

**$IC_{50}$ Curves**

BPPNM-luc cells (2.5k/well) were seeded in black-walled, clear-bottom TC-treated 96 well plate (Thermo #165305) and allowed to adhere overnight. Cells were dosed with various concentrations of free drug or NPs diluted in water (10% of the well volume). At the designated timepoints, the viability was assessed using the PrestoBlue HS resazurin-based assay (Invitrogen, #P50201) following the manufacturer instructions. Briefly, 10 µL of PrestoBlue HS reagent was directly added to each well of cells and incubated for 45 min at 37°C, 95% humidity, and 5% $CO_2$. The fluorescence of the wells was then read (560/590 nm) using a Tecan Infinite MPlex Plate Reader. Data was analyzed after subtracting the fluorescence of the blank (PrestoBlue HS incubated in media with no cells), and reported relative to the water treated control. Three technical replicates were used per group for each biological replicate.

**Flow cytometry staining**

For in vivo cellular biodistribution studies (Fig. 2, S11-13), cells were stained with Zombie UV (1:100), CD45-BUV395 (1:40), MHC-II-BUV805 (1:60), CD11c-BV421 (1:20); F4/80-BV785 (1:20), CD11b-PE-Dazzle594 (1:400), Gr-1-BUV661 (1:320), CD19-BV711 (1:80), CD3e-PE (1:20), NK1.1-APC-Cy7 (1:40).

For in vivo poly(I:C) delivery to and activation of antigen presenting cells (Fig. 5, S31-33), cells were stained with Zombie UV (1:100), CD45-BUV395 (1:40), MHC-II-BUV805 (1:60), CD11c-BV421 (1:20); F4/80-BV785 (1:20), CD11b-PE-Dazzle594 (1:400), Gr-1-BUV661 (1:320-1:400), CD206-BV605 (1:20-1:50), CD86-PE-Cy7 (1:20), CD19-APC-Cy7 (1:40), CD3e-APC-Cy7 (1:40), NK1.1-APC-Cy7 (1:40).

For ex vivo splenocyte NP association (Fig. S12), cells were stained with Zombie NIR (1:500), CD45-BUV395 (1:400), CD11b-PE594 (1:500), CD3e-PE (1:200), CD19-BV711 (1:400)

For poly(I:C) delivery and activation of RAW264.7 macrophages (Fig. 4a, b, e; S21), cells were stained with Zombie Violet (1:500), CD206-BV786 (1:400), MHC-II-PE-Cy7 (1:400), CD86-BV605 (1:300).

For poly(I:C) delivery and activation of BMDCs, cells were stained with the following corresponding to different figures, details of analyzed parameters, and cytometer used:

Fig. 4a-d, f; S12: Zombie UV (1:500), CD11c-BV421 (1:400), MHCII-BUV805 (1:400), CD40-PE (1:200), CD80-PE-Cy7 (1:200), CD86-BV605 (1:200);

Fig. 4g; S13a-g: Zombie Aqua (1:500), CD11c-BV421 (1:400), MHC-II-BUV805 (1:400), F4/80-PE (1:400), CD86-PE-Cy7 (1:300), CD3e-APC-Cy7 (1:200), CD19-APC-Cy7 (1:400), NK1.1-APC-Cy7 (1:400)

Fig. S13h-n: Zombie UV (1:500), CD11c-BV421 (1:400), MHC-II-BUV805 (1:400), CD86-BV605 (1:200), CD3e-APC-Cy7 (1:200), CD19-APC-Cy7 (1:400), NK1.1-APC-Cy7 (1:400)

Fig. S16: Zombie UV (1:500), CD11c-BV421 (1:400), MHC-II-BUV805 (1:400), CD40-PE (1:200), CD86-BV605 (1:200)

**Digestion of tumor and ascites for single cell suspension**

*Preparation of enzyme digestion solution:* Digestion mixture was prepared from 10X stocks in RPMI to a final concentration of 1 mg/mL collagenase type V, 0.25 mg/mL hyaluronidase, 0.1 mg/mL DNAse I and kept on ice until use. The 10X stocks were stored at -20°C.

*Tumor digestion:* Tumor was collected and stored in RPMI media on ice. To the gentleMACS™ C Tubes (Miltenyi #130-096-334), 2.5 mL digestion mixture and tumor were added. Tumor was minced using forceps and kept on ice. Tumors were digested using a gentleMACS Octo Dissociator (Miltenyi), m_imp-tumor-02 program. Tubes were incubated under shaking (150 rpm) at 37°C for 30 min. Tubes were returned to ice and digested using m_impTumor_03 program. Digested tumors were diluted 1:1 with FACS buffer (1% BSA and 2 mM EDTA) to final concentration of 1 mM EDTA and 0.5% BSA as EDTA quenches the enzymatic reaction. Dissociated tumors were filtered through a 70 µm filter (Miltenyi #130-110-916) into new 15mL tube to obtain a single cell suspension and filter was rinsed with 2 mL FACS buffer to complete the transfer. Cells were collected by spinning 15 mL tubes at 300xg for 5 min at 4°C. Supernatant was decanted and 2 mL ACK lysis buffer was added, tube gently vortexed to resuspend cells and incubated for 2 min to lyse red blood cells. To quench the ACK, 5 mL FACS buffer was added and the tube spun at 300xg for 5 min. This process was repeated until a white cell pellet was obtained. Supernatant was removed and cells resuspended in 200 µL of FACS buffer, counted, and left on ice until flow staining.

*Ascites cells collection and digestion*: Using a 3 mL syringe attached with an 18G needle (Air-Tite #ML3181), cold RPMI (2.5 mL) was injected into the peritoneal cavity. The peritoneal cavity was gently massaged and then the fluid was collected in the syringe and transferred to 15 mL tube and kept on ice. Tubes were centrifuged at 300g for 5 min at 4°C, supernatant decanted, and cells resuspended in 1 mL of enzyme digestion mixture by gentle vortex. For the digestion, cells were then incubated at 37°C for 30 min on an orbital shaker incubator under agitation (150 rpm). Following enzymatic digestion, 1 mL of FACS buffer was added to quench the enzymatic reaction. Cells were centrifuged at 300g for 5 min at 4°C. Supernatant was decanted and 2 mL ACK lysis buffer was added, tube gently vortexed to resuspend cells and incubated for 2 min to lyse red blood cells. To quench the ACK, 5 mL FACS buffer was added and the tube spun at 300xg for 5 min. Cells were resuspended in 200 µL FACS buffer by vigorous pipetting and passed through a 70 µm filter into a fresh tube to obtain a single cell suspension. Cells were counted and left on ice until flow staining.

**Table S1. Liposome formulations.**

| | DSPC:Chol:DSPG:Dye-lipid | |
|---|---|---|
| **Liposome formulation** | **Molar ratio** | **Mass ratio** |
| Empty (1% NBD) | 40.8 : 32.1 : 24.7 : 0.7 | 49 : 20 : 30 : 1 |
| Empty (0.25% NBD) | 41 : 34 : 25 : 0.25 | 49 : 20 : 30.4 : 0.4 |
| Empty (0.2% BDP650, ICG, or BDP-TMR) | 40.8 : 34 : 25.5 : 0.2 | 49 : 20 : 30 : 0.5 |
| | **POPC:Chol:POPG** | |
| Doxorubicin loaded | 49:45:6 | 60:28:7.5 |

**Table S2. Primary and secondary antibodies for microscopy**

| Antibody | Clone | Product # | Dilution |
|---|---|---|---|
| Primary Antibodies | | | |
| EEA1 Rabbit mAb | C45B10 | CST #3288S | 1:200 |
| LAMP1 Rabbit mAb | E5N9Z | CST #99437S | 1:400 |
| Mouse TLR3 Antibody Rat IgG2A mAb | 313129 | RD #MAB3005 | 1 μg/mL |
| RIG-I Recombinant Rabbit mAb | 35H2L48 | Thermo #PI700366 | 1:400 |
| Secondary Antibodies | | | |
| Anti-rat IgG (H+L) Alexa Fluor 555 Conjugate | | CST #4416S | 1:1000 |
| Anti-rat IgG (H+L) Alexa Fluor 488 Conjugate | | CST #4417S | 1:1000 |
| F(ab')2-Goat anti-Rabbit IgG (H+L) Alexa Fluor 488 Conjugate | | Thermo #A11070 | 1:1000 |
| F(ab')2-Goat anti-Rabbit IgG (H+L) Alexa Fluor 546 Conjugate | | Thermo #A11071 | 1:1000 |

**Table S3. Antibodies and stains for flow cytometry**

| Antibody | Fluorophore | Clone | Product # |
|---|---|---|---|
| anti-mouse CD16/32 | | 93 | BioLegend #101339 |
| CD45 | BUV 395 | 30-F11 | BD Biosciences #564279 |
| MHC-II (I-A/I-E) | BUV805 | M5/114.15.2 | BD Biosciences #748844 |
| MHC-II (I-A/I-E) | PE-Cy7 | M5/114.15.2 | BioLegend #107630 |
| CD11c | BV421 | N418 | BioLegend #117330 |
| F4/80 | PE | BM8 | BioLegend #123110 |
| F4/80 | BV785 | BM8 | BioLegend #123141 |
| Ly6C/Ly6G (Gr-1) | BUV 661 | RB6-8C5 | BD BioSciences #741470 |
| Ly6C | PE | HK1.4 | BioLegend #128007 |
| CD11b | PE-Dazzle 594 | M1/70 | BioLegend #101256 |
| CD8a | BV605 | 53-6.7 | BioLegend #100744 |
| CD103 | BV711 | 2E7 | BioLegend #121435 |
| CD206 | BV786 | C068C2 | BioLegend #141729 |
| CD206 | BV605 | C068C2 | BioLegend #141721 |
| CD40 | PE | 3/23 | BioLegend #124610 |
| CD80 | PE-Cy7 | 16-10A1 | BioLegend #104734 |
| CD86 | BV605 | GL-1 | BioLegend #105038 |
| CD86 | PE-Cy7 | GL-1 | BioLegend #105014 |
| CD3e | PE | 17A2 | BioLegend #100206 |
| CD3e | APC-Cy7 | 17A2 | BioLegend #100222 |
| CD19 | APC-Cy7 | 6D5 | BioLegend #115530 |
| CD19 | BV711 | 6D5 | BioLegend #115555 |
| NK1.1 | APC-Cy7 | PK136 | BioLegend #108724 |
| Zombie UV | | | BioLegend #423107 |
| Zombie Violet | | | BioLegend #423113 |
| Zombie Aqua | | | BioLegend #423101 |
| Zombie NIR | | | BioLegend #423105 |

**Table S4. Layering conditions of LbL NP library.** Liposomes labeled with 1 mol% NBD (compliant formulation) layered at 0.25 mg/mL final lipid concentration at the weight (wt) ratio of liposomal core to polymer listed and final buffer concentrations listed. All LbL NP formulations were purified using tangential flow filtration to remove excess polymer and buffer exchange into water.

| | Layering conditions | |
|---|---|---|
| Layer | Wt ratio (core:polymer) | Final buffer concentration |
| PLR | 1:0.3 | 25 mM HEPES, 20 mM NaCl |
| PLE-PEG | 1:0.3 | 25 mM HEPES, 20 mM NaCl |
| PLE | 1:0.5 | 25 mM HEPES, 20 mM NaCl |
| PLD | 1:0.3 | 25 mM HEPES, 20 mM NaCl |
| DXS | 1:0.4 | 25 mM HEPES, 20 mM NaCl |
| FUC | 1:0.5 | 25 mM HEPES, 20 mM NaCl |
| CSA | 1:0.5 | 25 mM HEPES, 20 mM NaCl |
| HA | 1:0.6 | 5 mM HEPES |
| CMDX | 1:0.6 | 5 mM HEPES |
| ALG | 1:0.5 | 5 mM HEPES |
| PAA | 1:0.3 | 25 mM HEPES, 20 mM NaCl |

**Table S5. Cell culture conditions and seeding densities for 96 well plate for NP association flow cytometry assays.** HUVEC – human umbilical vein endothelial cells; HGSC – high grade serous carcinoma; FTE – fallopian tube epithelial

| | | Cell culture conditions | | | |
|---|---|---|---|---|---|
| **Cell line** | **Cell type** | **Media** | **4 hr seeding density** | **24 hr seeding density** | **Special notes** |
| Human cell lines: | | | | | |
| A2780 | Ovarian endometrioid adenocarcinoma | RPMI, 10% FBS, 1% P/S | 40k | 25k | |
| Hep G2 | Hepatocellular carcinoma | DMEM, 10% FBS, 1% P/S | 40k | 25k | |
| HUVEC | Endothelial cells | EBM-2, EGM-2 supplements | 20k | 10k | EGM®-2 Bulletkit (Lonza #CC-3162). Cultured on flat bottom Delta treated plates |
| Kuramochi -mCherry | HGSC | DMEM:F12 1:1, 10% FBS, 1% P/S | 25k | 10k | |
| OVCAR8 | HGSC | RPMI, 10% FBS, 1% P/S | 25k | 10k | |
| THP-1 | Monocyte | ATCC-mod RPMI, 10% FBS, 1% P/S | 50k | 30k | ATCC-mod RPMI (2 mM L-glutamine, 10 mM HEPES, 1 mM sodium pyruvate, 4500 mg/L glucose, and 1500 mg/L sodium bicarbonate) (ATCC, #30-2001) |
| Mouse cell lines: | | | | | |
| BNL CL.2 | Normal liver | DMEM, 10% FBS, 1% P/S | 15k | 10k | |
| BPPNM-luc | HGSC model from FTE cells | DMEM-high glucose & HEPES, 4% heat-inactivated FBS, 1% P/S, ITS-G 1X | 25k | 10k | DMEM with high glucose & HEPES (Gibco #12430062), ITS-G (Gibco #41-400-045), heat-inactivated FBS (Sigma #F4135) |
| DC2.4 | Dendritic cell | RPMI, 10% FBS, 1% P/S | 40k | 25k | |
| Hepa 1-6 | Hepatoma | DMEM, 10% FBS, 1% P/S | 25k | 10k | |
| HM-1 | Ovarian cancer | MEM Alpha, 10% FBS, 1% P/S | 25k | 15k | |
| NIH3T3 | Fibroblast | DMEM, 10% FBS, 1% P/S | 10k | 5k | |
| RAW264.7 | Macrophage | DMEM, 10% FBS, 1% P/S | 40k | 25k | |

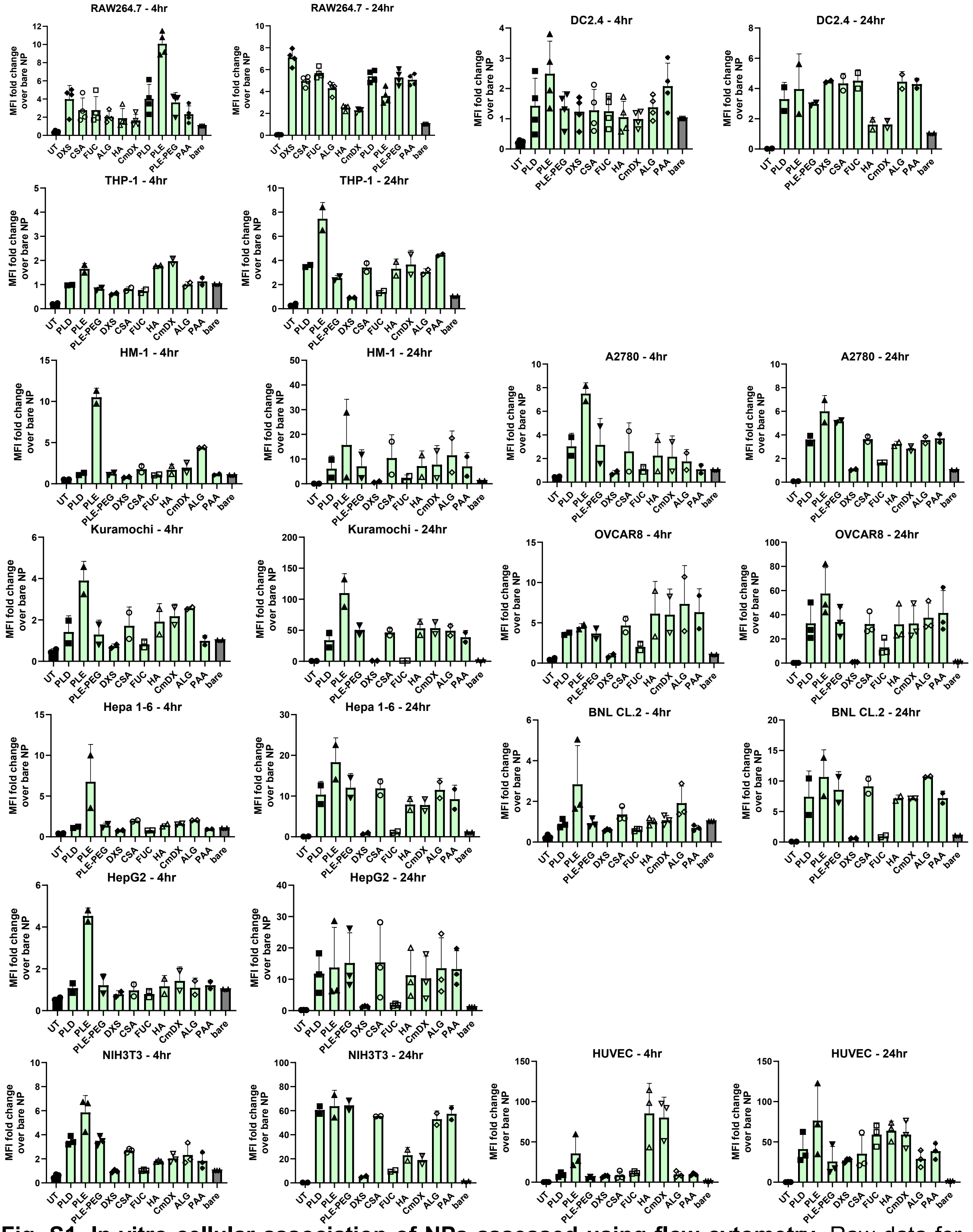


**Fig. S1. In vitro cellular association of NPs assessed using flow cytometry.** Raw data for Fig. 1d-e. Fold change of the median fluorescence intensity (MFI) of layered NBD-labeled NPs over the MFI of the bare, unlayered liposome after 4 hr and 24 hr incubation with cells. NPs dosed at 25 μg/mL per well in triplicate. Data points represent 2-3 biological replicates averaged from 3 technical replicates. Error bars represent the standard deviation from the mean of the biological replicates.

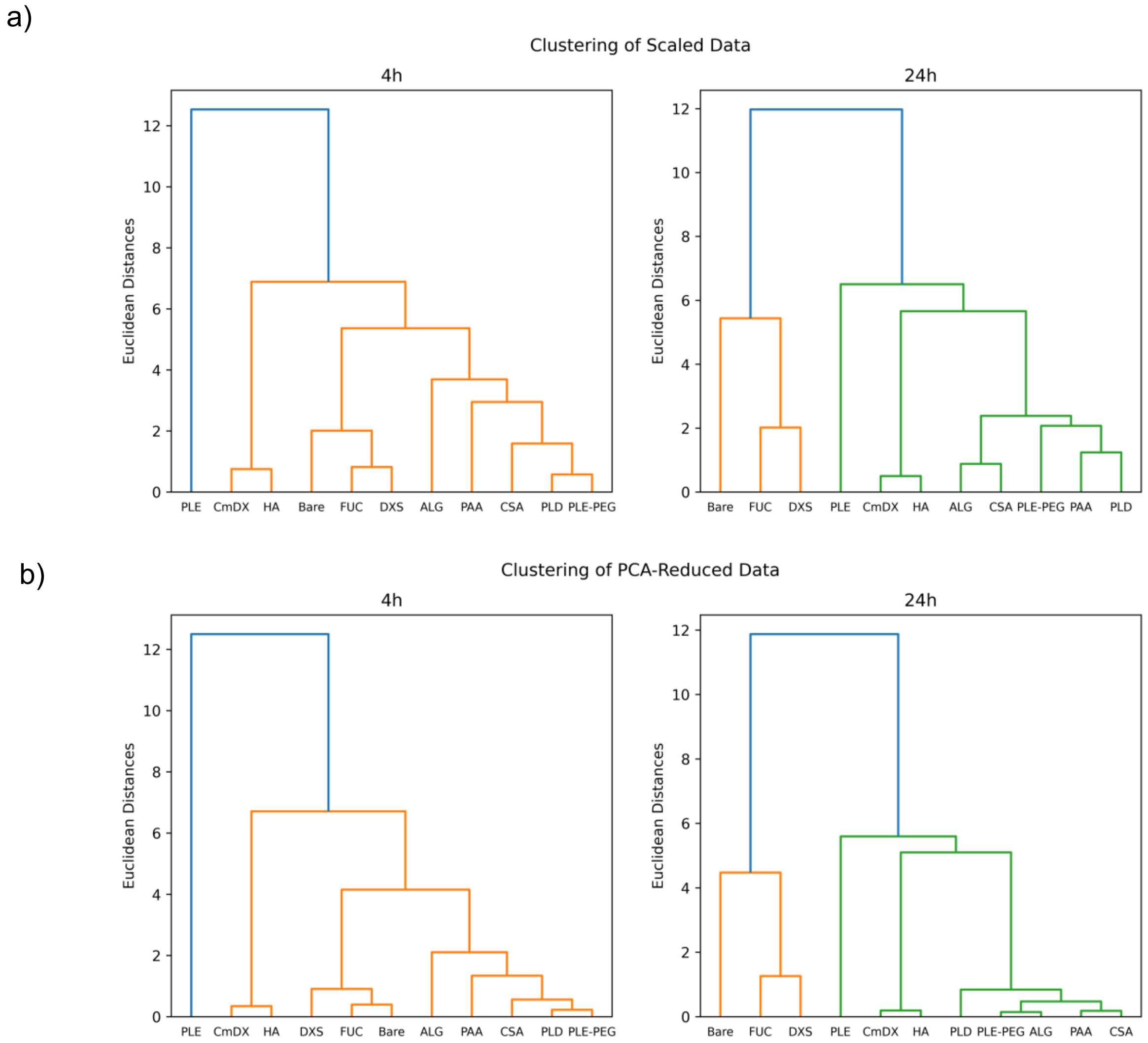


**Fig. S2. Hierarchical clustering analysis of NP surface chemistry cellular association.** Clustering analysis of Z-scored NP association (median fluorescence intensity relative to that of the bare liposomes for each cell line) in Fig. 1-S1 using a) Ward clustering on the standard scaled data—applied to Fig. 1d-e and b) Ward clustering on the principal component analysis transformed data.

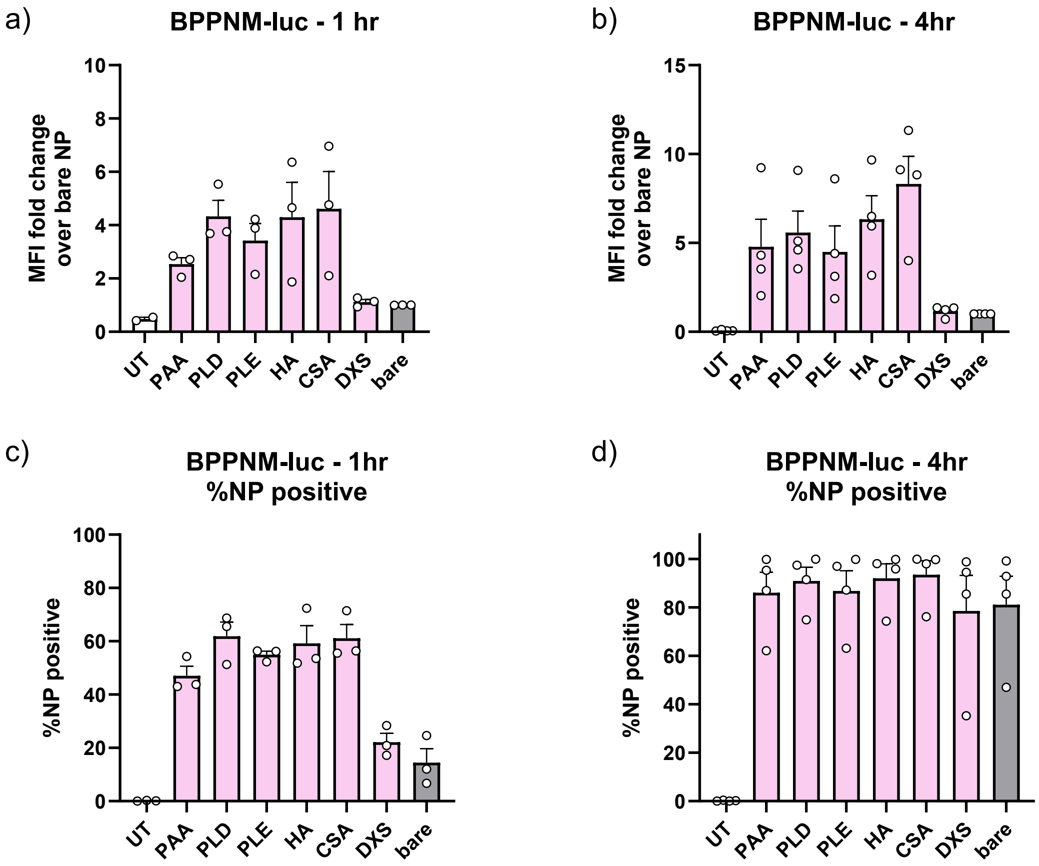


**Fig. S3. In vitro cellular association of NPs with BPPNM-luc assessed using flow cytometry.** Fold change of the median fluorescence intensity (MFI) of layered BDP-TMR-labeled NPs over the MFI of the bare, unlayered liposome after a) 1 hr and b) 4 hr incubation with BPPNM-luc ovarian cancer cells. Percentage of live cells that are positive for NPs after c) 1 hr and d) 4 hr. NPs dosed at 15 μg/mL per well in triplicate. Data points represent 3-4 biological replicates averaged from 3 technical replicates. Error bars represent the standard deviation from the mean of the biological replicates. BPPNM-luc cells express GFP so NBD-labeled NPs used in the previous library were incompatible, necessitating the use of a different fluorophore to label the NPs which is not in the green channel.

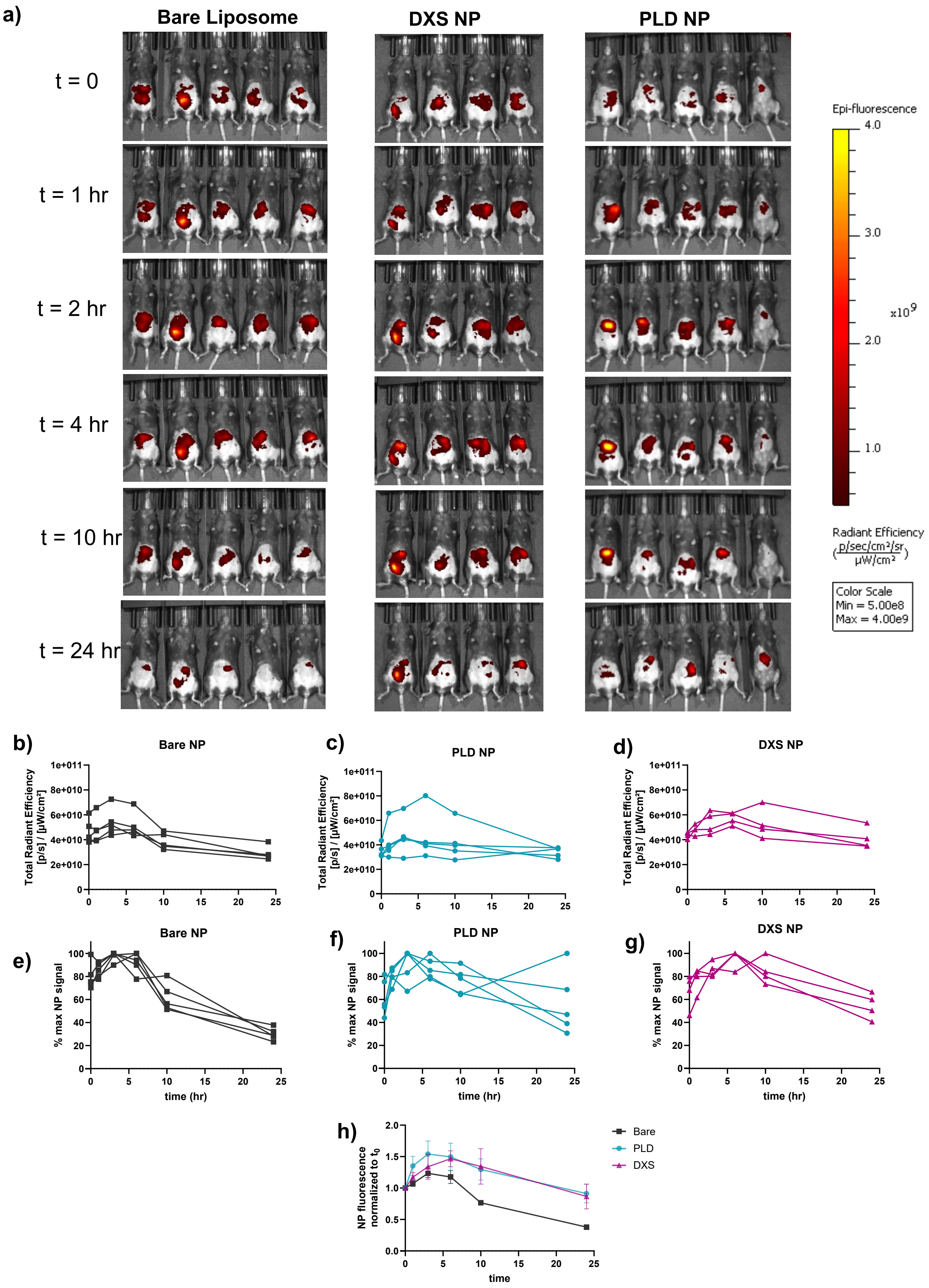


**Fig. S4. Pharmacokinetics of bare and LbL-coated liposomes.** Female C57BL/6 mice with BPPNM-luc tumors (3 million, IP) were injected with 200 µg of each NP formulation, labeled with BDP650, in 5% dextrose 10 days post tumor inoculation (N=4-5). Mice were maintained on alfafa-free diet to minimize fluorescence signal from the chow. a) Mice were imaged using an in vivo imaging system (IVIS) for BDP650 NP fluorescence at various timepoints. b-d) Raw values of NP fluorescence (total radiant efficiency) in peritoneal space for each mouse. e-g) NP signal in peritoneal space normalized to the maximum NP signal for each mouse. h) Average of the individually normalized NP signal in peritoneal space relative to NP signal at t=0.

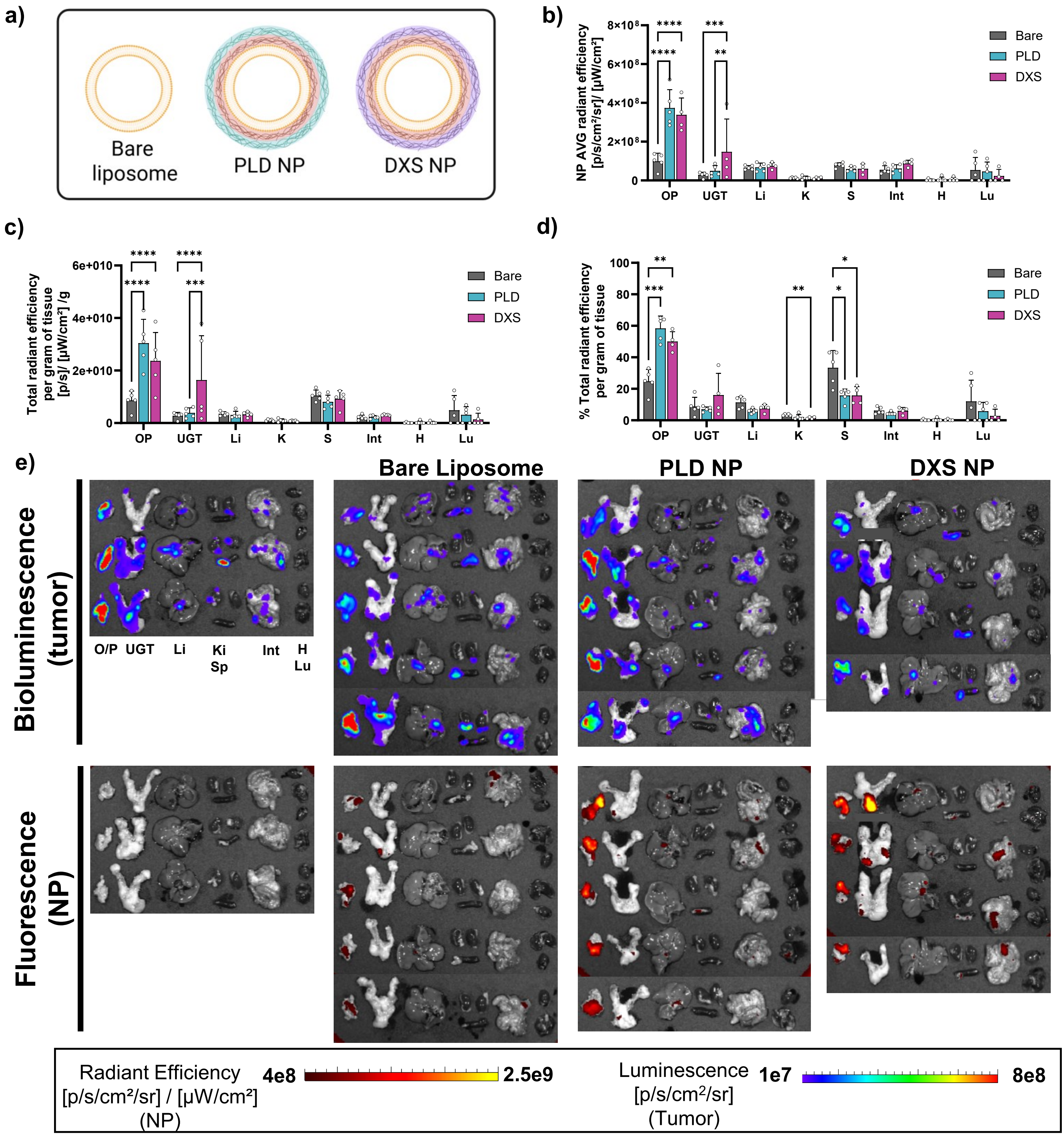


**Fig. S5. In vivo biodistribution of bare and LbL NPs upon intraperitoneal administration.** a) Female C57BL/6 mice with BPPNM-luc tumors were injected with 200 µg of each NP formulation, labeled with BDP650, in 5% dextrose 10 days post tumor inoculation (N=4-5). 24 hrs after NP injection, organs were removed and imaged for BDP650 NP fluorescence and tumor bioluminescence. Mice injected with 5% dextrose (vehicle control) were used to subtract background organ signal. NP fluorescence in each organ determined by b) average radiant efficiency—corresponding to a size normalization and c) total radiant efficiency normalized by organ weight. d) % recovered total radiant efficiency normalized by weight(total radiant efficiency in organ normalized by organ weight then divided by the sum of recovered average radiant efficiency—divided by weight of each organ—across all organs). e) Images of NP fluorescence (radiant efficiency) and tumor bioluminescence (total flux). Statistically significant comparisons from respective 2-way ANOVAs with Tukey multiple comparisons test. PLD – poly-L-aspartate; DXS – dextran sulfate; O/P – omentum/pancreas; UGT – upper genital tract; Li – liver; Ki – kidney; Sp – spleen; Int – intestines; H – heart; Lu – lungs.

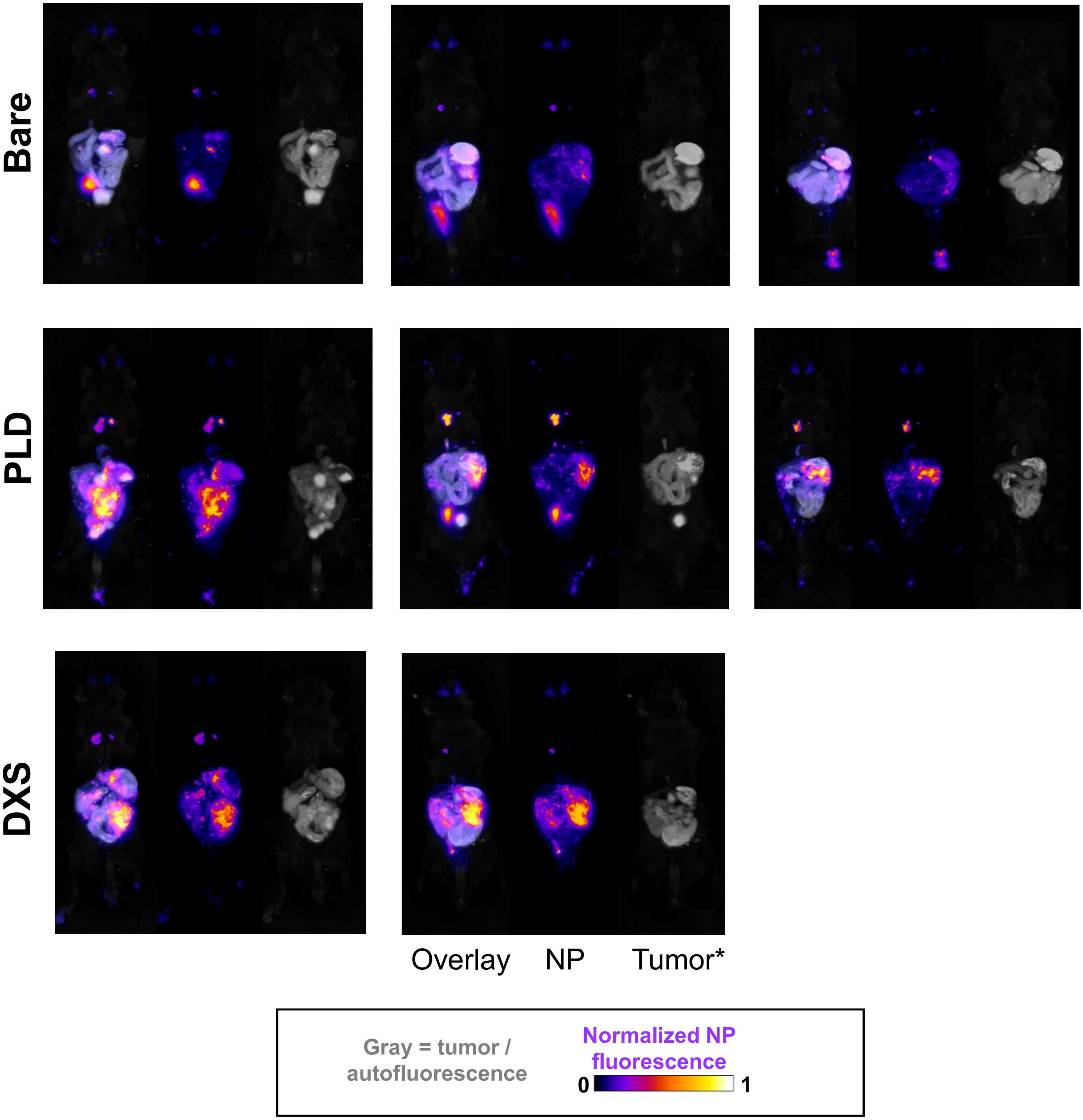


**Fig. S6. Distribution of bare and LbL NPs upon intraperitoneal administration using cryo-fluorescence tomography.** Female C57BL/6 mice with BPPNM-luc tumors (3 million, IP) were injected with 200 μg of each NP formulation, labeled with BDP650, in 5% dextrose 10 days post tumor inoculation. 24 hrs after NP injection, mice were humanely euthanized and prepared for cryo-fluorescence tomography to visualize the fluorescence of the BDP650 of the NPs and GFP from the BPPNM tumors throughout the mouse. Tissue autofluorescence is also detected in the GFP channel. Shown are 3D projections of max intensity for the overlay of NP fluorescence and BPPNM-GFP / autofluorescence as well as the single channel images. Mice with obvious fatpad injections were excluded from analysis. PLD – poly-L-aspartate; DXS – dextran sulfate.

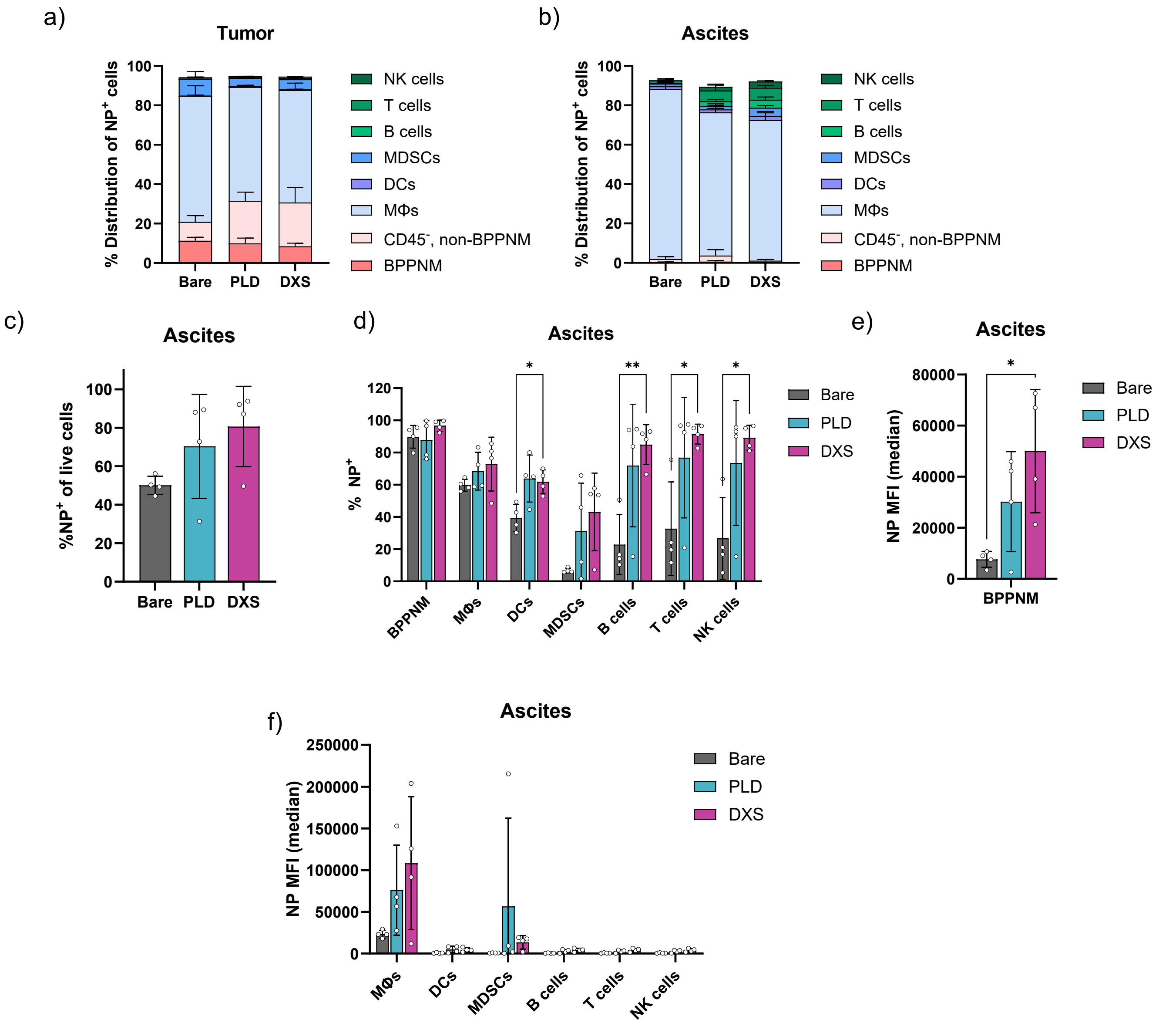


**Fig. S7. Cellular NP distribution in tumor and ascites.** Female C57BL/6 mice with BPPNM-luc tumors (3 million, IP) were injected with 200 μg of BDP650-tagged bare liposomes, PLD LbL NPs, or DXS LbL NPs in 5% dextrose 10 days post tumor inoculation (N=4-5). Ascites and tumors were collected 24 hrs after NP injection, digested, and subjected to flow cytometry analysis to assess cellular NP distribution. Gating strategy described in **Fig. S8**. Distribution of population of cells that are NP⁺ in the a) tumor and b) ascites. c) Percentage of live cells in the ascites that are positive for NP signal. d) Percentage of each cell type in the ascites that are positive for NP signal. Median fluorescence intensity (MFI) of NP in e) BPPNM and f) immune cells in the ascites. Statistically significant comparisons determined using 2-way ANOVA in d, f) and 1-way ANOVA in c, e) with Tukey multiple comparisons test. PLD – poly-L-aspartate, DXS – dextran sulfate, MΦs – macrophages, DCs – dendritic cells, MDSCs – myeloid-derived suppressor cells, NK – natural killer.

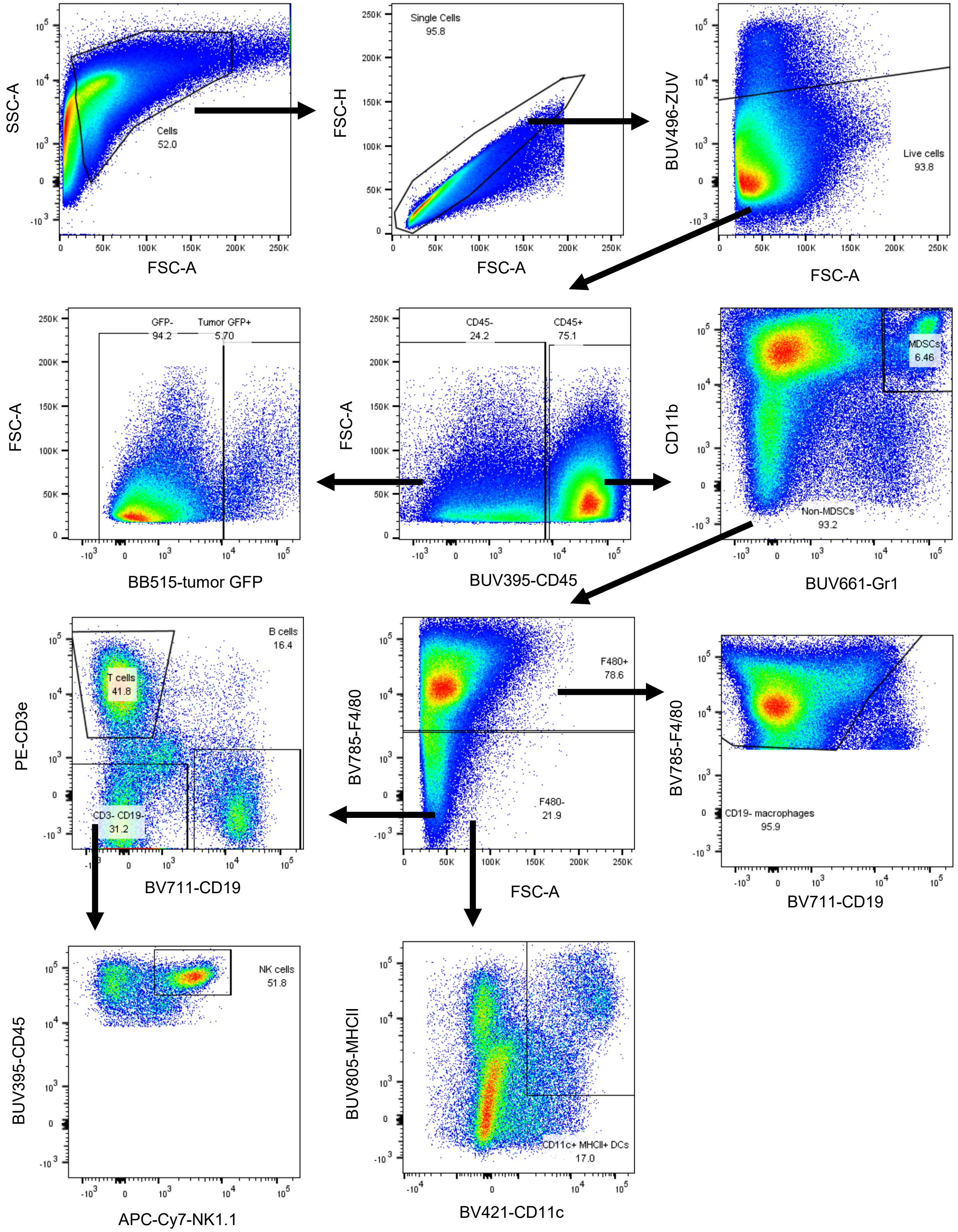


**Fig. S8. Gating strategy for cellular NP distribution in tumor and ascites.** Representative flow plots of cells from dissociated tumors gated for MDSCs (CD11b$^{+}$Gr-1$^{hi}$), T cells (CD3e$^{+}$), B cells (CD19$^{+}$), NK cells (CD3$^{-}$NK1.1$^{+}$), DCs (CD11c$^{+}$MHC-II$^{+}$), and macrophages (F4/80$^{+}$CD19$^{-}$). Gates set based on corresponding FMOs. The same gating strategy was applied to ascites cells.

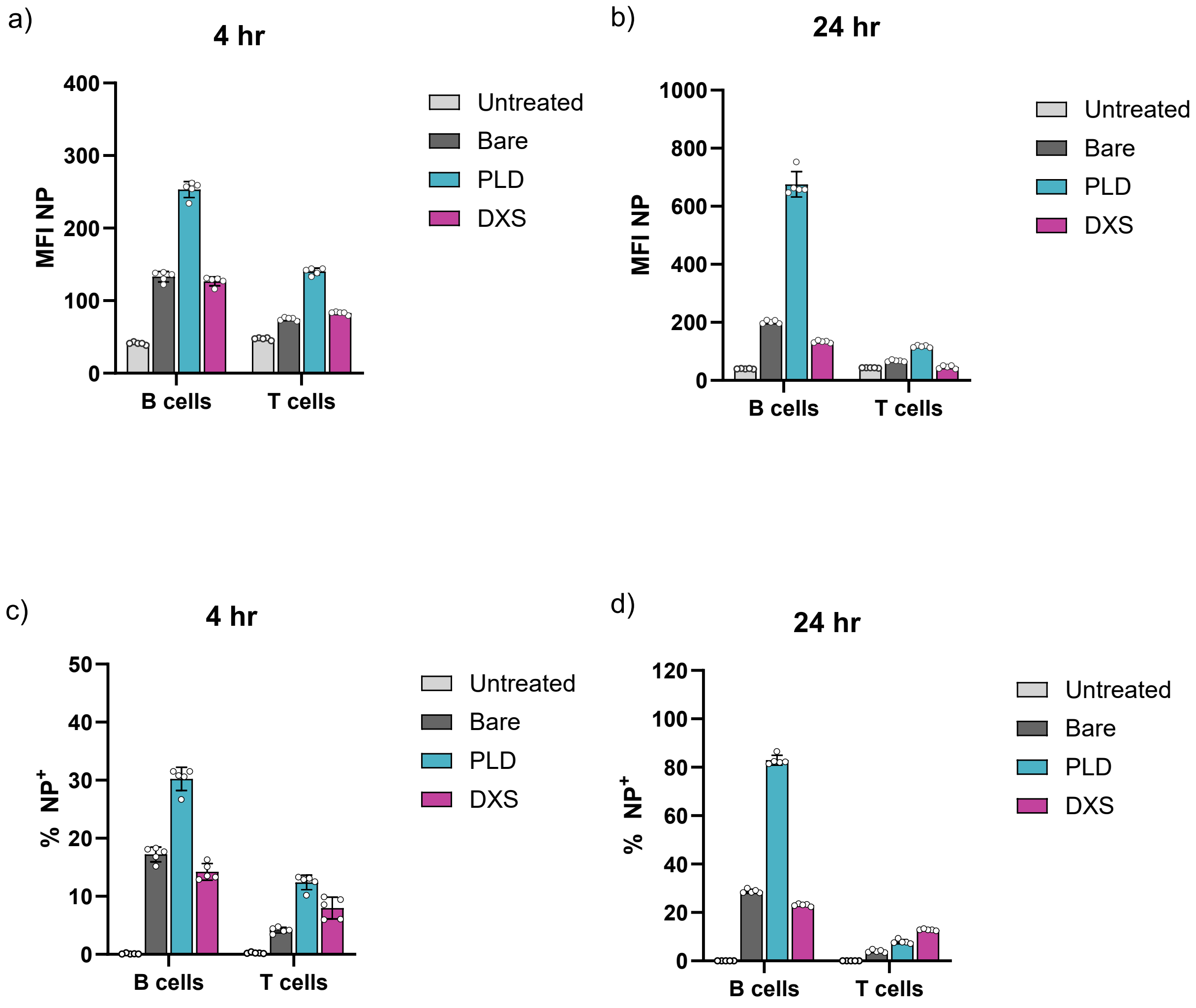


**Fig. S9. Cellular association of NPs with splenocytes assessed using flow cytometry.** Median fluorescence intensity (MFI) of bare and layered BDP-650-labeled NPs in B and T cells after a) 4 hr and b) 24 hr incubation with splenocytes. Percentage of live cells that are positive for NPs after c) 4 hr and d) 24 hr. NPs dosed at 20 μg/mL per well. Data points represent 5 technical replicates. Data represented as mean ± standard deviation.
PLD – poly-L-aspartate; DXS – dextran sulfate.

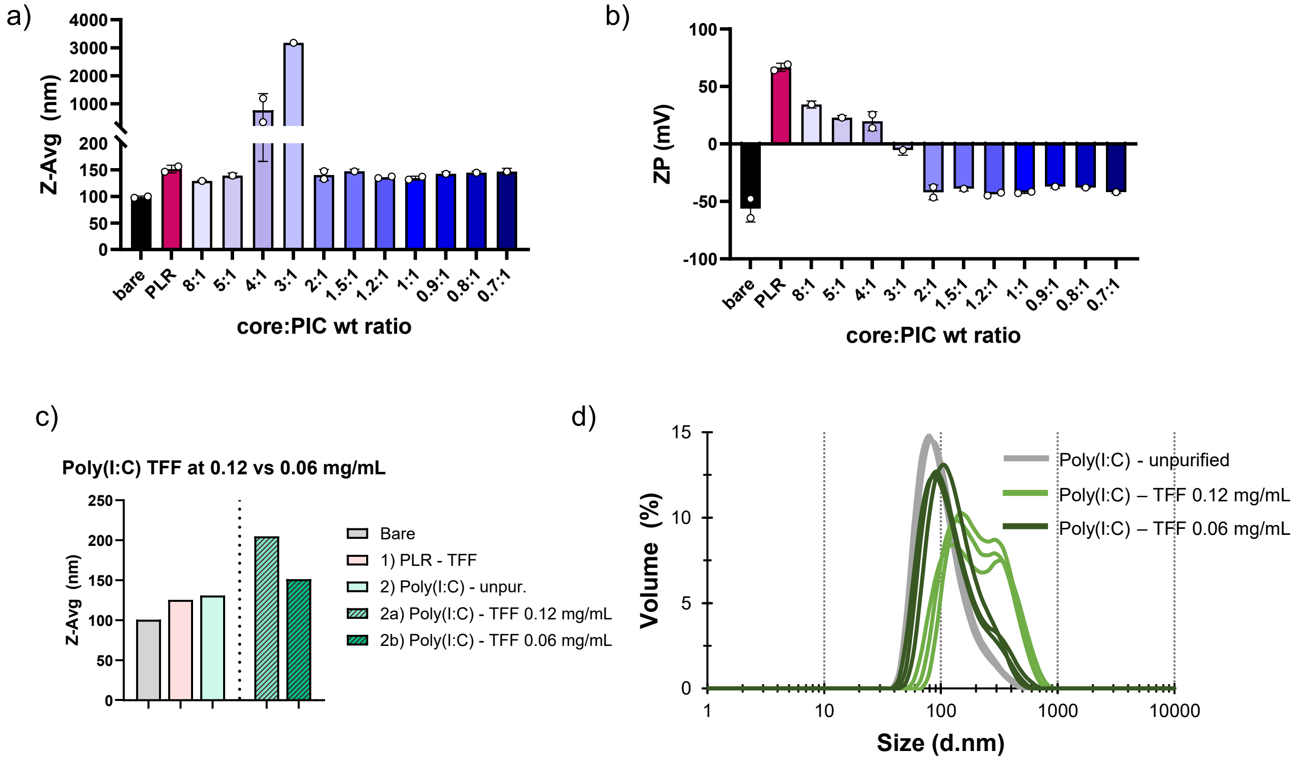


**Fig. S10. Optimization of poly(I:C) layering and purification conditions.** a) Size and b) zeta potential of liposomes layered with poly-L-arginine (PLR)—at weight ratio of 1:0.25 at final lipid concentration of 0.5 mg/mL in 25 mM HEPES, 20 mM NaCl then purified using tangential flow filtration (TFF)—and subsequently layered with various weight ratios of poly(I:C) at final concentration of 0.125-0.25 mg/mL lipid in ~10 mM NaAc, ~5 mM NaCl. N=1-2 independent batches of nanoparticles (mean±SD). c) To optimize purification conditions, a batch of PLR/Poly(I:C) layered NPs (0.8:1 core:poly(I:C) wt ratio) were split and purified using tangential flow filtration at 0.12 mg/mL or 0.06 mg/mL NP (lipid) concentration with a 750kD microKros mPES hollow fiber membrane at 7 mL/min feed flow rate (N=1). Purification at the lower NP concentration preserved the size of the NPs better. d) This is due to lower incidence of bridging between NPs with the lower purification concentration as observed in the volume size distribution of the NPs measured using dynamic light scattering.

**Table. S6. Optimized assembly and purification conditions for LbL Poly(I:C) synthesized using bulk layering and tangential flow filtration (TFF) purification.**

| | Layering conditions | | | Purification conditions | |
|---|---|---|---|---|---|
| Layer | Wt ratio (core:polymer) | Final liposome concentration | Final buffer concentration | Conc. During TFF | Concentration post TFF |
| 1) PLR | 1:0.3 | 0.25-0.5 mg/mL | 25 mM HEPES, 20 mM NaCl | 0.25-0.5 mg/mL | 0.25-0.5 mg/mL |
| 2) Poly(I:C) | 1:1-1.25 | 0.125 mg/mL | 10 mM NaAc, 5 mM NaCl | 0.06 mg/mL | 0.06 mg/mL |
| 3) PLR | 1:1-1.25 | 0.03 mg/mL | 25 mM HEPES, 20 mM NaCl | 0.03 mg/mL | 4X concentration – 0.12 mg/mL |
| 4) DXS | 1:0.4-0.5 | 0.06 mg/mL | 25 mM HEPES, 20 mM NaCl | 0.06 mg/mL | >0.2 mg/mL |

**Table. S7. Assembly conditions for LbL Poly(I:C) synthesized using microfluidics.**

| Layer | Flow rate (channel 1, 2) | Volume ratio (NP:polymer) | Wt ratio (core:polymer) | Final liposome concentration | Final buffer concentration |
|---|---|---|---|---|---|
| 1) PLR | 5 mL/min, 5 mL/min | 1:1 | 1:0.15 | 0.5 mg/mL | 10 mM NaAc |
| 2) Poly(I:C) | 5 mL/min, 5 mL/min | 1:1 | 1:0.6-0.8 | 0.25 mg/mL | 10 mM NaAc, 5 mM NaCl |
| 3) PLR | 5 mL/min, 5 mL/min | 1:1 | 1:0.4-0.54 | 0.12 mg/mL | 10 mM NaAc, 2.5 mM NaCl |
| 4) DXS | 5 mL/min, 5 mL/min | 1:1 | 1:0.3-0.38 | 0.06 mg/mL | 10 mM NaAc, 1.25 mM NaCl |

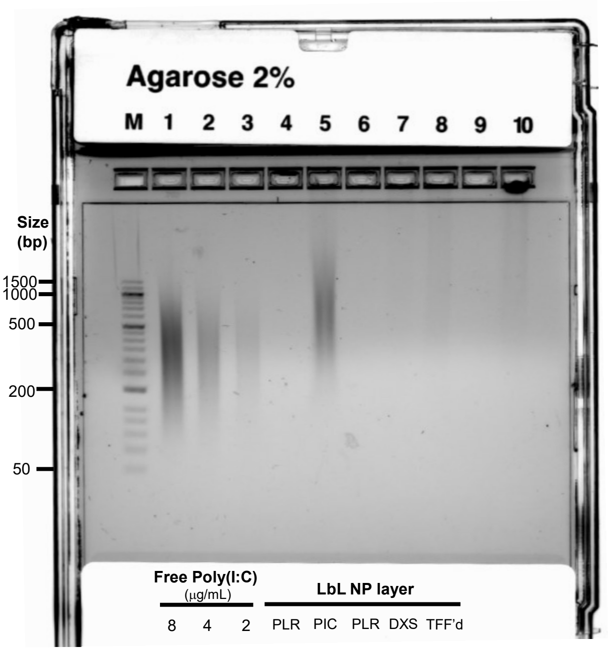


**Fig. S11. Uncropped gel - LbL assembly stably encapsulates poly(I:C) on NP**

Agarose 2% gel of free poly(I:C) at varying concentrations and LbL-Poly(I:C) NP after assembly of each layer. NP samples were concentration matched to 4 μg/mL poly(I:C), or equivalent liposome concentration for the first PLR layer as the NP as this stage doesn't yet contain poly(I:C). E-Gel™ Sizing DNA Ladder (Thermo, #10488100) utilized undiluted at 20 μL per well. E-Gel EX (Thermo, #G401002) loaded with 20 μL sample per well and ran on E-Gel iBase electrophoresis system according to the manufacturer's instructions and imaged on a ChemiDoc imaging system (BioRad) using SYBR gold setting.

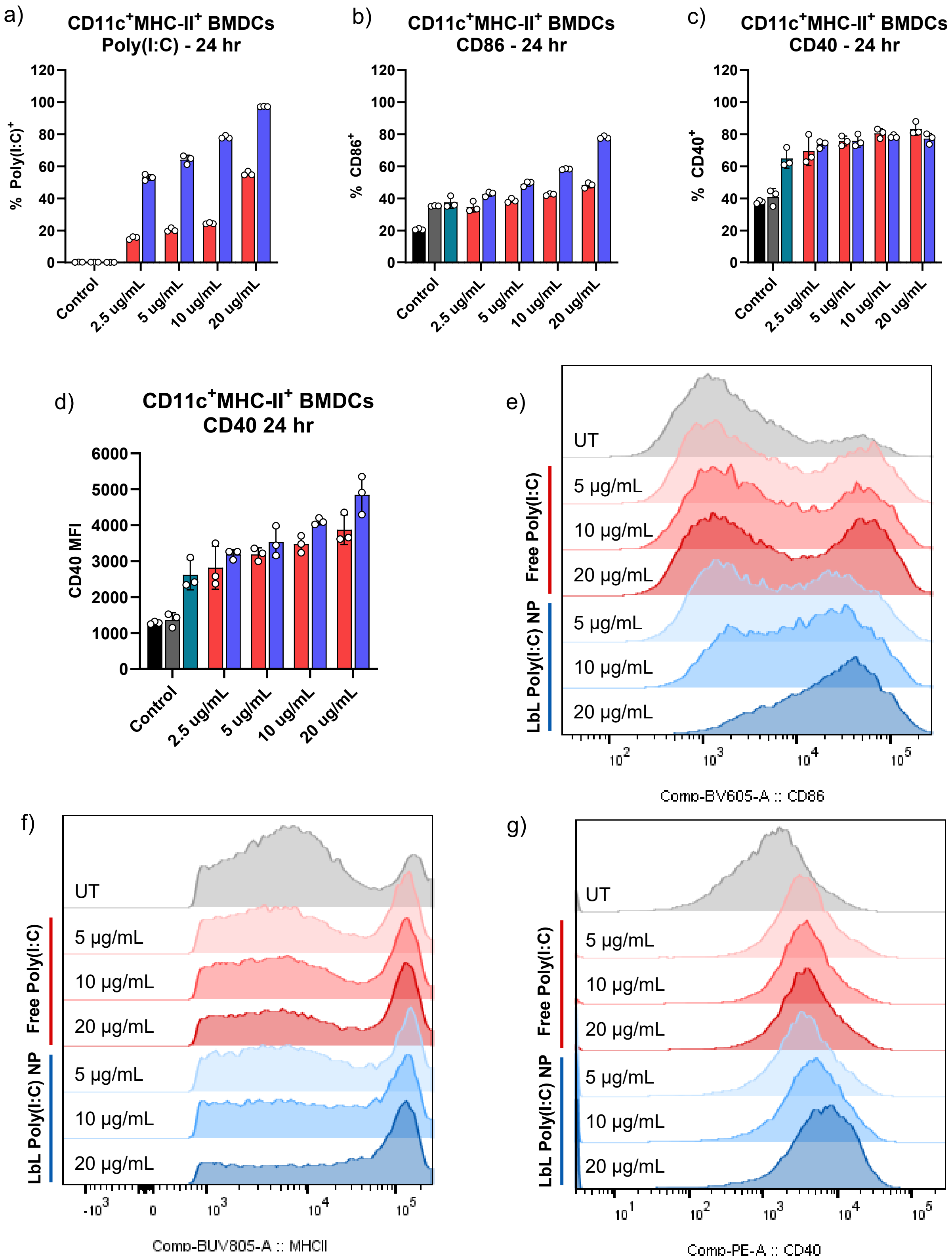


**Fig. S12. BMDC cellular response to LbL Poly(I:C) NPs vs. soluble poly(I:C).** Bone-marrow derived dendritic cells (BMDCs) (same batch as Fig 4a-d) were treated with free poly(I:C), LbL Poly(I:C) NPs (bulk layering + TFF method), or control NPs (bare liposome or PLR/DXS bilayer NPs) for 24hr at varying poly(I:C) final concentrations in the well (N=3 technical replicates). After 24hrs, percentage of cells that are positive for a) poly(I:C)-Cy5 fluorescence, b) CD86, and c) CD40 and d) median fluorescence intensity (MFI) of CD40 expression were measured using flow cytometry. Representative histograms of e) CD86, f) MHC-II, and g) CD40 expression at various poly(I:C) doses. BMDCs were gated as CD11c+/MHCII+ cell population.

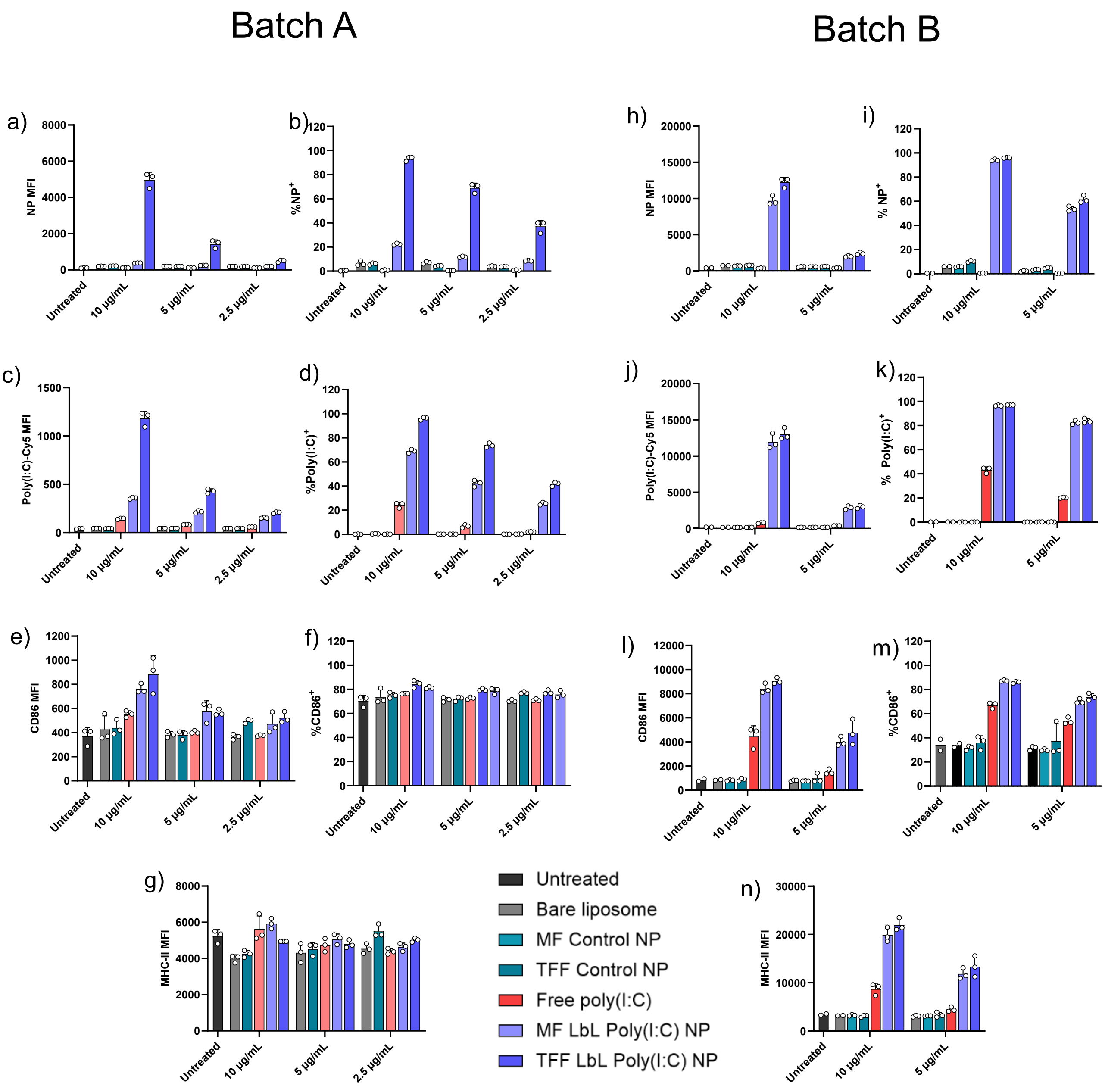


**Fig. S13. BMDC response to LbL Poly(I:C) NPs formulated with bulk layering + TFF vs. microfluidics.** Bone marrow derived dendritic cells were treated with LbL Poly(I:C) NPs at various final poly(I:C) concentrations in the well (N = 3 technical replicates). After 24hrs, NP core (NBD) fluorescence, poly(I:C)-Cy5 fluorescence, CD86 expression, and MHC-II expression were measured using flow cytometry. Data of 2 independent Batches of BMDCs / 2 independent batches of NPs synthesized by different researchers. BMDCs were gated as CD11c$^{+}$/MHC-II$^{+}$ cell population. Data in m) for controls and 10 μg/mL dose reproduced from Fig. 4g to enable comparison across doses.

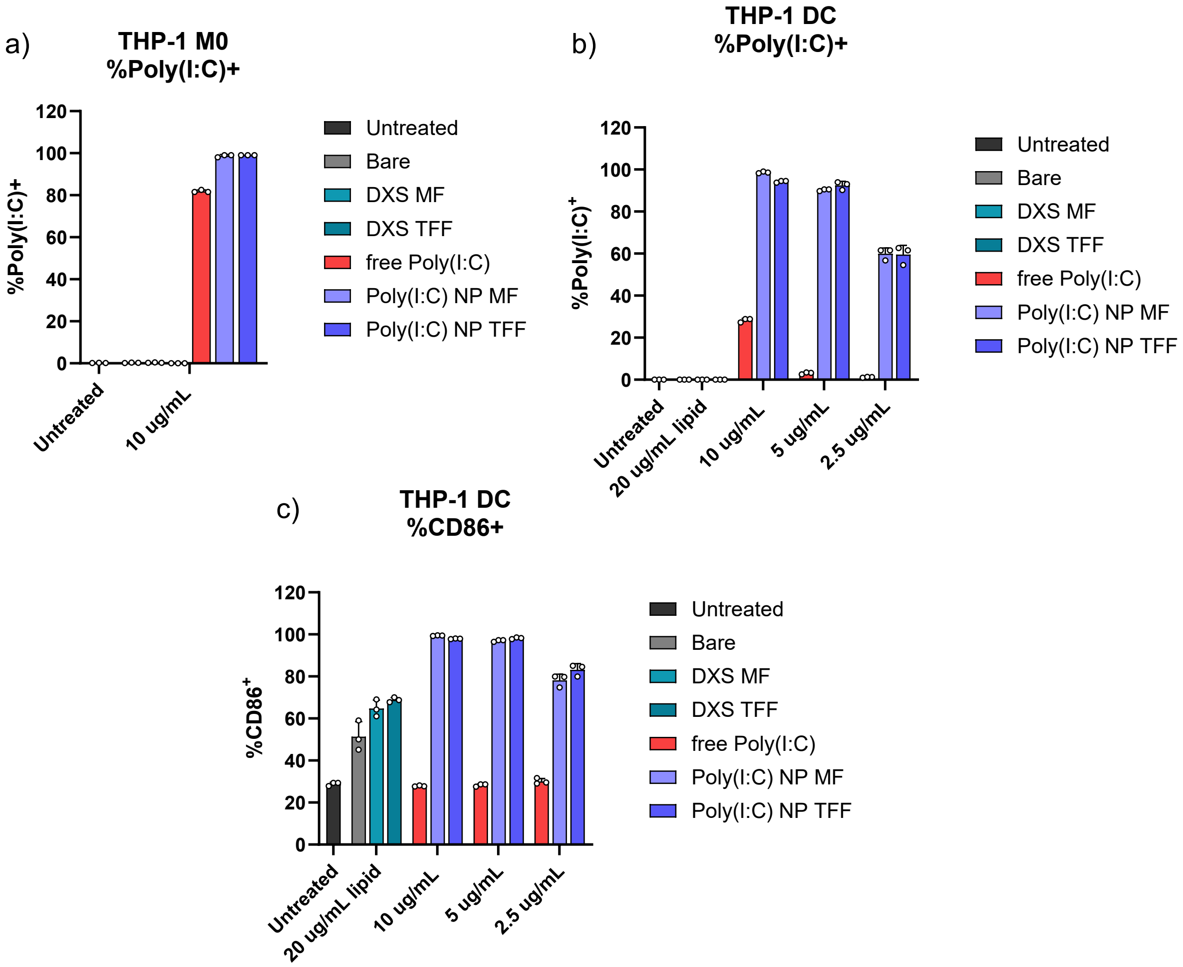


**Fig. S14. THP-1 derived macrophage and dendritic cells cellular response to LbL Poly(I:C) NPs vs. soluble poly(I:C).** THP-1 human monocytes were polarized to M0 macrophages or dendritic cells and then treated with free poly(I:C), LbL Poly(I:C) NPs (bulk layering + TFF method), or control NPs (bare liposome or PLR/DXS bilayer NPs) for 24hr at varying poly(I:C) final concentrations in the well (N = 3 technical replicates). After 24hrs, percentage of cells that are positive for a-b) poly(I:C)-Cy5 fluorescence and c) CD86 were measured using flow cytometry. Data at 10 µg/mL doses and controls reproduced from Fig. 5h-I to enable comparisons at various doses.

**THP-1 M0 generation:** THP-1 cells were centrifuged at 130xg for 10 min to pellet cells. Media was aspirated, cells resuspend in fresh THP-1 media and counted. Cells were diluted to $5x10^5$ cell/mL and final concentration of 50 ng/mL PMA, diluted from 1000X PMA stock (in DMSO). To a flat-bottom, TC-treated plate 150 µL cell suspension was added to each well (75k cells / well). After 2 days, the stimulation media was replaced: the media was removed, cells gently washed with 200 µL PBS, and 90 µL THP-1 standard media added to each well to allow the cells to rest from stimulation for 1 day before using in the assay.

**THP-1 DC generation:** THP-1 cells were centrifuged at 130xg for 10 min to pellet cells. Media was aspirated, cells resuspend in fresh THP-1 media and counted. Cells were diluted to $2x10^5$ cell/mL and final concentration of 55 µM beta-mercaptoethanol, 100 ng/mL IL-4, and 100 ng/mL GM-CSF and cultured in a T75 flask. After 72 hr, the cells were centrifuged, media removed, and cells resuspended in fresh, supplemented media and returned to a T75 flask for culture. After 48 hr, cells were centrifuged, media removed, cells counted and plated for the assay at 50k/well in 96 well, TC-treated flat bottom plate and used in the assay.

a)

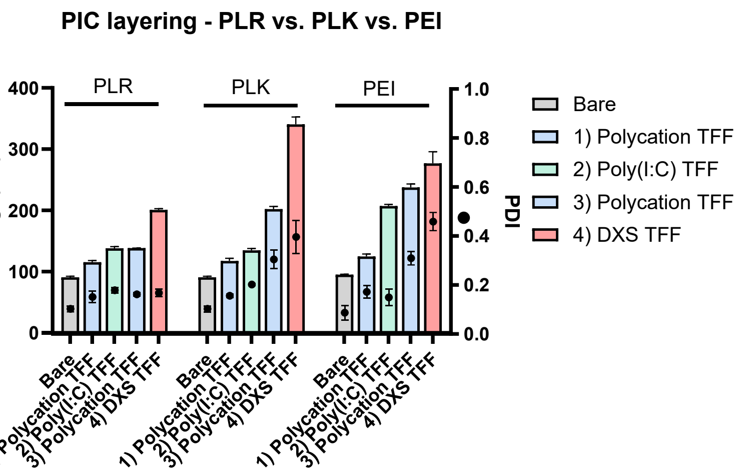


b)

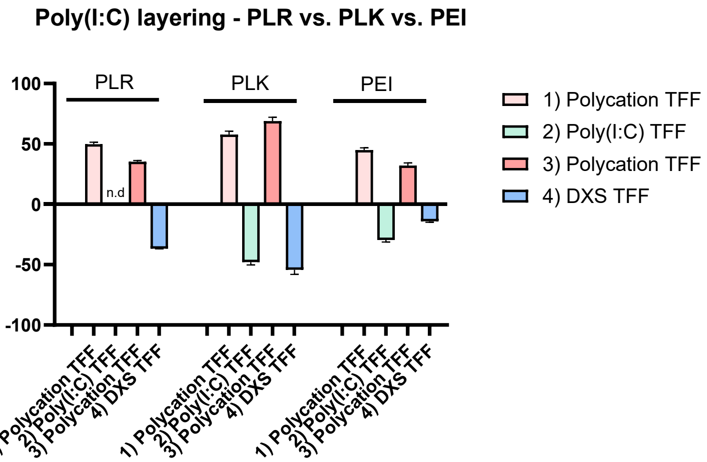


**Fig. S15. Characterization of LbL Poly(I:C) NPs with various complexing polycations.** LbL tetralayer Poly(I:C) NPs were generated using different polymers as the complexing polycation: poly-L-arginine (PLR, MW 38,500kD, 200 repeat units), poly-L-lysine (PLK, MW 41,000, 250 repeat units, and linear polyethyleneimine (PEI, 25kD), all with dextran sulfate (DXS) serving as the terminal polyanion layer. PLR and PLK were layered in 25 mM HEPES, 20 mM NaCl and PEI layered in 10 mM NaAc, 5 mM NaCl. All formulations were layered at the same core: PIC ratio (0.8:1). Each formulation was layered at optimized ratios for each polycation and purified using similar TFF conditions. a) Size, polydispersity index (PDI) and b) zeta potential (ZP) of LbL Poly(I:C) NPs. Data are shown as mean±standard deviation of each batch measured in triplicate.

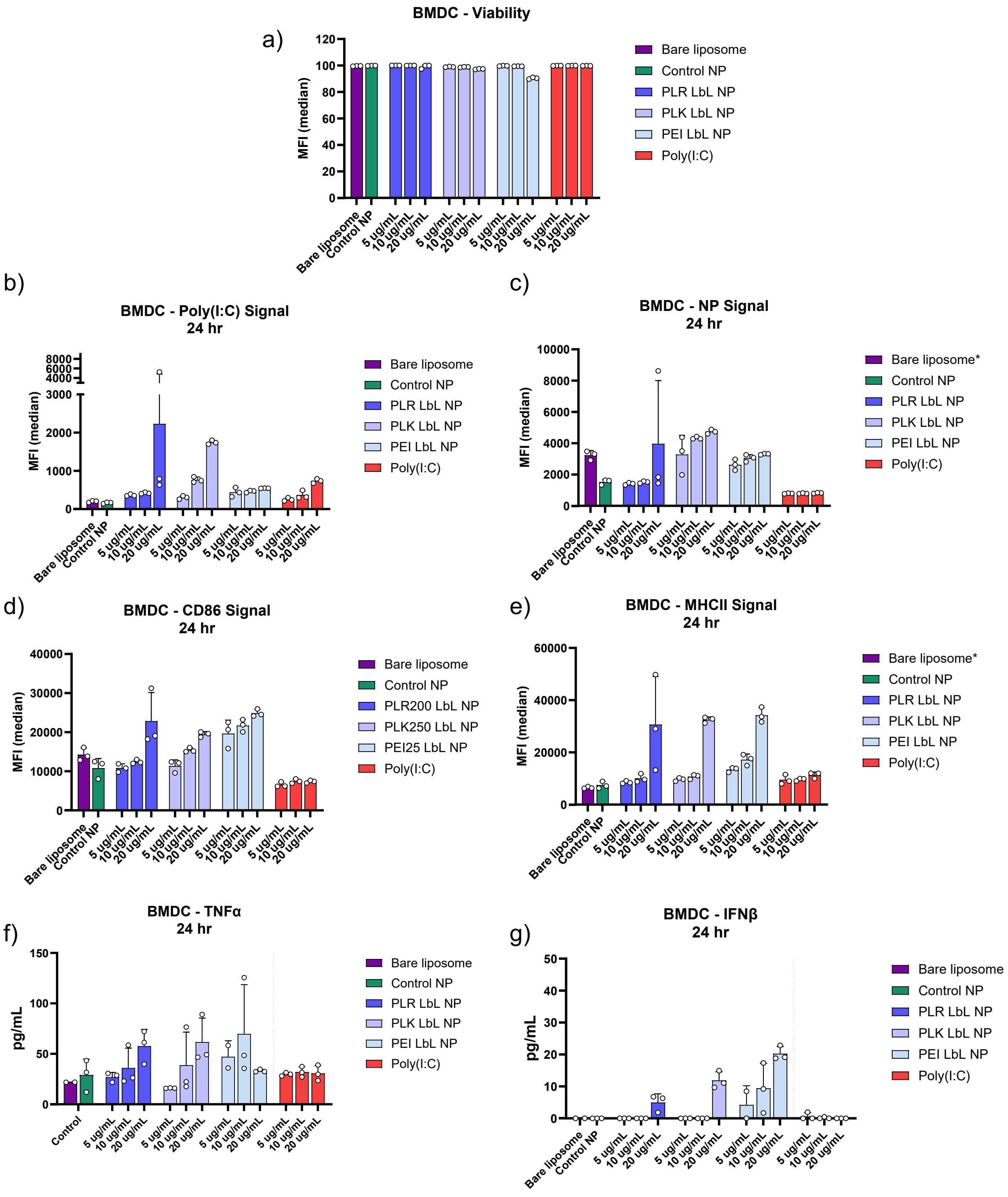


**Fig. S16. BMDC cellular response to LbL Poly(I:C) NPs formulated with various complexing polycations.** Bone-marrow derived dendritic cells (BMDCs) were treated with LbL Poly(I:C) NPs or control NPs (bare liposome or PLR/DXS bilayer NPs) for 24hr at 20, 10, 5 μg/mL poly(I:C) final concentration in the well (N = 3 technical replicates). After 24hrs, a) cell viability, b) poly(I:C)-Cy5 fluorescence, c) NP core (NBD) fluorescence, d) CD86 expression, and e) MHC-II expression were measured using flow cytometry. BMDCs were gated as CD11c⁺/MHC-II⁺ cell population. f) TNFα and g) IFNβ levels in the cell supernatant were measured using ELISAs. For TNFα ELISA, data for PLR LbL NP and free poly(I:C) reproduced from Fig 4. Values below limit of detection are plotted as 0.

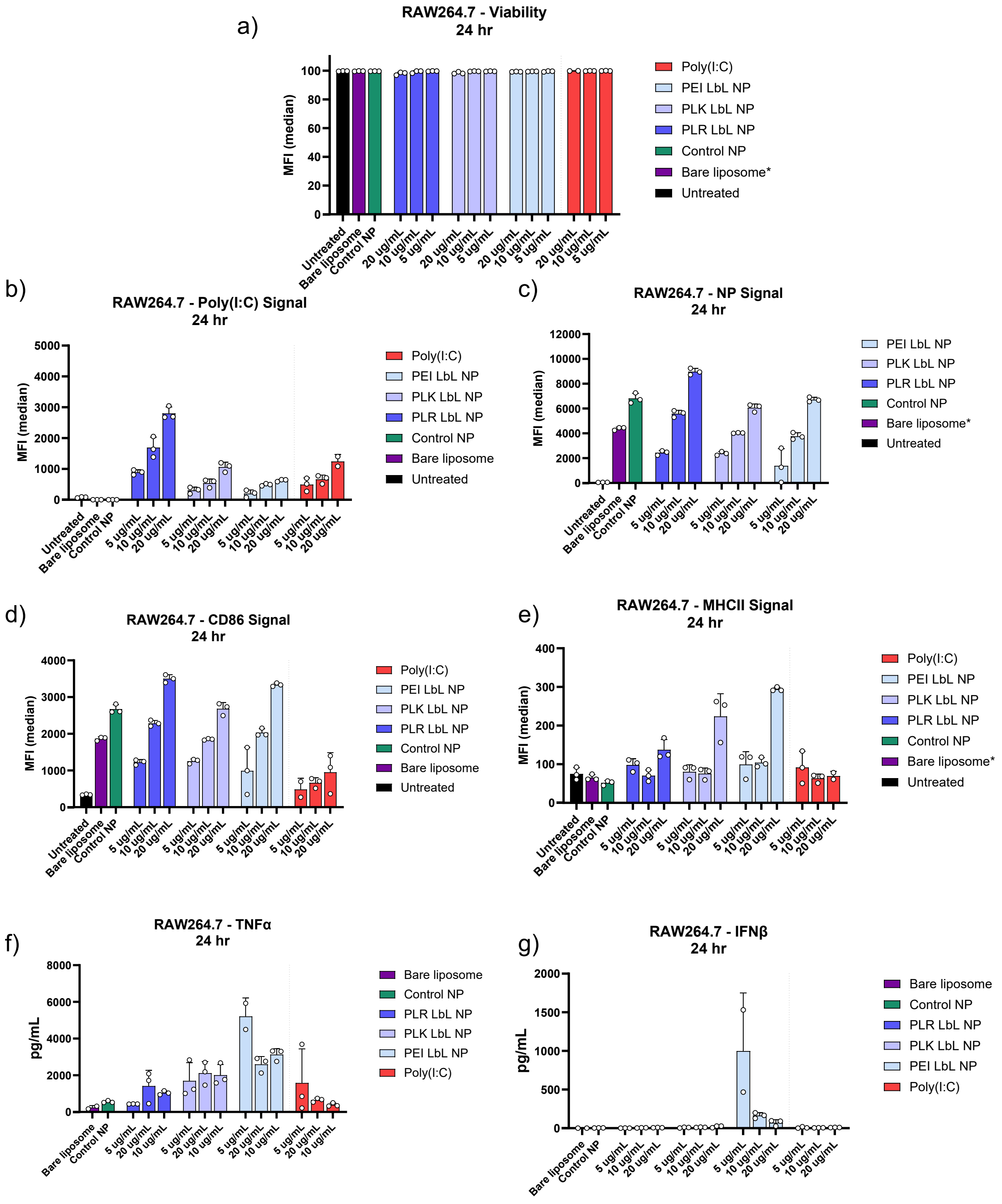


**Fig. S17. RAW264.7 cellular response to LbL Poly(I:C) NPs formulated with various complexing polycations.** RAW264.7 macrophages were treated with LbL Poly(I:C) NPs or control NPs (bare liposome or PLR/DXS bilayer NPs) for 24hr at 20, 10, 5 µg/mL poly(I:C) final concentration in the well (N = 3 technical replicates). After 24hrs, a) cell viability, b) poly(I:C)-Cy5 fluorescence, c) NP core (NBD) fluorescence, d) CD86 expression, and e) MHC-II expression were measured using flow cytometry. Data for PLR LbL NP, free poly(I:C) and controls reproduced from Fig. 4. f) TNFα and g) IFNβ levels in the cell supernatant were measured using ELISAs. Values below limit of detection are plotted as 0.

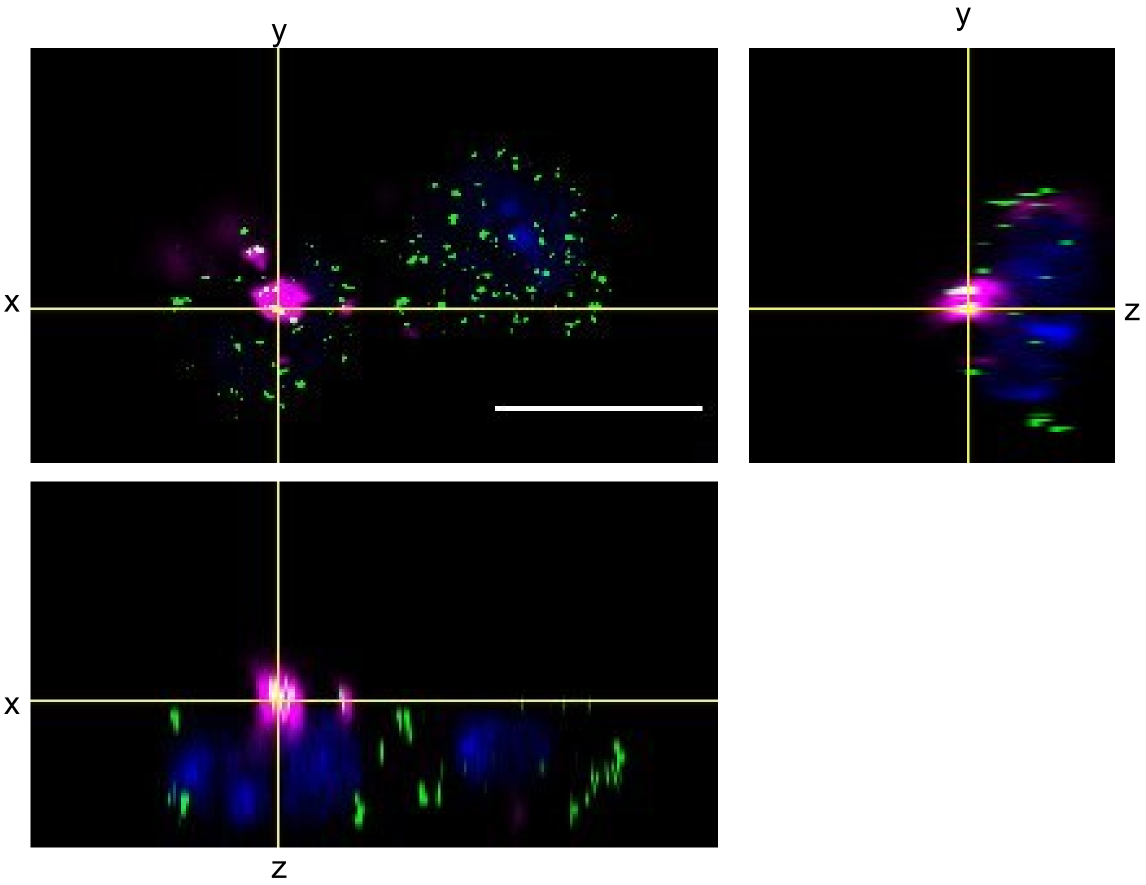


**Fig. S18. LbL poly(I:C) NP co-localizes with TLR3 on macrophage cell surface after 1 hr.** Orthogonal views of confocal microscopy of RAW264.7 cells treated with LbL Poly(I:C) NPs at 10 µg/mL for 1 hr. Cells were stained with Hoechst to visualize nuclei (blue) and anti-TLR3 antibody+fluorescent secondary antibody (green). 5% Cy5-poly(I:C) was included to enable its visualization (magenta). Images were acquired on an Evident FV4000 with 100X objective. Scale bar = 10 µm.

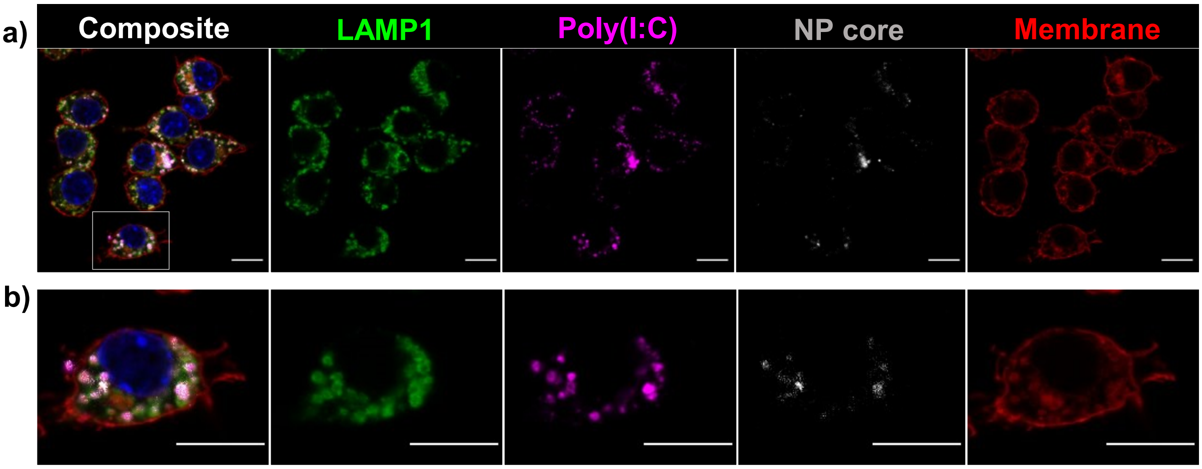


**Fig. S19. LbL poly(I:C) NP co-localizes with lysosomes after 4 hr.** Confocal microscopy of RAW264.7 cells treated with LbL Poly(I:C) NPs at 10 µg/mL for 4 hr. Cells were stained with Hoechst to visualize nuclei (blue) and 5% Cy5-poly(I:C) included to enable its visualization (magenta). Poly(I:C) is internalized and co-localizes with NP core (gray) and vesicles positive for LAMP1 (green), a marker for lysosomes. b) Zoom in of cell outlined by gray box in a. Images acquired on an Evident FV4000, 100X objective. Scale bar = 10 µm.

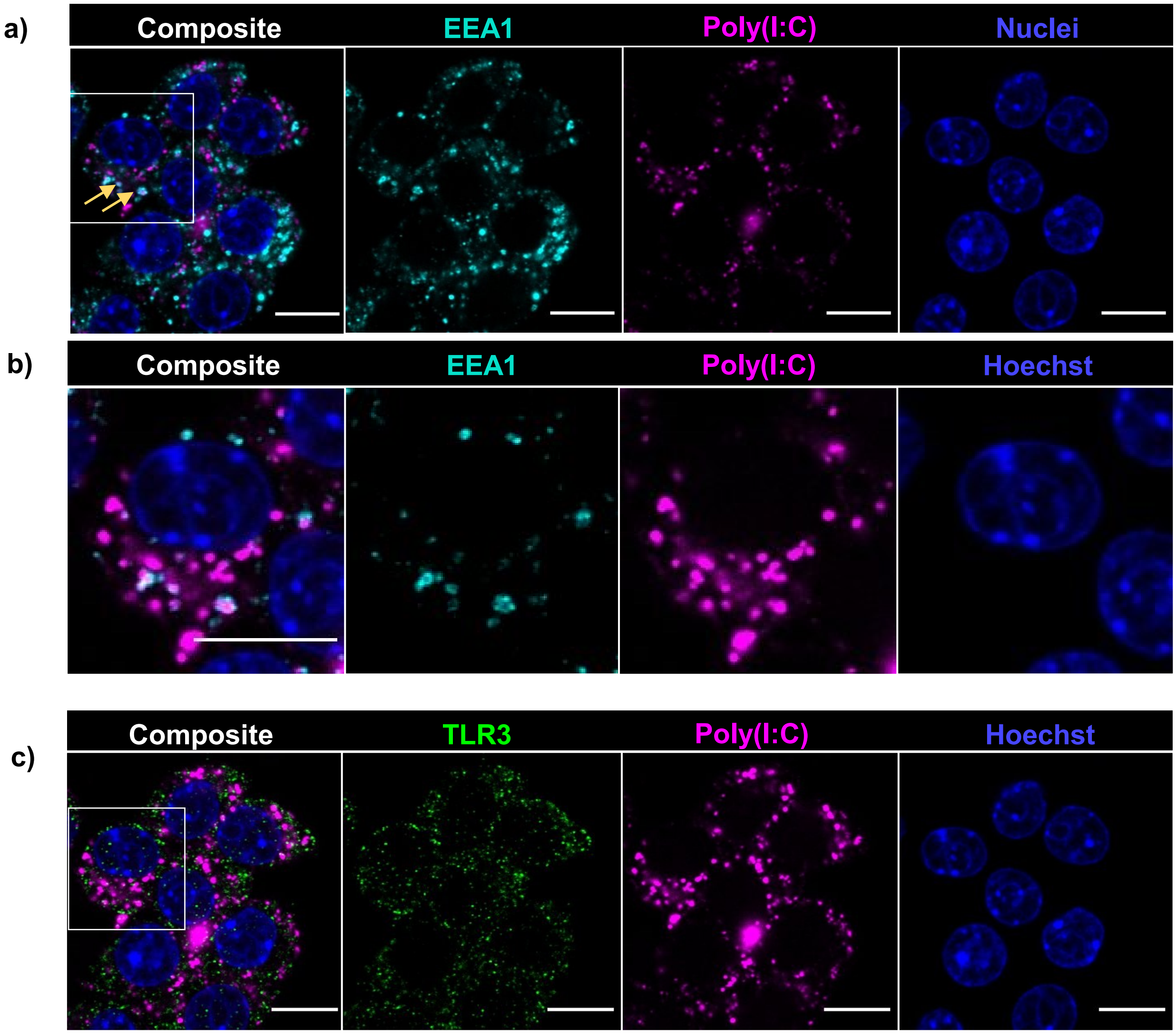


**Fig. S20. LbL poly(I:C) NP intracellular localization and interactions with dsRNA sensors.** Confocal microscopy of RAW264.7 cells treated with LbL Poly(I:C) NPs at 10 µg/mL for 4 hr. Cells were stained with Hoechst to visualize nuclei (blue) and 5% Cy5-poly(I:C) was included to enable its visualization (magenta). a) Poly(I:C) is found in EEA1-negative vesicles as well as EEA1-positive (cyan) endosomes (yellow arrows). b) Zoom in of cell outlined by gray box in a).

c) The same cells were also stained for TLR3 (green) – Some instances of poly(I:C) co-localization with TLR3 is observed. The zoomed in view of the cell outlined by gray box is found in Fig. 5b. Images were acquired on an Evident FV4000 with 100X objective. Scale bar = 10 µm.

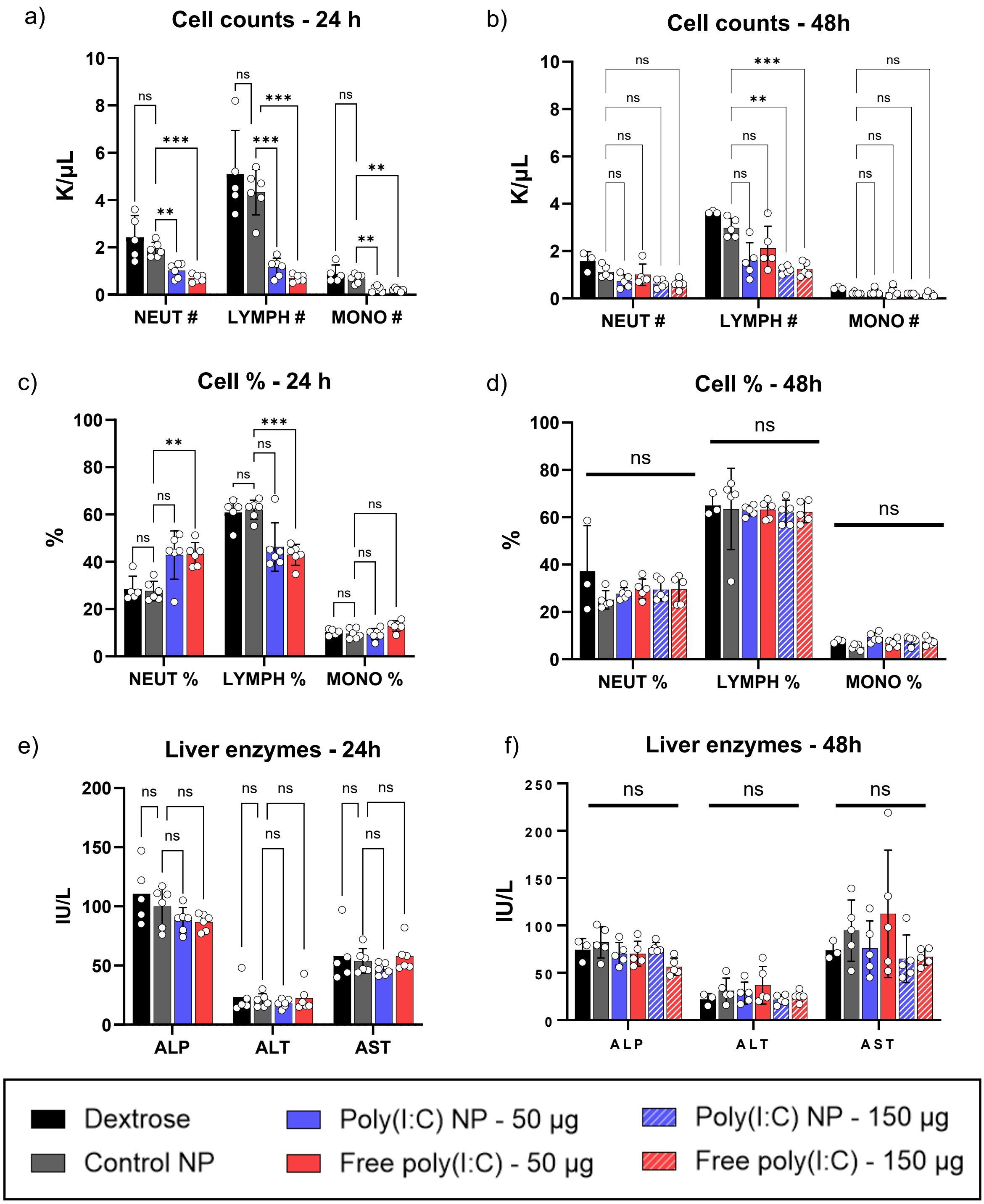


**Fig. S21. Toxicity of free vs. NP poly(I:C) 24hr and 48hr post single dose.** Mice with BPPNM-luc tumors were treated with 50 µg or 150 µg free poly(I:C) or LbL poly(I:C) NP, or with 5% dextrose (vehicle control) or control NP (PLR/DXS layered) at equivalent lipid dose intraperitoneally. After 24 and 48 hrs, blood was sampled via submandibular bleed to evaluate cell levels and serum chemistry for toxicity. a-b) concentration of cell populations (neutrophil, NEUT; lymphocytes; LYMPH; monocytes, MONO), c-d) percentage of cell populations, and e-f) serum level of the liver enzymes alkaline phosphatase (ALP), alanine transaminase (ALT), aspartate aminotransferase (AST). For 24hr, N = 6 for Control NP, Poly(I:C) NP, and free poly(I:C); and N = 5 for Dextrose control. For 48hr, N = 5 for Control NP, Poly(I:C) NP (50 & 150 µg), and free poly(I:C) (50 & 150 µg); and N = 3 for Dextrose control. Statistical significance for cell counts, cell%, and liver enzymes determined used two-way ANOVA with Tukey's multiple comparisons test.

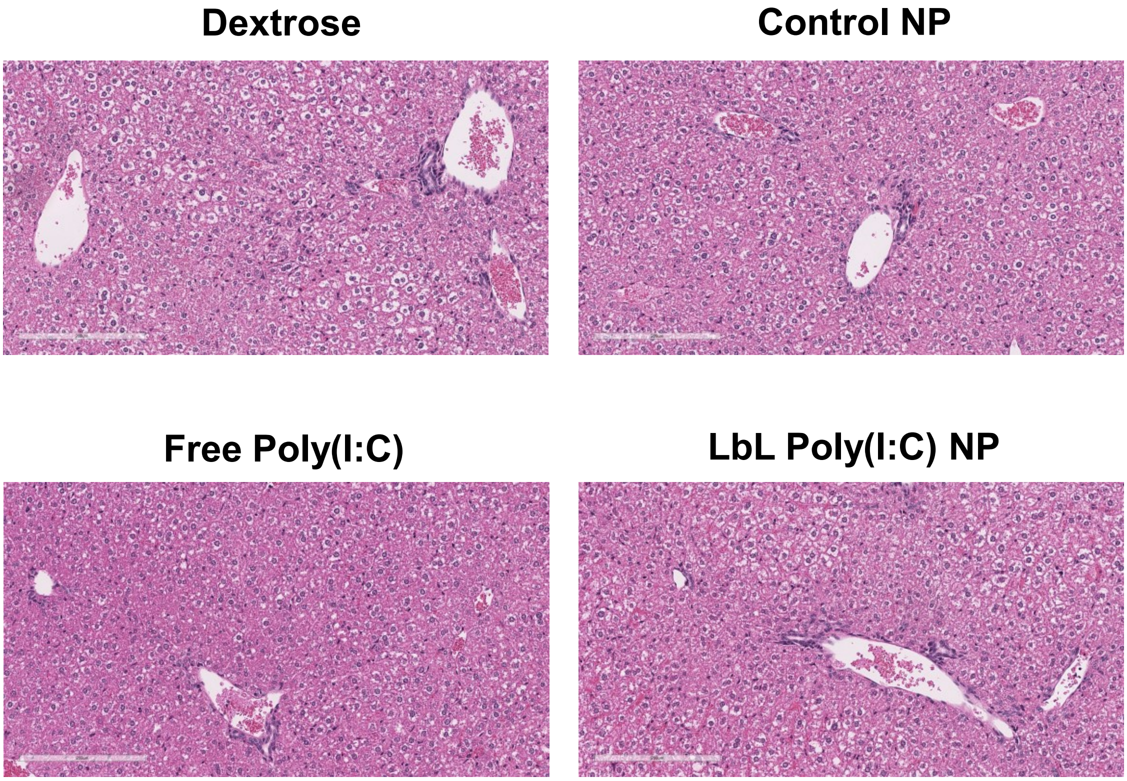


**Fig. S22. Liver histology 24hr after single dose of free vs. NP poly(I:C).** C57BL/6 mice with BPPNM-luc tumors were treated with 50 μg free poly(I:C) or LbL poly(I:C) NP or 5% dextrose (vehicle control) or control NP (PLR/DXS layered) at equivalent lipid dose intraperitoneally. After 24 hrs, livers were removed and processed for H&E staining and histopathological analysis. Scale bars = 200 μm. Shown are representative images from 5-6 biologically independent replicates.

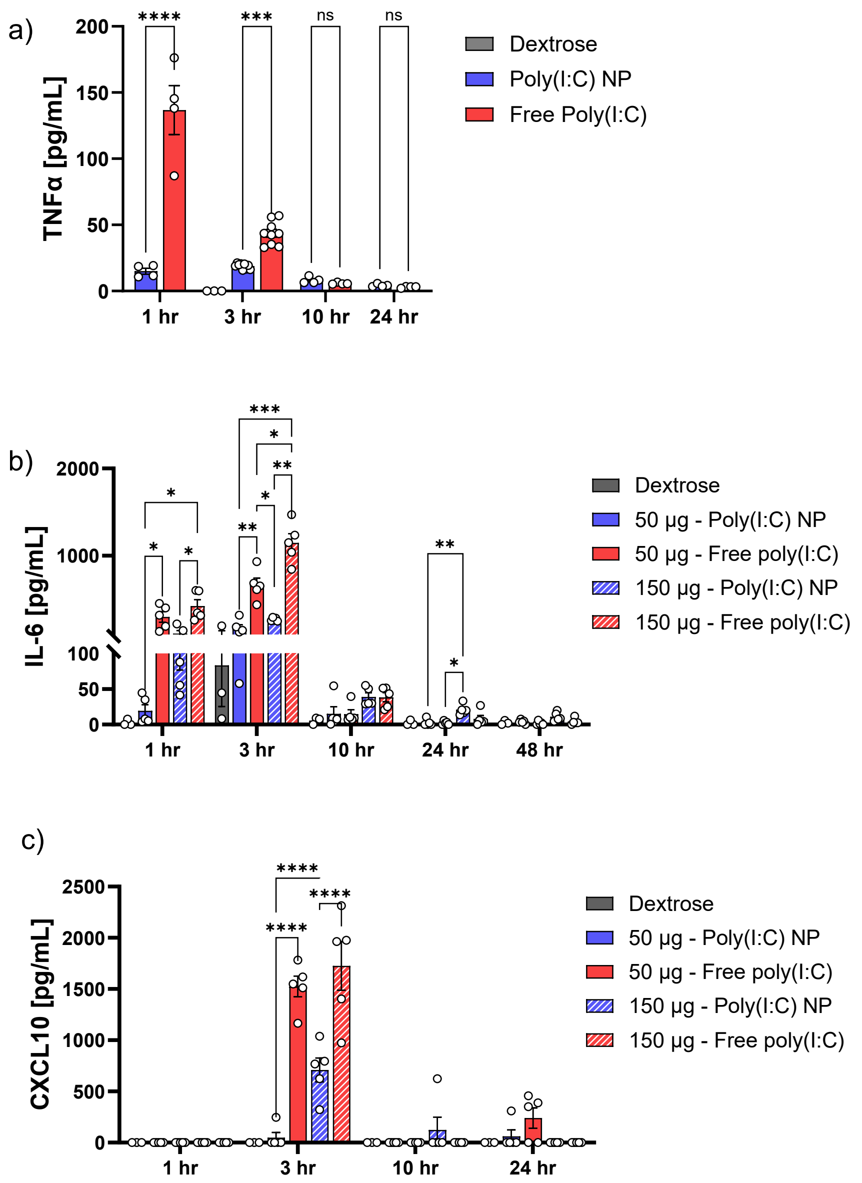


**Fig. S23. Serum cytokine levels post single dose of free or NP encapsulated poly(I:C).** Mice with BPPNM-luc tumors were treated with 50 μg or 150 μg free poly(I:C) or LbL poly(I:C) NP or 5% dextrose (vehicle control) intraperitoneally. Blood was sampled via submandibular bleed to evaluate serum levels of a) TNF-α, b) IL-6, and C) CXCL10, N = 3 for dextrose and N = 5 for Poly(I:C) NP (50 & 150 μg) and free poly(I:C) (50 & 150 μg); Statistical significance determined using Mixed effects analysis with Bonferroni's multiple comparisons test for a) and two-way ANOVA with Tukey multiple comparisons test for b-c.

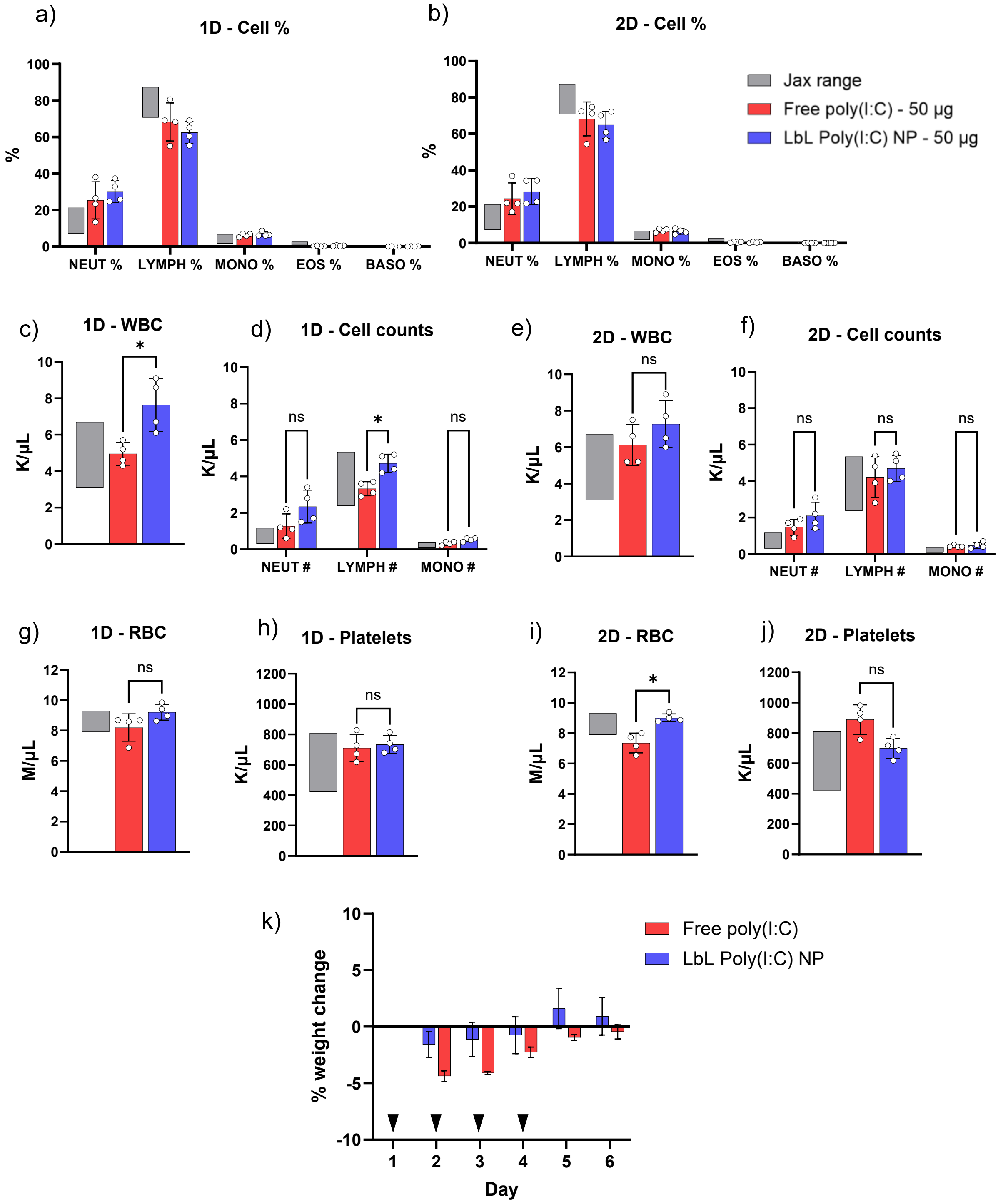


**Fig. S24. Toxicity of free vs. NP poly(I:C) 1D and 2D after four daily doses.** Mice with BPPNM-luc tumors were dosed intraperitoneally with 50 µg free poly(I:C) or LbL poly(I:C) NP in 5% dextrose daily for 4 days (Day 1-4). 24 and 48 hrs after the last dose, blood was sampled via submandibular bleed to evaluate cell levels for toxicity, N=4 per group. a-b) Percentage of cell populations (neutrophil, NEUT; lymphocytes; LYMPH; monocytes, MONO). c, e) Overall white blood cells (WBC) counts and d, f) counts of cell populations. g, i) Red blood cell (RBC) and h, j) platelet levels. k) Weight changes from baseline (Day 1, first dose) were monitored (mean±SEM). Black arrows denote dosing days. Statistical significance for cell percentages and cell counts determined using two-way ANOVA with Bonferroni multiple comparisons test. Statistical significance for WBC, RBC, and platelets determined using Mann-Whitney non-parametric U test.

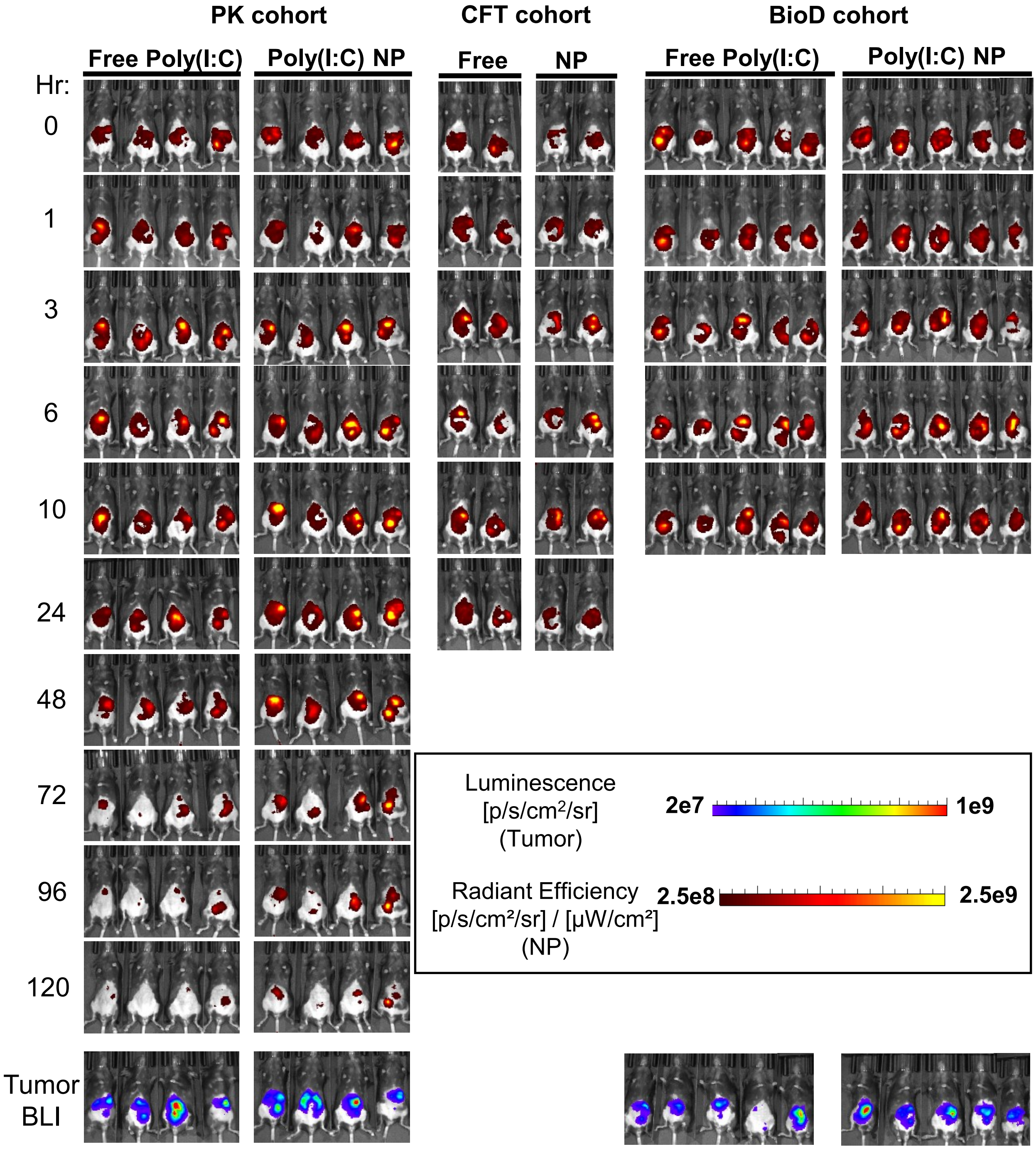


**Fig. S25. In vivo pharmacokinetics of free vs. LbL NP poly(I:C) upon intraperitoneal administration.** Female C57BL/6 mice with BPPNM-luc tumors (3 million, IP) were injected with 50 μg of free or LbL NP encapsulated poly(I:C), labeled with Cy5, in 5% dextrose 10 days post tumor inoculation. Mice were maintained on alfalfa-free diet to minimize fluorescence signal from the chow. Mice were imaged using an in vivo imaging system (IVIS) for Cy5-poly(I:C) fluorescence at various timepoints.

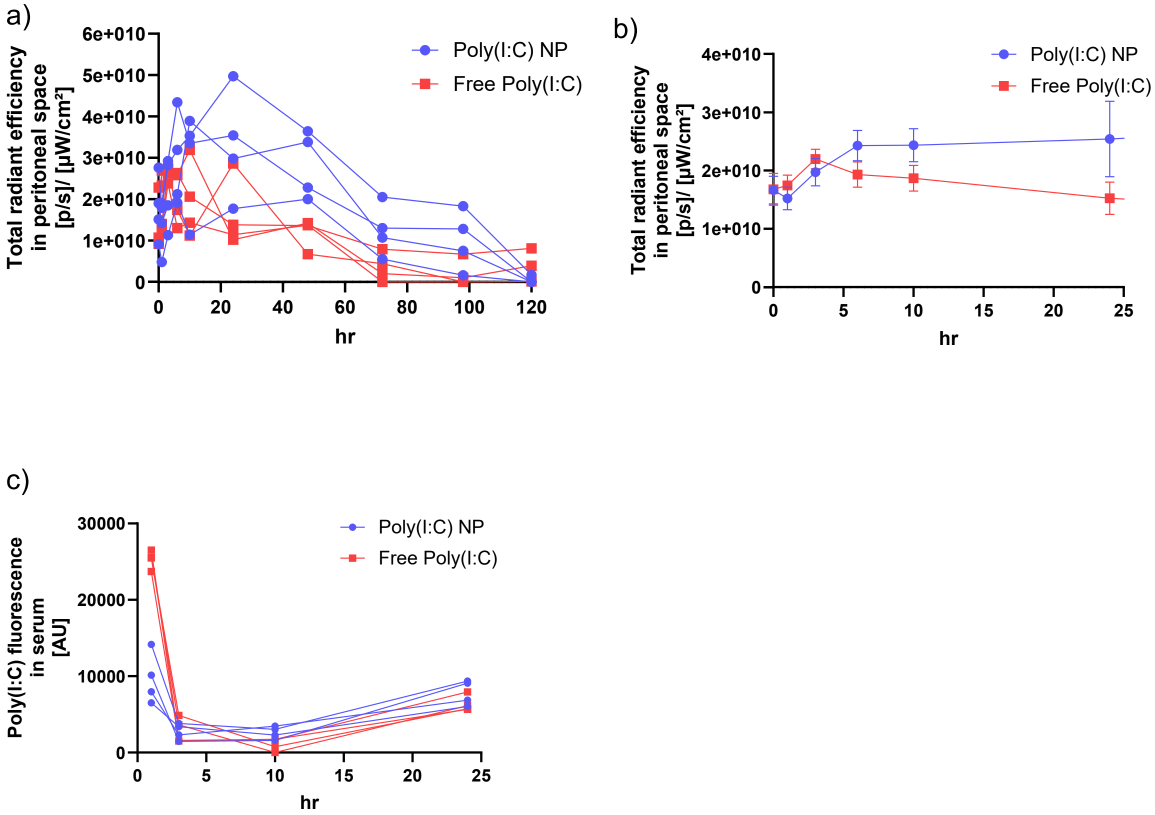


**Fig. S26. In vivo pharmacokinetics of free vs. LbL NP poly(I:C) upon intraperitoneal administration.** a) Female C57BL/6 mice with BPPNM-luc tumors (3 million, IP) were injected with 50 μg of free or LbL NP encapsulated poly(I:C), labeled with Cy5, in 5% dextrose 10 days post tumor inoculation. Mice were maintained on alfalfa-free diet to minimize fluorescence signal from the chow. Mice were imaged using an in vivo imaging system (IVIS) for Cy5-poly(I:C) fluorescence at various timepoints. a) Raw values of poly(I:C) fluorescence (total radiant efficiency) in peritoneal space over 120 hr post-dosing (N=4 , PK cohort in Fig. S25) b) Poly(I:C) fluorescence in peritoneal space averaged across all mice. N = 11 (0-10hr – PK, BioD, and CFT cohorts in Fig. S25) & N=6 (24hr – PK and CFT cohorts in Fig. S25) (mean±SEM) c) Cy5-poly(I:C) fluorescence in the serum for each individual mouse (N=4).

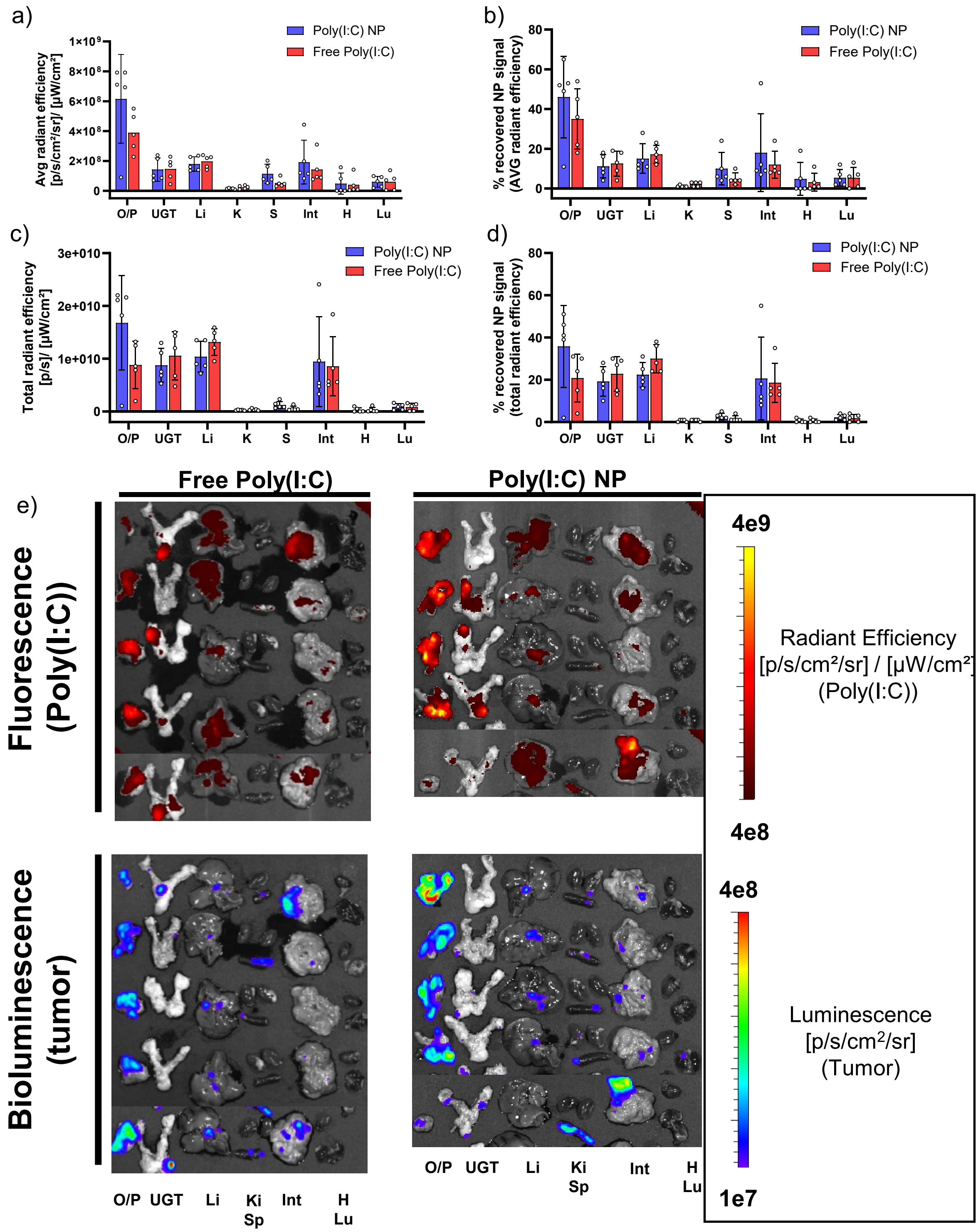


**Fig. S27. Biodistribution of free vs. LbL NP poly(I:C) 24hr after intraperitoneal administration.** Female C57BL/6 mice with BPPNM-luc tumors (3 million, IP) were injected with 50 µg of free or LbL NP encapsulated poly(I:C), labeled with Cy5, in 5% dextrose. Mice were maintained on alfalfa-free diet to minimize fluorescence signal from the chow. 24 hr after treatment, organs were collected and imaged using an in vivo imaging system (IVIS) for Cy5-poly(I:C) fluorescence and tumor bioluminescence. Poly(I:C)-Cy5 fluorescence—a) average radiant efficiency and c) total radiant efficiency normalized by organ weight—in each organ and b,d) corresponding percent recovered fluorescence across the organs. Fluorescence from dextrose-treated mice was used to subtract background. N=5 per group (mean±SD). e) Images of organs for poly(I:C) fluorescence and tumor bioluminescence. O/P – omentum/pancreas; UGT – upper genital tract; Li – liver; Ki – kidney; Sp – spleen; Int – intestines; H – heart; Lu – lungs.

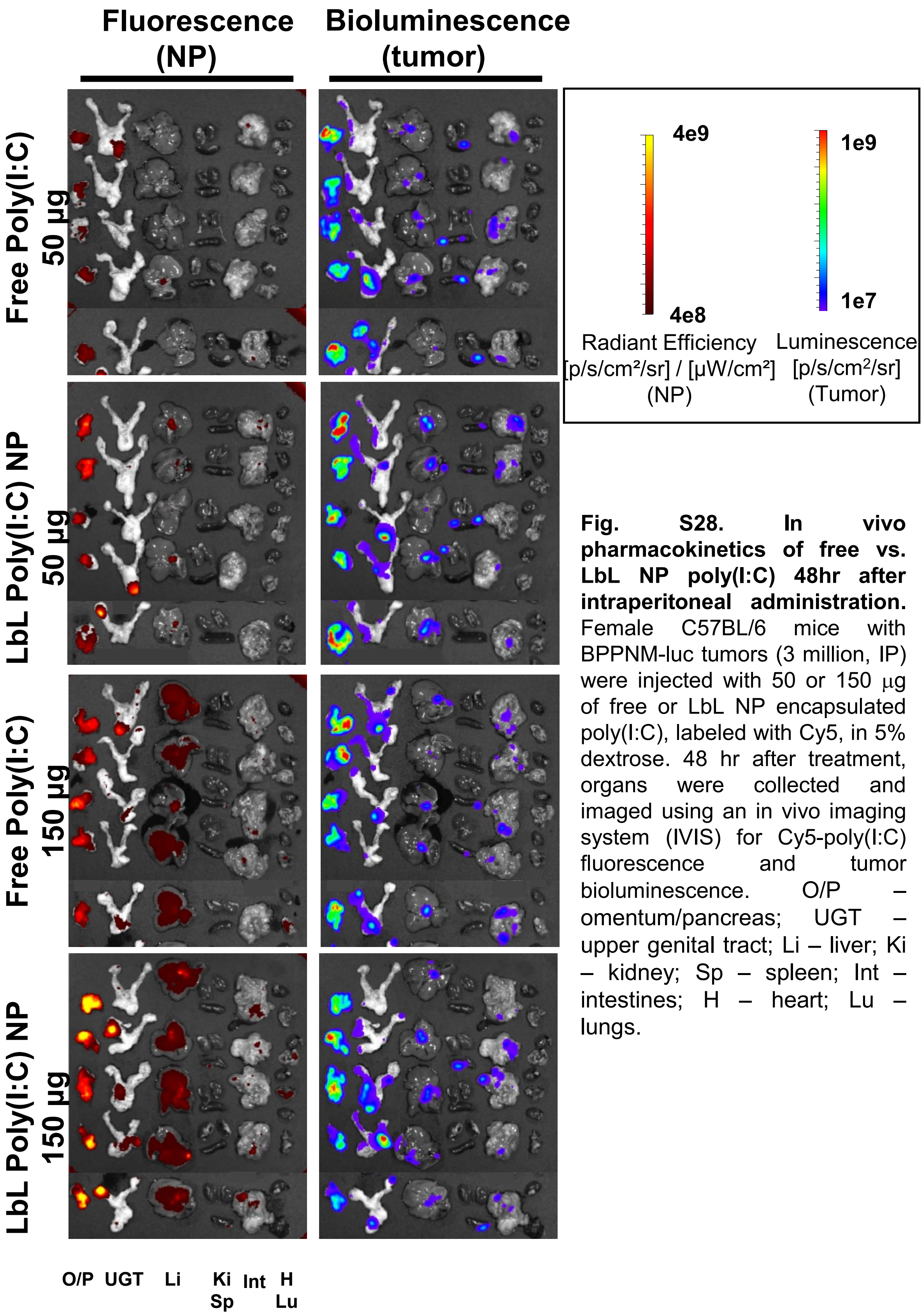


**Fig. S28. In vivo pharmacokinetics of free vs. LbL NP poly(I:C) 48hr after intraperitoneal administration.** Female C57BL/6 mice with BPPNM-luc tumors (3 million, IP) were injected with 50 or 150 µg of free or LbL NP encapsulated poly(I:C), labeled with Cy5, in 5% dextrose. 48 hr after treatment, organs were collected and imaged using an in vivo imaging system (IVIS) for Cy5-poly(I:C) fluorescence and tumor bioluminescence. O/P – omentum/pancreas; UGT – upper genital tract; Li – liver; Ki – kidney; Sp – spleen; Int – intestines; H – heart; Lu – lungs.

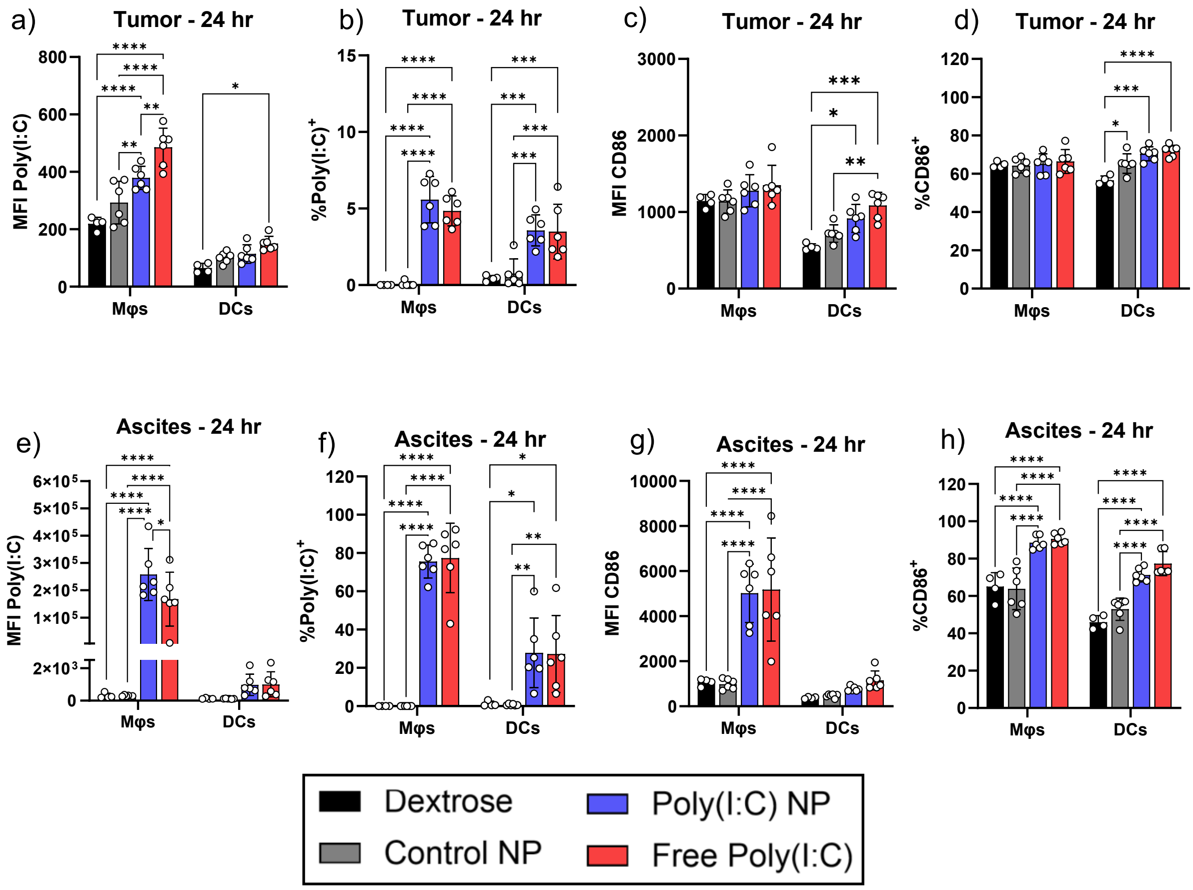


**Fig. S29. Cellular Poly(I:C) distribution in and activation of APCs in tumor and ascites 24 hr post-dosing.** Female C57BL/6 mice with BPPNM-luc tumors (3 million, IP) were injected with 50 µg poly(I:C) as free drug (Free Poly(I:C)) or encapsulated in LbL NP (Poly(I:C) NP). Ascites and tumors were collected 24 hrs after NP injection, digested, and subjected to flow cytometry analysis. Gating strategy described in **Fig. S31**. Median fluorescence intensity (MFI) of Cy5-poly(I:C) in antigen presenting cells (APCs)—macrophages (Mφs) and dendritic cells (DCs)—in a) tumor and e) ascites. Percentage of APCs that are Poly(I:C)$^+$ in the b) tumor and f) ascites. MFI of CD86 activation marker on APCs in c) tumor and g) ascites and the percentage of cells positive for CD86 in d) tumor and h) ascites. Data shown as mean±SD. Statistically significant comparisons determined using 2-way ANOVA Tukey multiple comparisons test.

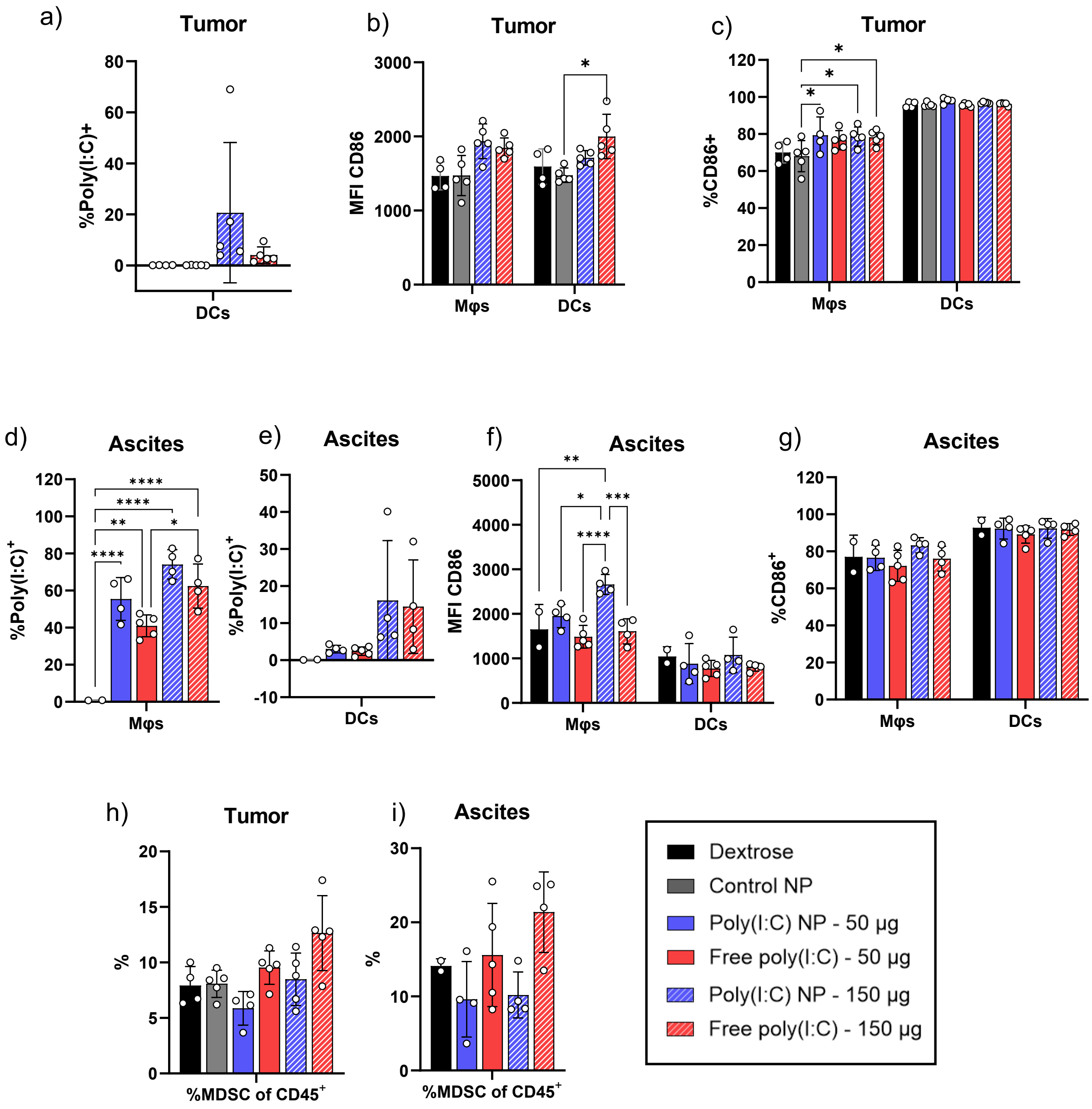


**Fig. S30. Cellular Poly(I:C) distribution in and activation of APCs in tumor and ascites 48 hr post-dosing.** Female C57BL/6 mice with BPPNM-luc tumors (3 million, IP) were injected with 50 or 150 µg poly(I:C) as free drug (Free Poly(I:C)) or encapsulated in LbL NP (Poly(I:C) NP). Ascites and tumors were collected 48 hrs after NP injection, digested, and subjected to flow cytometry analysis. Gating strategy described in **Fig. S31**. Percentage of APCs that are Poly(I:C)$^{+}$ in the a) tumor and d-e) ascites. MFI of CD86 activation marker on APCs in b) tumor and f) ascites and the percentage of cells positive for CD86 in c) tumor and g) ascites. Percentage of CD45$^{+}$ cells in the h) tumor and i) ascites that are myeloid-derived suppressor cells (MDSC). Data shown as mean±SD. Statistically significant comparisons determined using 2-way ANOVA Tukey multiple comparisons test.

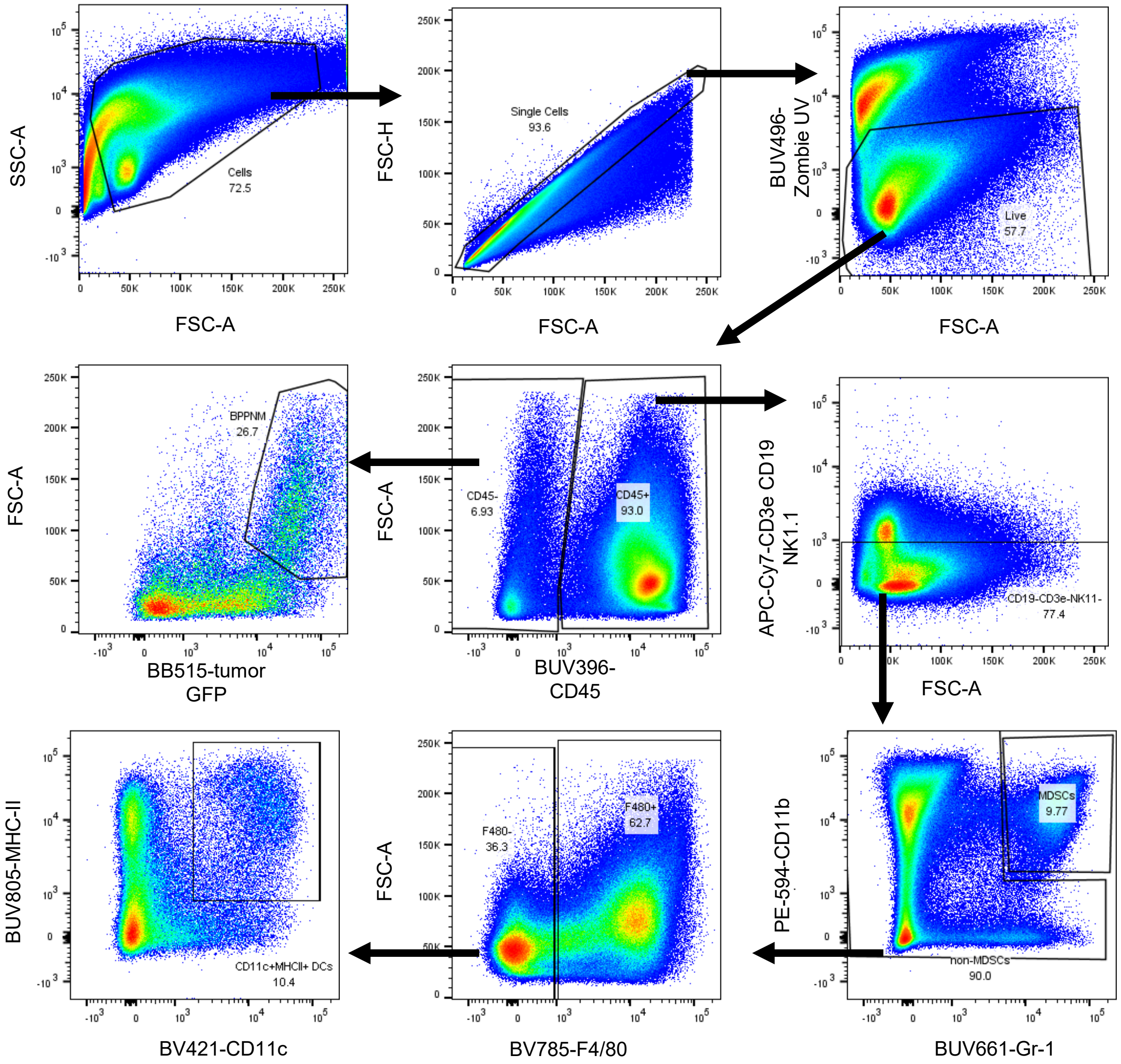


**Fig. S31. Gating strategy for cellular NP distribution in tumor and ascites.** Representative flow plots of cells from dissociated tumors gated for MDSCs (CD45$^+$CD11b$^+$Gr-1$^{hi}$), DCs (CD45$^+$F4/80$^-$CD11c$^+$MHC-II$^+$), macrophages (CD45$^+$F4/80+Gr-1$^{low}$), and BPPNM cells (CD45$^-$GFP$^+$FSC-A$^{hi}$). Gates set based on corresponding FMOs. The same gating strategy was applied to ascites cells.

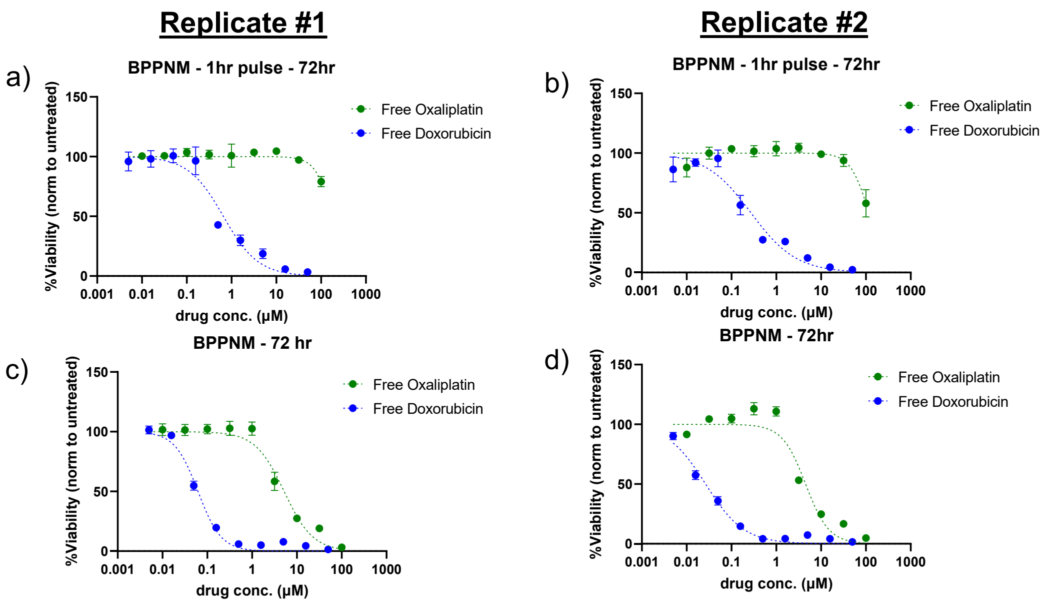


**Fig. S32. Sensitivity of BPPNM cells to free drug oxaliplatin and doxorubicin.** BPPNM-luc cells (2.5k/well) were seeded in black-walled, clear-bottom TC-treated 96 well plate and allowed to adhere overnight. Cells dosed with various concentrations of free drug in water (10% well volume). a-b) For 1hr pulse, the media was removed 1hr after dosing, and cells washed with 200 μL PBS, and fresh media added and cells cultured until 72hr timepoint at which viability was assessed. c-d) Cells were cultured with the drug for 72hr without any media replacement. At 72hr, the viability was assessed using PrestoBlue HS resazurin-based assay, relative to untreated cells. Data is presented as two independent biological replicates. (N = 3 technical replicates).

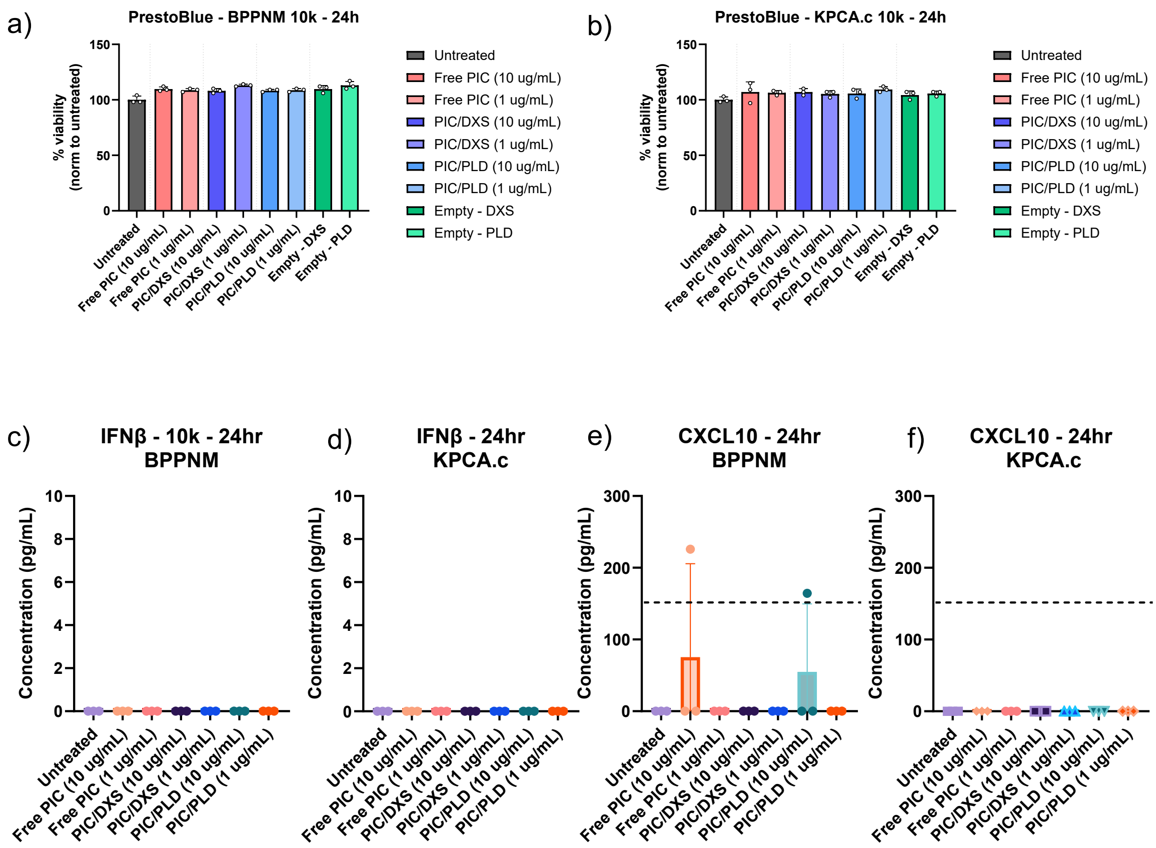


**Fig. S33. Ovarian cancer cellular response to LbL Poly(I:C) NPs formulated.** a) BPPNM or b) KPCA.C ovarian cancer cells were seeded in black-walled, clear bottom 96-well plates at 10k cells per well and allowed to adhere overnight. Cells were then treated with free poly(I:C) (PIC), LbL Poly(I:C) NPs with dextran sulfate (DXS) or the tumor-targeting layer poly-L-aspartate (PLD) as the terminal layer, or control bilayer NPs coated with DXS or PLD at 10 or 1 µg/mL poly(I:C) final concentration in the well or equivalent lipid dose. After 24hrs, a-b) cell viability was measured using PrestoBlue HS assay and c-d) IFNβ and e-f) CXCL10 measured using ELISA. Dotted line denotes limit of quantification (LOQ). Free poly(I:C) and LbL Poly(I:C) NPs did not have an impact in cell viability nor significant induction of inflammatory cytokines in these cell lines.

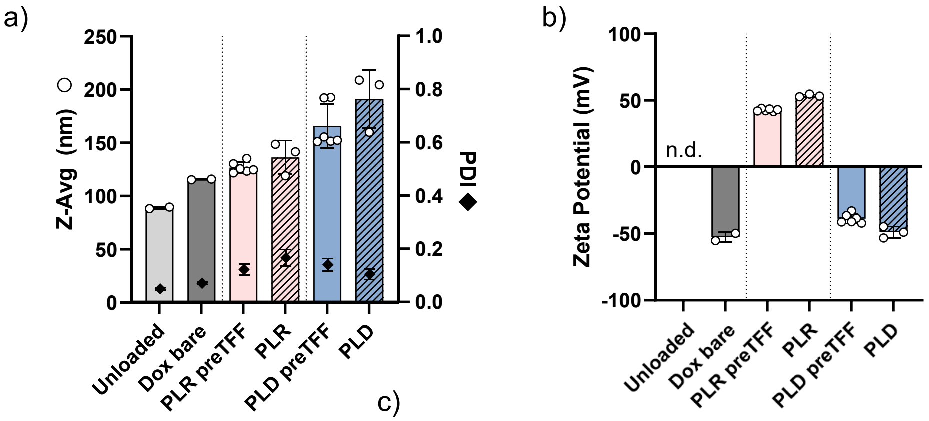


c)

| Layer | Wt% loading |
|---|---|
| PLR/PLD | 13.4 ± 0.8 |

**Fig. S34. Characterization of Doxorubicin-loaded LbL NPs,** a) Size, polydispersity index (PDI), b) zeta potential (ZP), and c) drug loading of doxorubicin loaded LbL NPs. Liposomes were formed via thin film hydration with ammonium acetate (unloaded), dialyzed against NaCl to remove unencapsulated ammonium acetate, then doxorubicin hydrochloride (Dox) was added to the liposomes to allow for active loading of Dox based on a pH gradient. Uncapsulated Dox was removed using dialysis (Dox bare) and then the liposomes were layered with poly-L-arginine (PLR) then poly-L-asparate (PLD) as the terminal anionic layer. NPs were purified using tangential flow filtration after each layer. Wt% loading is calculated as [mass_Dox / (mass_Dox + mass_carrier)] x 100%.

**Table S8. Layering conditions of LbL Doxorubicin NPs.** Anionic doxorubicin-loaded liposomes were layered with polymers at the weight (wt) ratio of liposomal core to polymer and final lipid and buffer concentrations listed. All LbL NP formulations were purified using tangential flow filtration to remove excess polymer and buffer exchanged into water after each layer.

| | Layering conditions | | |
|---|---|---|---|
| Layer | Wt ratio (core:polymer) | Final liposome concentration | Final buffer concentration |
| 1) PLR | 1:0.3 | 0.5 mg/mL | milliQ water |
| 2) PLD | 1:35 | 0.5 mg/mL | milliQ water |

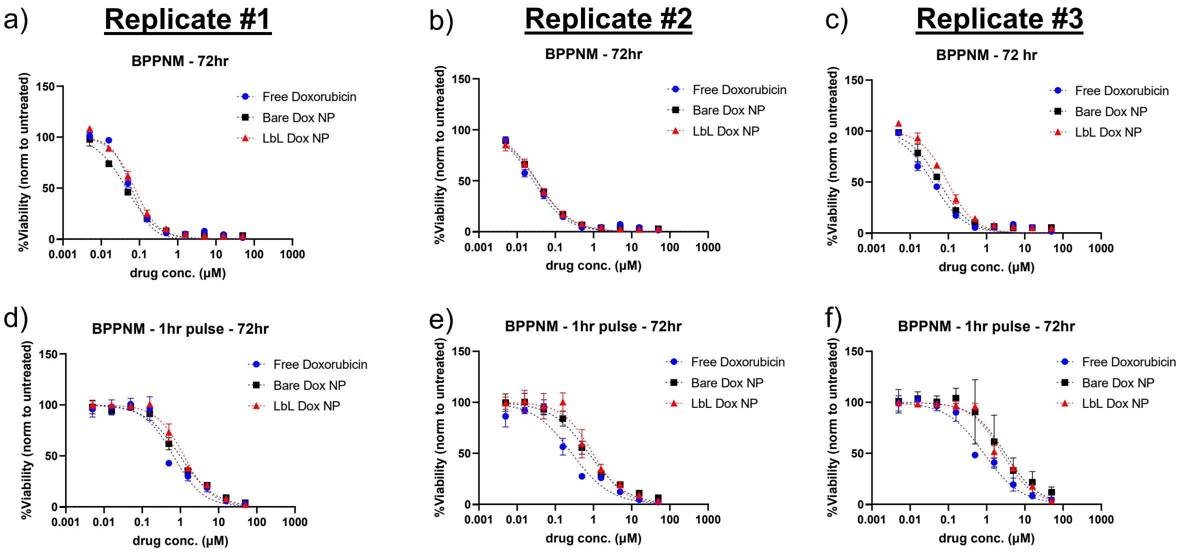


**Fig. S35. Sensitivity of BPPNM cells to free, liposomal, and LbL-NP doxorubicin.** BPPNM-luc cells (2.5k/well) were seeded in black-walled, clear-bottom TC-treated 96 well plate and allowed to adhere overnight. Cells dosed with various concentrations of Dox formulations (10% well volume). a-c) Cells were cultured with the drug for 72hr without any media replacement. d-f) For 1hr pulse, the media was removed 1hr after dosing, cells washed with 200 uL PBS, and fresh media added. At 72hr, the viability was assessed using PrestoBlue HS resazurin-based assay, relative to untreated cells. Data is presented as three independent biological replicates. (N = 3 technical replicates).

**Table. S9. $IC_{50}$ of free and NP formulations in BPPNM cells after 72 hr with or without 1 hr pulse treatment**. Data are shown as mean ± standard deviation of the three biological replicates in Fig. S35.

| $IC_{50}$ (μM) | Free Oxaliplatin | Free Doxorubicin | Bare Dox Liposome | LbL Dox Liposome |
|---|---|---|---|---|
| 72 hr | 4.85 ± 0.47 | 0.042 ± 0.018 | 0.045 ± 0.012 | 0.067 ± 0.032 |
| 1hr pulse – 72 hr | 149 ± 45 | 0.592 ± 0.293 | 1.59 ± 1.26 | 1.59 ± 0.86 |

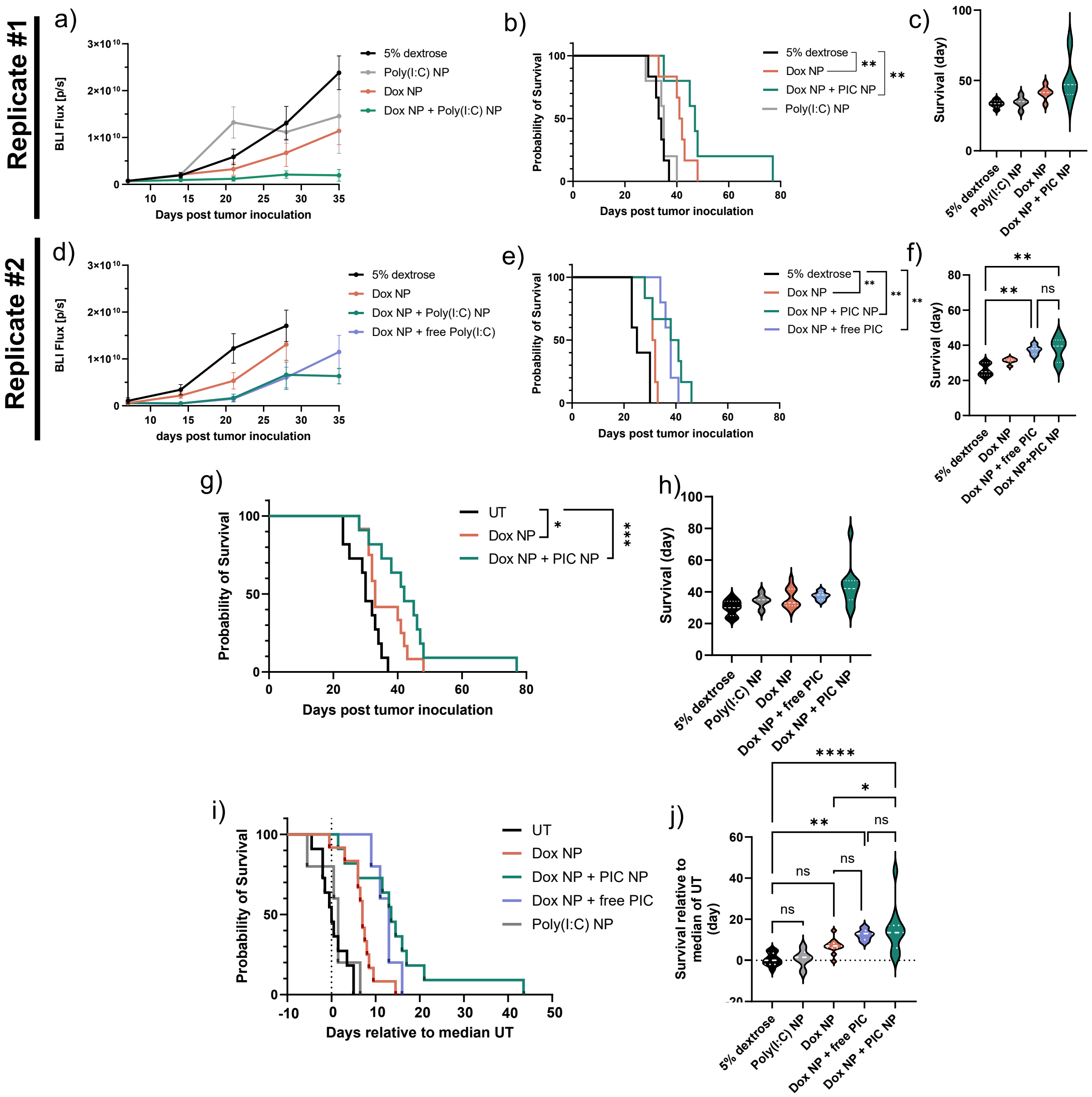


**Fig. S36. Survival analysis of BPPNM treated with Dox NP and Poly(I:C)**. BPPNM-tumor bearing mice were treated with Dox NP (2 mg/kg) and/or poly(I:C) (50 µg) as free drug or encapsulated in LbL NP. a,d) Tumor bioluminescence over time; b,e) Kaplan-Meier survival analysis, and c,f) Survival post tumor inoculation for both independent studies. g) Kaplan-Meier survival analysis and h) Violin plot of day of survival post tumor inoculation for both replicate studies combined without additional normalization. To account for differences in tumor growth rates between the two studies, survival data was normalized by plotting the survival relative to the median survival day of the Untreated group within each independent experiment: i) Kaplan-Meier survival analysis and j) Violin plot of day of survival relative to median survival day of Untreated group .

**Table. S10. Median survival for each experiment**

| | Median survival (days) | | | | |
|---|---|---|---|---|---|
| **Replicate** | **UT** | **Dox NP** | **Dox NP + Poly(I:C) NP** | **Dox NP + Free Poly(I:C)** | **Poly(I:C) NP** |
| #1 | 33.5 | 41.5 | 47 | — | 35 |
| #2 | 25 | 31.5 | 39.5 | 38 | — |
| Combined | 30 | 33 | 42 | | |

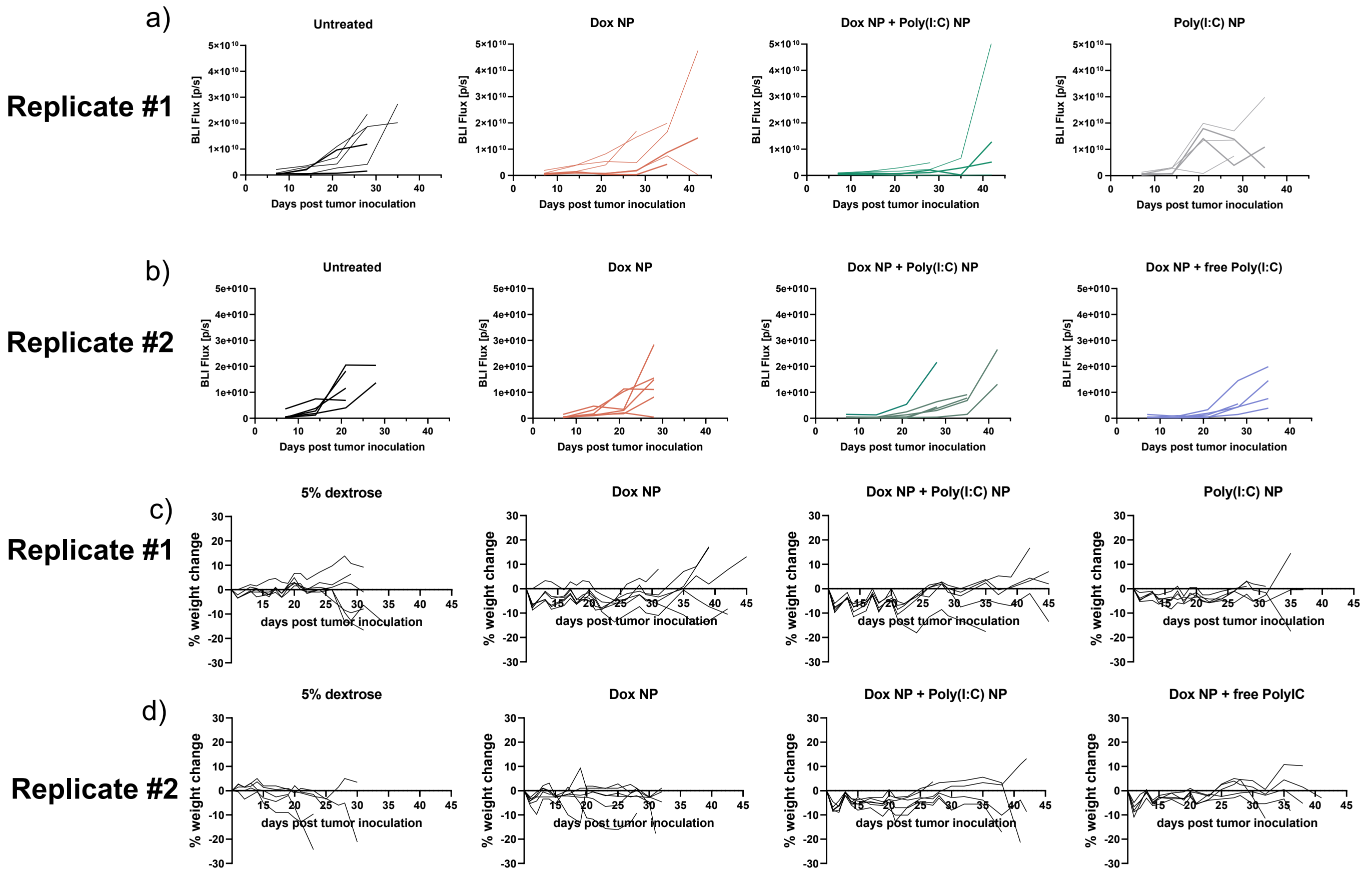


**Fig. S37. Individual tumor burden and weight changes of BPPNM mice treated with Dox NP and Poly(I:C)**. a-b) Tumor bioluminescence over time of each mouse over time for each replicate experiment. c-d) Change in weight from baseline for each mouse from both replicate studies.

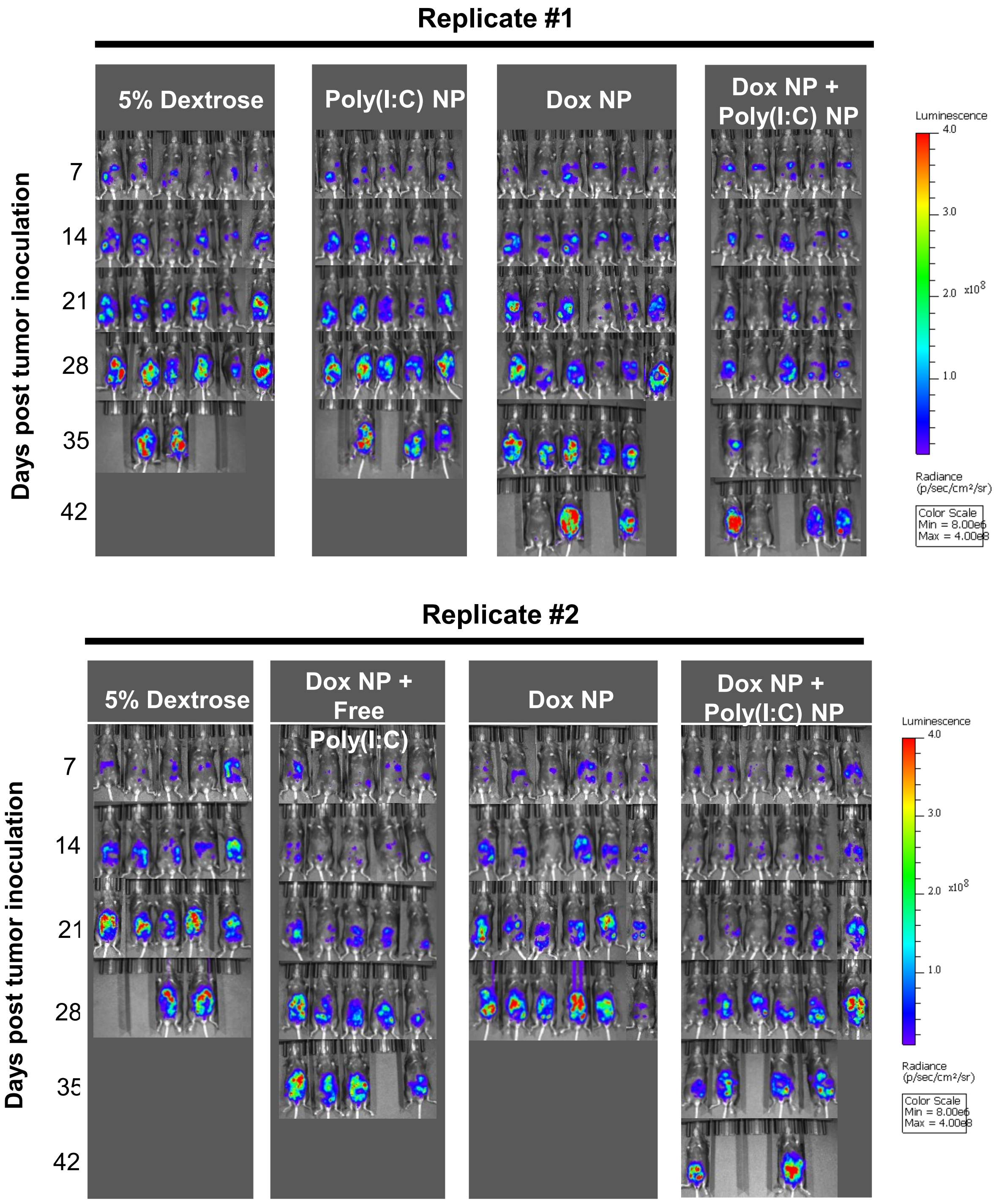


**Fig. S43. Tumor bioluminescence of BPPNM mice treated with Dox NP and Poly(I:C)**.
